# A signaling inspired synthetic toolkit for efficient production of tyrosine phosphorylated proteins

**DOI:** 10.1101/2024.12.22.629992

**Authors:** Margaret M. Ryan, Reagan Portelance, Graham F. Newman, Gabrielle Martinez, Swathi Shekharan, Anqi Wu, Savannah Angel, Katherine E. Schaberg, Petra Gilmore, Robert Sprung, Reid Townsend, Kristen M. Naegle

## Abstract

Tyrosine phosphorylation is an important post-translational modification that regulates many biochemical signaling networks in multicellular organisms. To date, 46,000 tyrosines have been observed in human proteins, but relatively little is known about the function and regulation of most of these sites. A major challenge has been producing recombinant phosphoproteins in order to test the effects of phosphorylation. Mutagenesis to acidic amino acids often fails to replicate the size and charge of a phosphorylated tyrosine residue and synthetic amino acid incorporation has high cost with relatively low yield. Here, we demonstrate an approach, inspired by how native tyrosine kinases find targets in cells – through a secondary targeting interaction, augmenting innate catalytic specificity of a tyrosine kinase, without overriding it. We engineered complementary vector systems for multiple approaches to producing high yields of phosphoprotein products in E. coli. Here, we test phosphorylation as a function of the targeting interaction (an SH3-polyproline sequence) affinity, different reaction methods across kinases of different specificity. This system presents an inexpensive and tractable system to producing phosphoproteins and phosphopeptides and we demonstrate how it can be used for testing antibody specificity on targets of EGFR and PD-1. This methodology is a generalizable approach for enhancing the enzymatic action on a recombinant protein via the flexibility of in vitro reactions and co-expression approaches. We refer to this as SISA-KiT, for Signaling Inspired Synthetically Augmented Kinase Toolkit.

## Introduction

There has been a massive expansion in the identification of tyrosine phosphorylation (1) across the proteome since its discovery (2) thanks to the development of enrichment methodologies coupled with mass spectrometry. More than 40,000 sites of tyrosine phosphorylation have now been identified in the human proteome (3). Tyrosine phosphorylation is a key signaling molecule that is tightly regulated and controls many of the biochemical networks that govern cellular processing of cytokines, growth factors, and other extracellular cues in eukaryotic cells. Despite its importance, our understanding of the function and regulation of these sites has not kept pace with their discovery – less than 5% of phosphorylation sites having a known kinase or function (4). One barrier to understanding the effects of phosphorylation on a protein’s fold, activity, or interactions is the difficulty around experimental testing. Phosphomimics by mutagenesis of a protein to a glutamic or aspartic acid fails to replicate the sheer charge and shape of a phosphotyrosine (pTyr). For example, phosphomimics fail to allow engagement of the necessary binding energy of SH2 domains (5), which is an important function for some unknown fraction of the phosphotyrosines in the human proteome. Therefore the most accessible of research tools is not tractable for understanding the large and highly charged version of tyrosine that is phosphorylated.

In addition to phosphomimics, two other technologies have been developed for the study of tyrosine phosphorylation. These are: 1) chemical ligation, the linkage of a synthesized phosphopeptide fragment to the N-terminus of a recombinant protein (6) and 2) incorporation of an amino acid analog using the amber codon and engineered synthetases. Unfortunately, chemical ligation is limited by the fidelity of phosphopeptide synthesis, meaning that only tyrosines that are within about 10 to 30 amino acids of the N-terminus can be studied by this method, a limitation for the types and locations of tyrosine phosphorylation in the human proteome. The use of synthetic analogs for pTyr have been hampered by the challenge of cell uptake, requiring a method to cover the negative charge. Pmp and pCMF are two analogs that have been created to mimic phosphotyrosine, which have had some success, although they do not exactly replicate size, charge, and pKa of a pTyr (7, 8). More recently, Hoppmann et al. (9) created a neutral phosphoramidate, which can be efficiently transported across the E. coli membrane. Although incorporation rates were high (90%), the process to deprotect the phosphate group requires incubation of the recombinant protein at pH 1.0 for 36 hours, which results in limitations for the study of natively folded proteins. Luo et al. (10), overcame the membrane transport issue by delivering Lys-pTyr or Lys-Pmp dipeptides (requiring phosphatase inhibitors). Unfortunately, the incorporation efficiency of the Luo strategy is poor (10%) and we noted that the authors only reported results for the Pmp analog in a biological case study, suggesting Lys-pTyr may have been insufficient for the study of pTyr function on proteins. The Pmp analog failed to show the same results as previously demonstrated by other studies, consistent with the pKa difference of the analog relative to pTyr (8, 10, 11). However, despite these challenges, chemical ligation and analogs have excellent site-specific incorporation of pTyr analogs and hold better promise for studying native phosphorylation than aspartic and glutamic acid mutants.

A final method researchers have used for studying tyrosine phosphorylation is to phosphorylate protein and with a purified, recombinant tyrosine kinase. A major challenge in this approach is the expense and complication of producing or purchasing tyrosine kinases. Additionally, our limited understanding of kinase-substrate relationships makes systematic exploration difficult. With roughly 90 tyrosine kinases in the human proteome, exhaustive exploration is prohibitive. Luckily, this barrier may be decreased thanks to the recent tour de force profiling of kinase specificity (12), which may enable fewer overall experiments from a set of likely candidates. However, we anticipate some deviations of kinase specificity for full protein targets since kinases invoke structural specificity and not just linear motif specificity (13). The expense of commercially available tyrosine kinases is often driven by their incompatibility with high-yield E. coli systems. Researchers have found that to produce tyrosine kinases in E. coli, they need to be co-expressed with phosphatases (14–16). However, this creates a competition between the kinase and phosphatase activity on a substrate and dephosphorylation of possible activation sites on the kinase itself.

We wished to develop an experimental approach that was easily usable by the majority of molecular biology labs with high efficiency of incorporation (avoiding the necessity for synthetic chemistry) and results in true tyrosine phosphorylation (overcoming the challenges of phosphomimic mutagenesis). Inspired by kinase biology, where kinase targeting to substrates is usually aided by additional spatial constraints through membrane localization or by additional protein interaction domains, we developed a synthetic kinase toolkit that targets kinases to a substrate of interest through an engineered SH3 domain interaction with small polyproline motifs. We developed several approaches, including an in vitro reaction, which does not require purification of the kinase or the addition of phosphatases, and a co-expression approach in E. coli. We demonstrate that although the targeting interaction can boost innate catalytic specificity, it does not over-ride it entirely, showing that the combination of phosphotyrosines incorporated on substrates with multiple tyrosines can be controlled. This toolkit represents a system that has less specificity than synthetic amino acid analogs, but is immediately usable by a wide array of research laboratories, modular, and effective with relatively high yields. The ability to vary the phosphoprotein patterns by selecting alternate targeting or kinase combinations will be useful for overcoming the challenges of producing complex phosphoprotein patterns on full proteins. Here, we demonstrate the principle of the toolkit on receptor tails, producing intact multisite phosphorylation of the epidermal growth factor receptor (EGFR) C-terminal tail and the PD-1 intracellular sequences, along with the LYN SH2 domain, and demonstrate how this approach can be used to validate phosphospecific antibody reagents and test the tradeoff between targeting affinity and catalytic specificity. Moving from complex substrates to simple oriented tyrosine peptides, we also demonstrate the ability to produce inexpensive phosphopeptide control reagents for antibody testing and other purposes. We refer to this as SISA-KiT, for Signaling Inspired Synthetically Augmented Kinase Toolkit.

## Results

### Toolkit design, components, and optimization

We sought to create two orthologous vectors that allow for the independent expression of kinase and substrate and the ability to purify recombinant, phosphorylated substrate protein at sufficient scale. We selected E. coli as the expression system for its high recombinant yield and lack of native tyrosine kinases. Our toolkit consists of two expression vectors – a substrate vector (based on a pGEX backbone) and a kinase vector (based on a pBAD backbone). We inserted an E. coli codon-optimized smt3 (yeast sumo) domain, given its long-standing demonstration as a solubility tag (17) on the N-terminal end of both vectors. For engineered targeting, we used an SH3 interaction domain interaction, as it interacts with a small peptide sequence with a defining characteristic of containing multiple proline residues (referred to as PxxP). Prior work by Pisabarro et al. (18) had established synthetic PxxP sequences with defined affinity with the ABL SH3 domain, varying from 0.4*µ*M to 34*µ*M, which we used as the basis of our targeting interactions. Hence, our toolkit consists of constitutively active kinases fused to the ABL SH3 domain and the substrate of interest fused to one of the PxxP sequences (Fig. 1A) or with no PxxP targeting sequence. We designed and tested in vitro- and co-expression-based reactions to evaluate the degree and type of phosphorylation as a function of targeting sequence (i.e. affinity), catalytic specificity (i.e. catalytic domain), and reaction types across various substrate targets. Throughout this work, we used a range of substrates. One of our substrates, the EGFR intracellular domain or C-terminal tail (Ctail), was selected due to the presence of nine physiologically important tyrosines, which have variable sequence motifs surrounding them (Fig. 1D). We used this as a primary model system in order to understand the tradeoff between catalytic specificity and the engineered secondary interaction.

**Fig 1.**
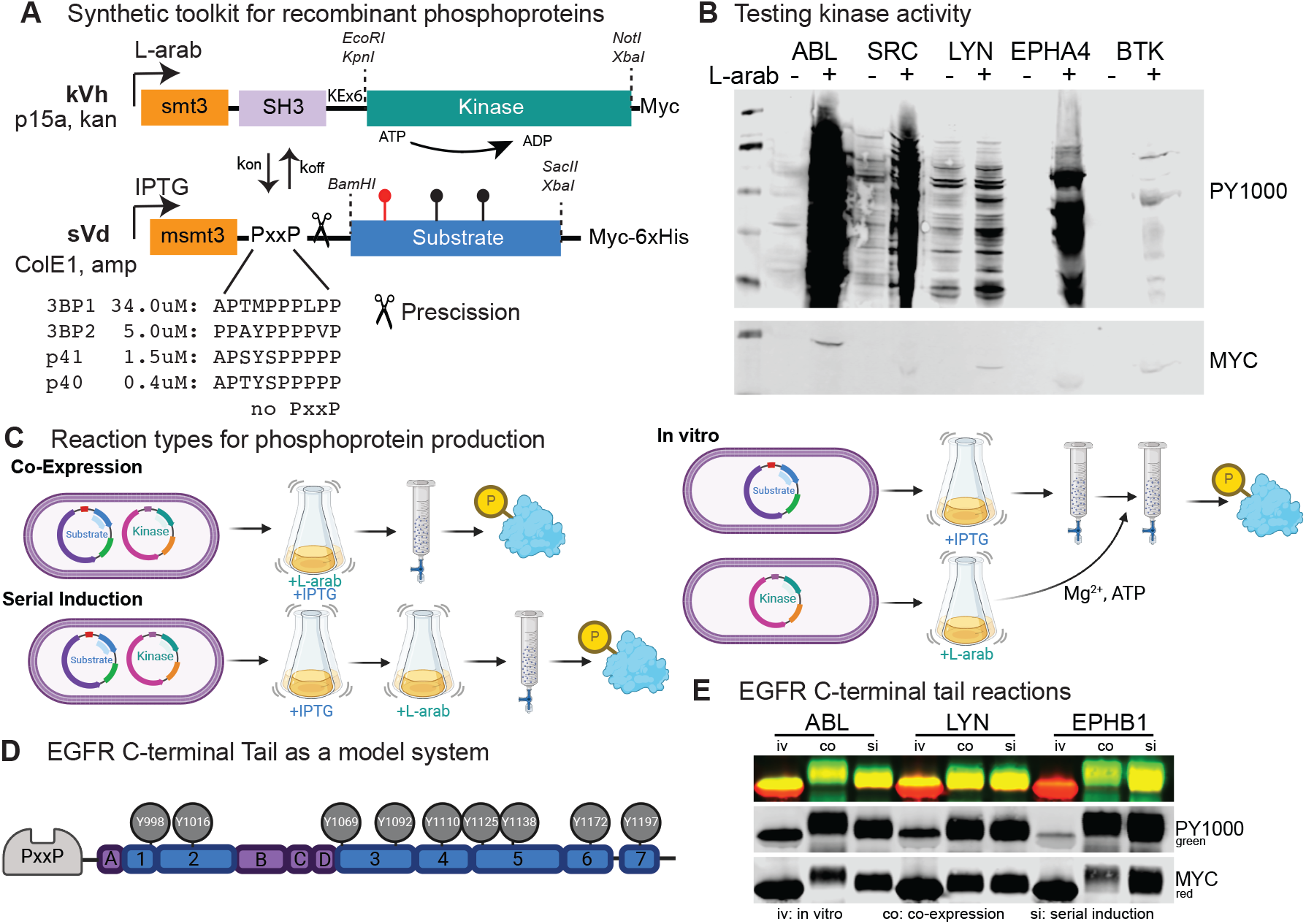
Overview of the synthetic toolkit for phosphoprotein production. A) We engineered orthogonal vector systems for production of an SH3-fused, constitutively active tyrosine kinase and a polyproline-fused substrate for recombinant production in E. coli. Origins of replication, resistance mechanisms, and other fusion tags along with vector cloning sites are indicated. The affinity of the engineered ABL SH3-polyproline interactions are from Pisabarro et al. (18). A PreScission protease site allows for the removal of the targeting sequence from the substrate and msmt3 refers to a (Y67F) mutated form of the yeast sumo domain. B) Example testing for constitutive tyrosine kinase activity by pan-specific phosphotyrosine antibodies (PY1000) and the comparative expression by MYC-based detection in E. coli without (-) and with (+) L-arabinose induction. First lane is a ladder. C) We developed three approaches to producing phosphoprotein. Co-expression involves co-transformations being simultaneously induced for substrate (IPTG) and kinase (L-arab) expression, before lysis and protein purification (generically indicated by the columns). Serial induction involves co-transformations first being induced for substrate (IPTG) and then switched to a kinase production phase (L-arab). In vitro reactions involve the purification of a single transformant induced for substrate and after binding and washes, the beads are resuspended in kinase-bearing E. coli lysate and supplemented with magnesium and ATP, before finishing protein purification. D) We selected the EGFR C-terminal tail (EGFRCtail) as one of our key substrates since seven tryptic fragments cover nine physiologically relevant tyrosine phosphorylation sites representing diverse sequences. E) Comparison of the reaction of three kinases (with p40 targeting) of the EGFRCtail, where the protein has been purified and cleaved using PreScission protease and probed with PY1000 and MYC antibodies.

#### Kinase Vector Engineering

We selected the pBad/Myc-6xHis vector system (L-arabinose induction) for expressing human kinases fused with the targeting SH3 domain, allowing for an independent induction of kinase, when in co-expression with substrates (on a Lac-repressor system). In order to allow for co-transformation and selection of dual-plasmid containing E. coli, we replaced the ampicillin resistance with kanamycin resistance and the origin of replication with p15A (to avoid competitive loss of replication with ColE1 origin of the substrate vector). Our first kinase construct insertion into the pBAD-smt3 backbone was ABL kinase with a TEV protease sequence on the C-terminal, allowing for removal of the epitope and purification tags (MYC, 6xHis). Subsequently, we replaced ABL (leaving the SH3 domain) with a new kinase, introducing a short linker (alternating glutamic acids and lysine for a flexible, but elongated linker), and multiple 5’ and 3’ restriction sites for easy cloning of new kinases. We intended to use TEV protease to remove the epitope tags to avoid cross-contamination with substrate purification. However, our attempts to cleave the Myc-6xHis epitopes on kinases failed, which we attribute to phosphorylation on the protease recognition sequence (ENLYFQ/S), which we detected by both western and by mass spectrometry on one of our library kinases (ABL). We tested for active kinase by probing E. coli lysates with pan-specific pTyr antibodies (e.g. PY1000, 4G10, or PY100). Although the L-arabinose expression of the pBAD system results in lower expression than a system like pGEX, especially with the lower copy origin p15a, we find that this expression is sufficient to drive phosphoprotein production, while maintaining E. coli health and avoids the need for additional phosphatases, typically required for high expression of active tyrosine kinases in E. coli (14–16). In fact, we find that maximal activity, as observed by the high phosphorylation of background E. coli proteins (using pan-specific pTyr antibodies), occurs for active kinases at relatively low amount of kinase according to MYC-based detection (Fig. S1). Hence, our final kinase vector is an L-arab-based expression system with the low copy number p15a origin, kanamycin selection, and includes convenient restriction sites for kinase cloning between an N-terminal smt3-SH3-linker and a C-terminal MYC detection tag. Some of our original kinases were cloned into this backbone which still carried a 6xHis C-terminal fusion (referred to as kVc kinases). We later removed the 6xHis tag (referred to as kVh kinases) to avoid competing with substrate purification that moved from GST to Nickel based purification.

#### Kinase library

To test the fundamental hypothesis of the toolkit, we wished to select tyrosine kinases from across a range of families, spanning different innate catalytic specificities. Since kinases are tightly regulated and E. coli may not produce functional forms of all human kinases, our selection was based on prior evidence of commercial products for recombinant tyrosine kinases – suggesting how they have been made as constitutively active products (such as through truncations of negative regulatory components). An initial panel for testing included ABL, SRC, LYN, EGFR, FAK, EPHA4, and BTK. Of these kinases, all but EGFR and FAK demonstrated activity (despite making the FAK truncation moving the regulatory FERM domain (19, 20)). We next made phosphomimic mutations of the activation loops of EGFR (Y869E) and FAK (triple mutant of Y570E, Y576E, Y577E). Activation loop mutations were not successful in activating these tyrosine kinases (Fig. S1), which highlights the limitation of phosphomimic mutations for tyrosines in replicating pTyr behavior in protein function. Additionally, we noted that on the five active kinases of our library, the kinases were themselves tyrosine phosphorylated, suggesting an important facet for activity is likely self-activation by phosphorylation (lacking in the EGFR and FAK kinases). Hence, we concluded that beyond testing of alternate forms of possible kinase boundaries, exhaustive exploration of the space for producing active kinase was not worth pursuing and instead we focused on expanding to alternate kinases. In later stages of the project, after establishing the methods of the toolkit approach, we expanded the tyrosine kinase library by subcloning all available tyrosine kinases from the open kinase library resource (Albanese et al (16)). Our current library consists of 17 tyrosine kinases, 15 constitutively active kinases, where 12 of them are highly dependable and consistent in their use of producing phosphoprotein. All kinases, their vectors and components, are detailed in Table 2.

#### Substrate Backbone

We selected the pGEX-6P-1 backbone for protein expression, a GST-fusion with a precision cleavage site before the protein insertion with a ColE1/pBR322 origin of replication, ampicillin resistance, and lac-based expression (we use the non-hydrolyzable IPTG analog for induction). We cloned the polyproline sequences (Fig. 1A) into the smt3-GST backbone between the GST and the PreScission sequence, yielding five substrate vectors (including the no targeting sequence version with the four targeting sequences). This configuration allows for the removal of targeting sequences from the substrates by protease to avoid targeting sequence effecting downstream functional testing. Our first substrate cloning into this backbone included the extension of a Myc and 6xHis sequence on the 3’ side to give us an alternate purification tag (if needed) and an antibody epitope sequence for detection. Similar to the kinase vector backbone, we engineered two 5’ and 3’ sites for easy cloning of substrate proteins into the set of 5 vectors, spanning different PxxP sequences. We refer to the final substrate backbone as sVd.

When we commenced testing phosphorylation of substrates with kinases, using an in vitro or co-expression approach as described below, we found the addition of targeting resulted in significantly increased phosphorylation of the protein fusion. However, following PreScission-based cleavage, we noted that the phosphorylation occurred predominantly on the N-terminal cleavage product (the smt3-GST-PxxP-PreScission fragment, Fig. S2), likely explained by the 14 tyrosine residues within GST. We anticipated that GST introduced several complications, including: 1) the possibility that GST-purification would be kinase-dependent, should phosphorylation effect purification, 2) screening phosphorylation of a target requires GST cleavage, increasing complexity and expense, and 3) the possibility that GST, with its abundance of possible kinase targets, could act as a competitor of the desired substrate phosphorylation. Since GST-based purification is typically a superior approach for yield and purity, we wished to retain the GST if possible and we synthesized a GST sequence with all tyrosines mutated to the non-phosphorylatable, but physiochemically similar phenylalanine (termed GSTm). Although we found that GSTm, had no discernible impact on production and solubility of the substrate protein, it also did not interact with the anti-GST matrix for protein purification (Fig. S3). Hence, we selected to delete GST from the substrate backbone instead, relying on the 6xHis for purification. Finally, in latter stages of this project, in order to eliminate non-substrate tyrosines, we mutated the single smt3 tyrosine to phenylalanine (Y67F, named msmt3) and a tyrosine in the construct to the N-terminal side of the smt3, and found this had no discernible effect on protein production or solubility (Fig. S4).

#### Optimizing a co-expression system

A challenge of independently inducing two proteins in a single E. coli strain is the complex coupling between metabolic carbon utilization, the strain genotype, and the repression systems used in the technologies chosen. Both lactose and arabinose are secondary carbon sources for E. coli and genes needed to enable the utilization of them first require the absence of preferred sources (glucose and mannose (21)), leading to increased cAMP and then activation of the global regulator CRP in a process referred to as carbon catabolic repression (22). E. coli strains have been engineered for particular repression systems, such as the 10*ω*, which was engineered for arabinose-based expression (23). In order to identify the E. coli strain that would enable both L-arab and IPTG-driven co-expression, we tested commercially available strains, TOP10 and BL21 DE3, along with a series of strains engineered for co-induction of both systems (MK01 (24), KL390 (CGSC#6207), and KL385 CGSC#6206). For these tests, we selected the LYN SH2 domain as a substrate, which represents a folded domain, where a prior study had shown that phosphorylation of LYN Y194 (within the SH2 domain) impacts binding of the SH2 domain (25), which we have found is conserved across a broad range of SH2 domains (26).

Across the five strains, we measured protein expression in single transformations and co-transformations, with and without inducer to evaluate leaky expression, control over expression, and total expression. Early in the project, we found that compared to TOP10, BL21 DE3 produced both higher yields of substrate protein and better control of kinase expression. However, we hypothesized that strains engineered specifically for both expression systems might produce higher total phosphoprotein, by increasing the relative kinase to substrate ratios, compared to the DE3 strain. We found that MK01 and KL390 appear to increase expression of kinase, at least based on relative levels of MYC-based detection, but all three strains (MK01, KL390, and KL385) produce low substrate protein and appear to lack ability to control substrate expression (i.e. all strains show basal leaky expression of substrate that does not increase upon coinduction, Fig. S5). However, to generate a true comparison, we purified protein from DE3, MK01, and KL385 (Fig. S6), which suggests that the rate limiting step in phosphoprotein production is producing large amounts of substrate protein. Therefore, we selected the BL21 DE3 strain as the major strain for production.

Finally, we tested the ability indole-3-acetic acid (IAA) (27) and cyclic AMP (cAMP) to improve expression by altering central metabolism or boosting a requisite cofactor, respectively. We found both factors had moderate effects on expression. In an IAA serial dilution curve (Fig. S7A), we observes slight increases in substrate expression with IAA around 1mM in the MK01 strain, which did not bear out in a change within the DE3 lineage (Fig. S7B). Within DE3, we observed a slight increase in kinase expression, as detected by MYC in the co-expression and single expression systems with the addition of cAMP. However, addition of cAMP did not appear to significantly change the phosphosubstrate production (Fig. S7B). Given the additional expense, without significant benefit, we used co-expression systems in LB media with no additional supplements beyond inducers and antibiotics.

#### Testing flexibility of expression and purification

In addition to co-induction of the kinase and substrate, we hypothesized that the secondary targeting interaction would enable an in vitro reaction, without the need to purify the kinase. We developed an in vitro reaction approach, where during the purification of substrate, after binding and initial washes, we resuspend the substrate-bound resin in an E. coli kinase lysate, supplemented with ATP and magnesium. Following kinase incubation, we proceed with washes and elution of the phosphoprotein. In vitro kinase reactions are capable of producing phosphoprotein, but at lower efficiency than the co-expression system (Fig. 1E). However, there are some advantages to the in vitro purification approach, which includes the ability to use alternate expression systems as needed for kinases that may not be active in the E. coli or to combine kinase specificities, such as co-expression of a substrate with one kinase and then continuing in an in vitro reaction with a second or a mix of additional kinases to complement specificities.

Additionally, we tested a “serial induction” approach, whereby we first produce substrate protein in the absence of kinase, avoiding modification of the protein during translation, and then induce kinase. For this, we developed a process by which substrate is induced overnight at 18°C and then the culture is shifted to 37°C, media is replaced, and kinase induction is started, stopping the culture 4 hours later. In a comparison of the p40-EGFRCtail production with three kinases (ABL, LYN, and EPHB1) by co-expression or serial induction, we found that the serial induction produced superior kinase expression, as seen by the MYC-based detection of the kinases (Fig. S8). This is likely due to the absence of the competing galactose analog during the kinase production phase. The time in the absence of substrate induction does not appear to lead to an appreciable loss of substrate, suggesting that serial induction is a helpful method for producing phosphoprotein. However, the overall production of phosphorylated EGFR Ctail by either co-expression or serial induction seems to be somewhat comparable, or even possibly superior in the co-expression, despite lower levels of kinase, as indicated by the upward shift of EGFR Ctail in the ABL co-expression result, which is likely due to increased time of kinase-substrate interaction. Although we did not identify a target by which folding and solubility was affected by kinase action during substrate translation, we anticipate that serial induction might be useful for such cases, and for increasing the phosphorylation of only surface accessible tyrosine sites. We used available EGFR phosphospecific antibodies to explore site-specific yields across in vitro, co- and serial-induction of p40-EGFRCtail with three different kinases (Fig. S9). Generally, we see high correlation between global phosphorylation (Fig. 1E) and site-specific readouts. For example, co-expression and serial induction generally produce higher phosphorylation than the in vitro reactions. The in vitro reactions yield interesting information about the preference of kinases for different sequences. For example, ABL appears to phosphorylate Y1016, Y1138, and Y1092 relatively strongly in the in vitro reactions, but not Y1069, Y1172, or Y1197. In the increased reaction lengths of co-expression and serial induction, ABL can phosphorylate Y1172 and Y1197, though it remains consistently low on Y1069. Despite having relatively low effectiveness in the in vitro reaction with EPHB1 across most of the sites, serial induction with EPHB1 produces very strong phosphorylation across all sites monitored by phosphospecific antibodies.

### EGFR Ctail phosphorylation under different reactions

To continue testing the fundamental hypothesis that we can augment innate catalytic specificity without overriding it, we used the EGFRCtail model system across kinases and targeting reactions. First, we wished to understand whether phosphorylation could be controlled by varying the level of affinity of the targeting reaction. For this, we co-transformed EGFR Ctail with LYN tyrosine kinase, where the EGFR substrate was fused to no targeting sequence or the four different PxxP sequences (Fig. 1) and monitored EGFR Ctail phosphorylation by a phosphospecific antibody for pY1173/pY1197 (we demonstrate later that the pY1173 antibody does not cross-react with the targeting sequence tyrosines, Fig. S12, though it may cross-react with EGFR Ctail sites beyond pY1173). Without induction of tyrosine kinase, as expected, there is no evidence of tyrosine phosphorylation. There is detectable levels of phosphorylation on EGFRCtail with co-expression of LYN kinase, even in the absence of targeting, which doubles upon adding a low affinity interaction (3BP1). The relative amount of phosphorylation doubles again with the next affinity interaction (3BP2), but appears to reach saturation (at least by western-based analysis) at higher targeting affinities. These results strongly suggest that adding an engineered secondary interaction between a kinase and a substrate can greatly enhance phosphoprotein production.

Next, to evaluate the complex degree of phosphorylation across the nine EGFR Ctail sites and to estimate the relative stoichiometry, we evaluated the EGFRCtail/LYN kinase reaction products by Phos-Tag analysis (28) (Fig. 2). Similar to the pY1173-based western analysis, we observe a significant increase in all forms of phosphoproducts as a result of targeting the kinase to the substrate via any one of the targeting sequences (Fig. 2B). Additionally, we observe that the dominant form of phosphoprotein produced on the Ctail by LYN kinase reactions results in a singly phosphorylated form, but we additionally see clear multisite phosphorylated forms that range from singly phosphorylated to many phosphorylated sites, with increasingly complex phosphorylation occurring with the higher affinity targeting sequences. Using the ratio of the MYC signal of the non-phosphorylated band to the area covering the various phosphoforms, we can estimate the approximate yield of phosphoprotein from this process, suggesting that the non-phosphorylated product drops from roughly half to one quarter when moving from non-targeted reactions to targeted reactions.

**Fig 2.**
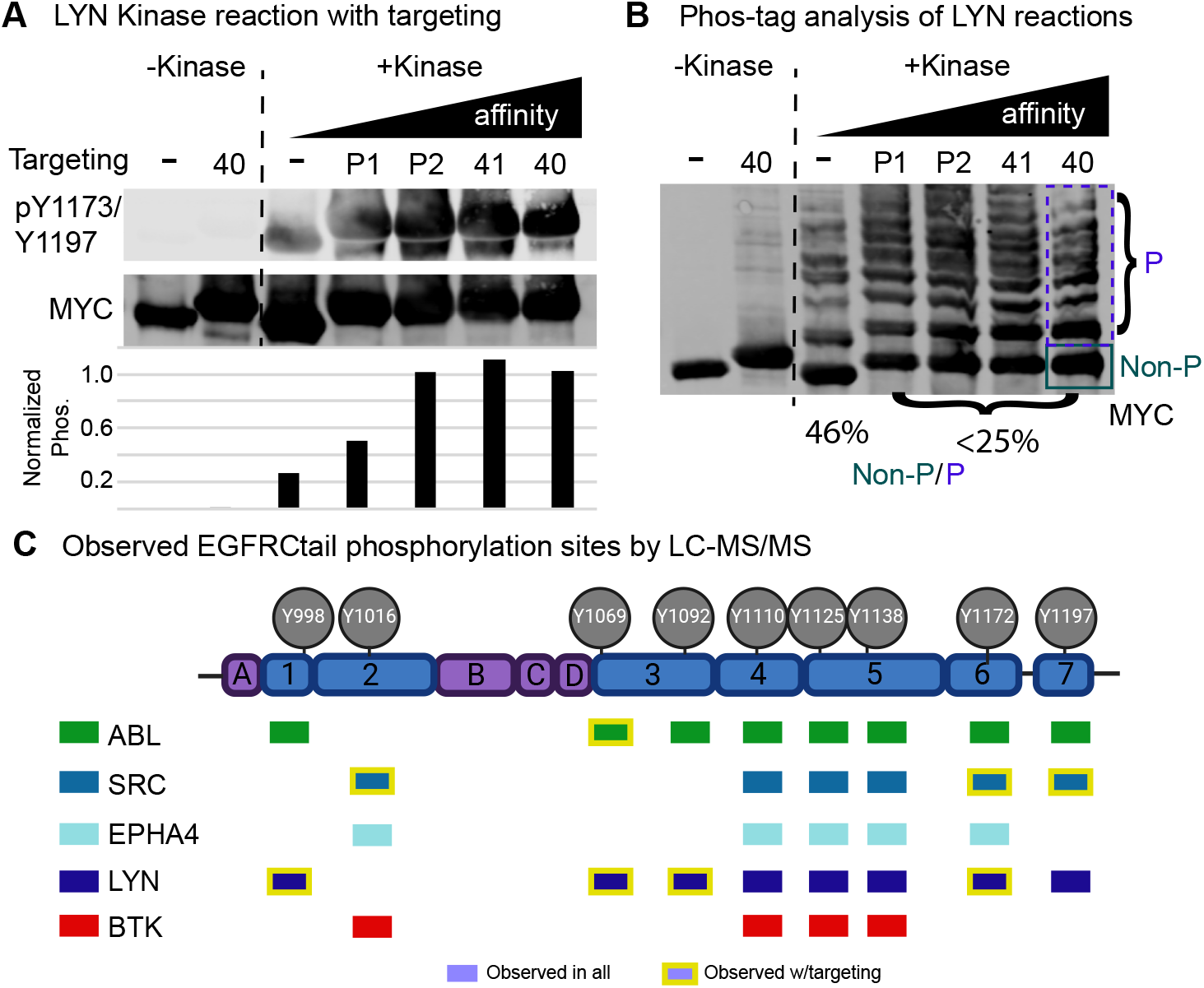
Evaluation of targeting affinity on phosphoprotein production of the EGFR Ctail construct. A) The sVd-EGFR Ctail was co-transformed with kVc-LYN kinase and purified from E. coli. EGFR phosphospecific antibody pY1173 (site Y1197 in the canonical isoform) was used to probe for changes in phosphorylation as a function of the assumed targeting affinity. Normalized signal is shown as the ratio of phosphosignal to MYC signal. B) The purified EGFRCtail samples were analyzed by Phos-tag analysis with anti-MYC antibody. The higher order forms were summarized by densitometry and divided by the non-phosphorylated band of the Phos-tag membrane to estimate the proportion of protein found as phosphorylated to some degree. C) We evaluated 24 samples of EGFRCtail by LC-MS/MS and identified whether high quality peaks were observed for the MS1 ion of the phosphorylated form of the tryptic fragment. A box indicates the phosphorylation site was observed across replicates and all conditions and a yellow outline indicates it was only seen under a targeting condition. Tryptic fragments A-D are non-tyrosine containing fragments that we used in testing SILAC ratios (Fig. S10).

#### Mass spectrometry-based analysis of EGFRCtail phosphopatterns

In order to evaluate the tradeoff between targeting affinity and kinase specificity in driving phosphorylation on specific sites, we used LC-MS/MS to evaluate phosphorylation on the EGFRCtail under different kinase reaction conditions using co-expression. We produced a SILAC standard spike-in, with the goal of using non-phosphorylated peptide forms of the light experimental phosphoprotein, compared to the SILAC standard, to estimate the degree of phosphorylation – attributing the loss of the non-phosphorylated form of the peptide fragment to the concomitant gain of the phosphorylated form. For the heavy SILAC EGFR Ctail standard, we adapted NMR-style protein production using a lysine/arginine auxotroph (29), yielding higher than 97% incorporation of the SILAC standard (a purity that is close to the theoretical maximum, due to the 1-2% of impurity in the heavy amino acids). In a preliminary study of 20 samples, spanning kinases and replications, we analyzed admixtures of the light phosphoproteins with the SILAC standard protein at a 5:1 ratio. We measured light:heavy protein ratios by quantifying tryptic fragments that lack tyrosines, representing “quantitative barcodes” that can be used to standardize protein content across different tyrosine kinase reactions. Of the four possible fragments seen across all samples, three of them gave highly reproducible ratios (higher than 0.997 correlation coefficients across all 20 samples). Ratios according to the three barcodes (Fig. S10) suggested that all of our samples were systematically lower in ratio than our target of 5:1, including some even lower than a 1:1 ratio. Given the high reproducibility, we hypothesized that our estimation of protein concentration by Coomassie comparison to a BSA standard curve may have systematic errors and those errors might be attributed to the high degree of phosphorylation in our samples. To test this, we treated samples with lambda phosphatase and compared Coomassie staining (Fig. S11), which demonstrated that Coomassie estimation is significantly affected by phosphorylation. Following this discovery, we treated sample with lambda phosphatase prior to protein concentration estimation and performed another large LC-MS/MS analysis on 26 samples, this time using a 25:1 sample to SILAC standard, following a back-titration experiment. This time, all sample ratios, as determined by each of the three barcode peptides, exceeded the SILAC standard by the target ratio. However, many were significantly higher than the target ratio, and we saw less concordance across the three different barcode peptides (Fig. S10B). When we tested normalization of the nonphosphorylated tyrosine-containing tryptic fragments to the global QBC standards, to estimate the percent remaining unphosphorylated, some ratios were above one and most ratios were inconsistent with western and Phos-tag based analysis, suggesting this SILAC normalizastion approach of remaining non-phosphorylated samples was not usable in this experiment.

We chose to focus on the binary analysis of whether a kinase phosphorylated a site and whether it changed with targeting. Focusing on the 26 sample experiment targeted by a 25:1 SILAC ratio, we evaluated MS1 peak quality for parent ions whose secondary fragmentation resulted in matches to the phosphorylated forms of tryptic fragments across the EGFRCtail (Fig. 2C). All no targeting and p40-targeting reactions were performed in duplicate and we required peaks be identified in both replicates in a condition to call it a site observed under a reaction condition. Replication was generally high, with 75.2% of sites being consistent between replicates in the no targeting conditions and 82% consistency within p40-targeting conditions (Table 1). Based on these criteria, some sites were only consistently observed under targeting, such as ABL phosphorylation of 1069, where other sites were phosphorylated by all kinases in all conditions (Y1110, Y1125, and Y1138). Globally, different patterns of phosphorylation occur across the Ctail, dependent on the kinase, where targeting generally increases the number of sites observed under a specific reaction condition. ABL kinase reactions with targeting leads to the most total observed phosphorylation sites, which is consistent with observations we made across the span of EGFRCtail experiments and might be explained by the processesive nature of ABL (30), where the relatively pleiotropic kinase specificity is typically defined by its binding domains. One complexity this highlights is that the ABL kinase, along with SRC and BTK families, used in this experiment included the SH2 domain, and multisite phosphorylation along the Ctail may create secondary targeting interactions through kinase SH2 domain recruitment. The EGFRCtail also has predicted ABL SH3 binding sites, according to Scansite (31). So although the binary analysis of the MS data of EGFRCtail reactions provides interesting insight into the patterns of phosphorylation and demonstrates that targeting generally increases phosphorylation, while still preserving some kinase specificity, it represents complex possibilities of multiple possible interactions of kinase binding domains, beyond the engineered SH3-polyproline interaction. Therefore, instead of continuing with more quantitative MS-based approaches to analyzing the EGFRCtail (such as using SILAC phosphopeptide spike-ins or optimizing the SILAC standard approach), we next chose to create a more refined experiment for testing the direct tradeoff between targeting affinity and kinase specificity for specific sites of the EGFR Ctail.

#### Phosphopeptide-based expression

To more directly test the hypothesis that we can augment innate specificity while retaining some specificity, we decided to clone specific EGFR tyrosine sites and the 10 amino acids directly surrounding the tyrosine on each side (referred to as 21-mers – each peptide is 21 amino acids with a central tyrosine). In order to make these small peptides large enough for purification and easy analysis by SDS-PAGE, we fused the mutated GST sequence (GSTm, where all tyrosines are mutated to phenylalanines) to the C-terminal side of our 21-mer (Fig. 3A). We hypothesized from earlier testing that GSTm would act as an inert and relatively large fusion domain that avoids introducing additional phosphorylatable tyrosines, without causing protein yield issues. We selected five EGFR tyrosine peptides, based on antibody availability and their pattern specificity seen in the EGFR C-tail mass spectrometry experiments (Y1016, Y1045, Y1092, Y1138, and Y1172). All but Y1138 had available phosphospecific antibodies. We generated a 21-mer control sequence, mutating the substrate 1069 tyrosine to phenylalanine to test for the phosphorylation of the targeting sequences containing tyrosines (3BP2, p41, and p40) and tested cross-reactivity with EGFR phosphospecific antibodies (Fig. 3B and S12A). Based on cross-reactivity we determined that when probing EGFR Y1069 and Y1138 with antibodies, the 21-mers need to be separated from targeting sequences by PreScission-based cleavage, whereas phosphospecific antibodies for Y1016, Y1092, and Y1172 can be used on full substrate product. We hypothesized that the homologous ERBB2 site antibody (ERBB2 Y1196) might cross-react with EGFR Y1138 and that this is a method that can help determine that. Interestingly, the noPxxP versions of many 21-mers appeared to have difficulty in expression or growth, with the 1092 21-mer being impossible to recover sufficient expression in the absence of targeting. However, we found sufficient expression of the remainder of the constructs to test the fundamental ability of a kinase to phosphorylate a specific site using co-expression.

**Fig 3.**
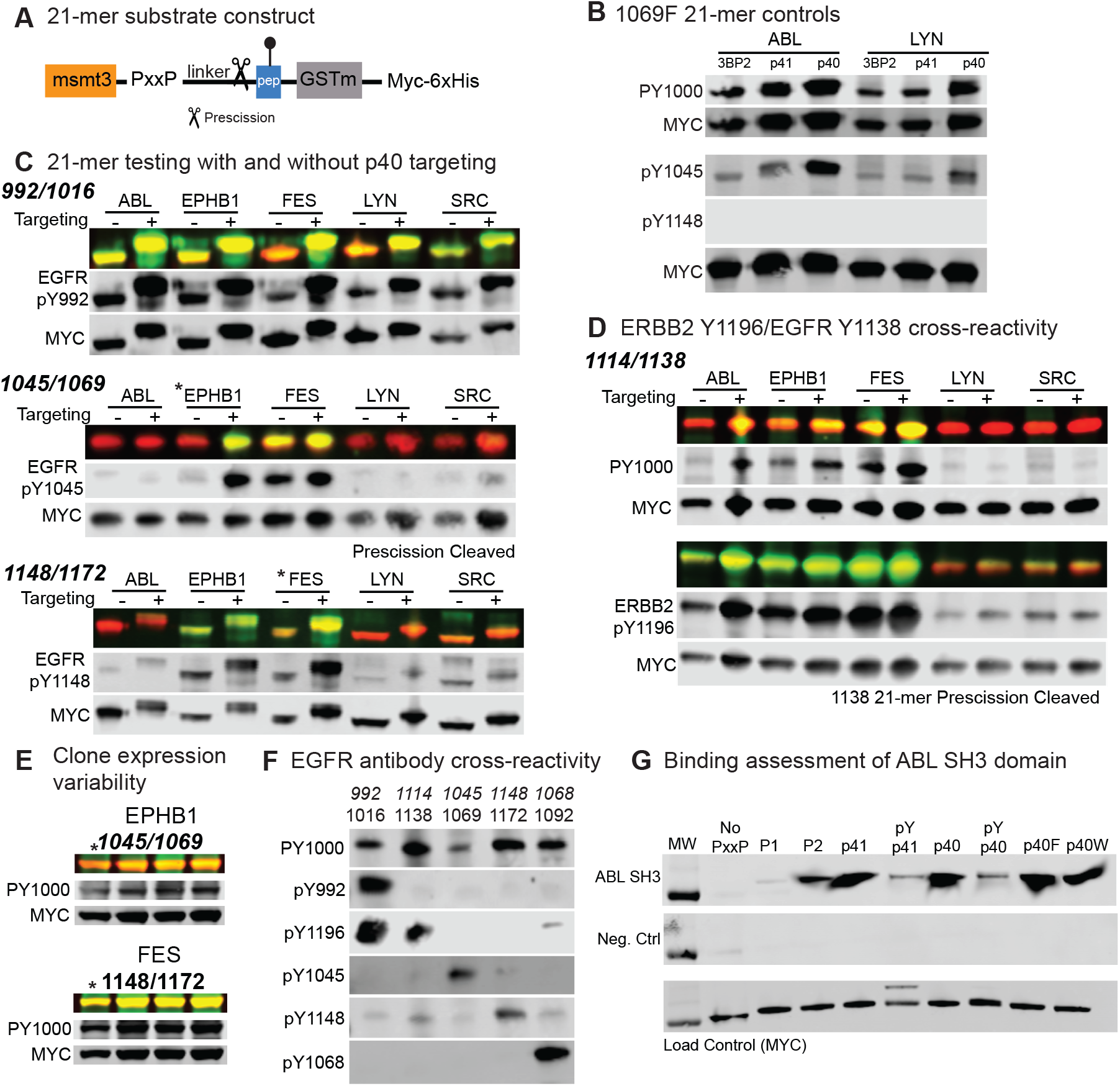
Construction and testing of specific tyrosine sites in a small peptide construct. A) Design of the substrate construct for small phosphopeptide production. GSTm is used as a large, inert handle for post-PreScission recovery of the peptide by SDS-PAGE. B) We cloned the 1069 EGFR 21-mer into each targeting construct and mutated the 1069 tyrosine (1069F), leaving just the targeting sequence tyrosine available for phosphorylation. ABL and LYN phosphorylate all targeting sequence tyrosines (as detected by pan-specific PY1000). This shows an example of an EGFR antibody (pY1045) which cross-reacts with the targeting pTyr sites and one that does not (pY1148) (all results in Fig. S12). C) The 21-mers were co-expressed with different kinases, with and without the p40 targeting sequence and PreScission cleaved, where necessary, for antibody specificity. MYC (red) and pTyr antibodies (green) are shown as combined in the top row, then separated in gray scale below. D) A 21-mer experiment with the 1114/1138 EGFR tyrosine site was used to test if the homologous ERBB2 site antibody is capable of detecting EGFR Y1138, where ERBB Y1196 appears to be more sensitive than the PY1000 to detecting EGFR Y1138 phosphorylation. E) We tested the variability of the EPHB1 and FES expression by inducing a replicate of the no PxxP glycerol stock (from panel D) and three independent clones of EPHB1/1069 and FES/1172 cotransformations. F) Having identified reactions that produce maximal phosphorylation of each EGFR site, we then tested cross-reactivity/specificity of the EGFR phosphospecific antibodies. G) We used a far western approach to measure relative binding of a biotinylated ABL SH3 domain with different targeting sequences, using the 1069F 21-mer construct. SH3 domain binding was detected by streptavidin-conjugated infrared dye. The negative control lacked the incubation with the SH3 domain and a separate load control western of the same samples is provided (Fig. S16 contains MYC and PY1000 confirmation).

Across the series of 21-mer peptide experiments co-expressed with five different kinases(Fig. 3), we observed the following general patterns: 1) phosphorylation of a site is predominantly determined by the kinase – i.e. some sites are well phosphorylated by some kinases but not all kinases – consistent with the concept that even in co-expression and with targeting the toolkit does not override the innate specificity of a kinase, 2) we generally observed targeting resulted in similar or sometimes more phosphorylation than the non-targeted condition – consistent with the concept that targeting likely augments innate specificity, and 3) some degree of kinase expression variability occurs, which can be evaluated by supernatant screening using pan-pTyr antibodies (Fig. S13). Generally saw the robust ability of kinases and substrates to express at sufficient level – of approximately 40 total experiments, only 3 resulted in low kinase expression. We compared the original clone to three new clones for two of these (EPHB1/noPxxP-1069 and FES/noPxxP 1172). Replication (Fig. 3E and Fig. S14) showed all replicates gave high expression of both kinase and substrate. This data suggests that FES phosphorylates 1148 without targeting and EPHB1 phosphorylates 1069 to a small degree, consistent with targeting enhancing 1069 phosphorylation by EPHB1. Additionally, it appears that when poor kinase expression has occurred, a replication of the induction or selection of a new clone is likely sufficient to boost yield.

In addition to using phosphopeptides to test kinase specificity and reaction conditions for specific tyrosine sites, the phosphopeptides are also directly usable as products. Here, we used the phosphopeptides to optimize antibody probing conditions (Fig. S15), which suggests that including a synthetic 21-mer positive control on a western could be helpful when probing human cell lysates with phosphospecific antibodies. Additionally, the production of a phosphorylated EGFR Y1114/Y1138, allowed us to test ERBB2 pY1196 antibody as a cross-reaction with EGFR pY1114 (Fig. 3D), which showed excellent binding characteristics. Expanding on this, we took conditions of high phosphorylation across each of our 21-mers and tested the specificity/cross-reactivity of each phosphospecific antibody with other EGFR sites in our 21-mer constructs(Fig. 3F). We found that ERBB2 pY1196, in addition to binding EGFR pY1138, binds strongly to EGFR pY1016, and to some degree pY1092. Most of the antibodies were fairly specific (especially pY1068), with the exception of pY1148, which though it bound most strongly with its target, had noticeable off target binding with most of the EGFR phosphopeptides.

#### Targeting sequence recruitment

Although we generally observed an increase in the phosphopeptide experiments with targeting, we observed less of an augmentation than anticipated, compared to other experiments (such as in the full EGFRCtail experiments Fig. 2, or even phosphorylation of GST in our original substrate constructs, present only with targeting (Fig. S2). There are a few possibilities for the muted targeting results. First, on larger substrate constructs, which might contain additional recruitment sites (either for SH2 or the SH3 domain) the processive nature of kinases amplified initial recruitment. Another possibility is that the linkers we engineered in the peptide construct may not be ideally matched to the various kinase construct separation between the SH3 domain and the catalytic domain, which would be detrimental for these small constructs. Finally, three of our targeting sequences contain tyrosines, which may themselves be subject to phosphorylation by a kinase in a co-expression. To test if our high affinity recruitment interactions are affected by phosphorylation, we performed a binding assay and included two non-phosphorylatable targeting sequences (p40W and p40F, where the p40 tyrosine is mutated to tryptophan or phenylalanine) as candidates. Using a far western approach and the control 1069F peptide, which allows us to present the targeting sequence in phosphorylated and unphosphorylated forms, we tested relative binding affinity with the ABL SH3 domain (Fig.3G – using biotinylation of a C-terminal AviTag (32) sequence and a streptavidin-conjugated dye for detection). The qualitative binding of the ABL SH3 domain to the target sequences matches very well to the Pisabarro determined affinities (18) (Fig. 1A). Interestingly, we observe a loss in binding of the p41 and p40 sequences when tyrosine phosphorylated, placing the phosphorylated forms of these high affinity sequences closer to the effective binding affinity of the 3BP1 targeting sequence. Based on binding, the p40W and p40F also appear to be relatively high affinity interaction sequences, which present possible sequences that lack the additional complexity of being modified by kinases within the toolkit. Hence, we conclude that while the phospho-p40 sequence still contributes a targeting interaction, its affinity is significantly lower than the unphosphorylated p40 sequence, which creates a complication that might be kinase specific, depending on the degree to which a particular kinase phosphorylates it and the rate it which it phosphorylates it (relative to the SH3 domain interaction). Finally, we should note that these binding data are with an unmodified SH3 domain and we cannot discount the possibility that the SH3 domains fused to kinases might themselves undergo phosphorylation, that impacts the binding affinity with engineered target sequences (an effect that is known to be physiologically relevant (33)).

### Application to PD-1 phosphorylation and antibody testing

Across the span of toolkit development, we evaluated the capabilities of producing complex phosphopatterns on the full EGFRCtail, targeted peptides, along with some degree of phosphorylation on folded domains, such as the LYN SH2 domain. We also learned a great deal about the complexity of engineering interactions, variability in expression, and methods for increasing yields. We wished to test this culmination of work in a final application that might be immediately relevant to important biological applications and selected the PD-1 C-terminal tail. PD-1 has two tyrosine sequences, the ITIM and ITSM sequences, which act to recruit key downstream regulators and helps to determine how PD-1 as a cofactor will reshape activation of Tcells, a process known to be at work in hiding cancer cells from the immune system (34). There is still a lack of fundamental knowledge regarding PD-1 and the factors that fully describe whether it will ultimately stop or not activation cascades and hence a reagent for testing antibody reagents and binding would be highly useful.

We cloned the PD-1 intracellular domain, spanning the last 95 amino acids (covering the ITIM and ITSM tyrosines, which are the only tyrosines in the construct) into a substrate construct carrying the p40W targeting sequence. We included an N-terminal ALFA-tag (a non-tyrosine containing epitope that allows for additional detection (35)) and a C-terminal AviTag (32) for biotinylation. We produced single tyrosine to phenylalanine mutants for each tyrosine (Y223F and Y248F, as well as a “FF” control, where both tyrosines are mutated to phenylalanines). ABL performed well, phosphorylating both individual sites (Fig. 4A). We purchased Abcam’s PD1 pY248 antibody, but failed to see binding of that antibody to our Y248 phosphorylated proteins. We tested this across two batches of antibody, with identical results. We then purchased the BioXcell PD-1 pY248 antibody and observed that this binds the phosphorylated PD-1 wild type construct and the Y223F, suggesting that it recognizes pY248 and does not cross-react with the pY223 site. These results demonstrate the power of producing inexpensive controls using this synthetic toolkit – highlighting the possibility that the Abcam antibody in publications may be detecting something other than PD-1 (or that both batches of antibody simply did not work in our hands across replications) and that the BioXcell antibody can be used for the purposes as described, including not cross-reacting with the second PD-1 site.

**Fig 4.**
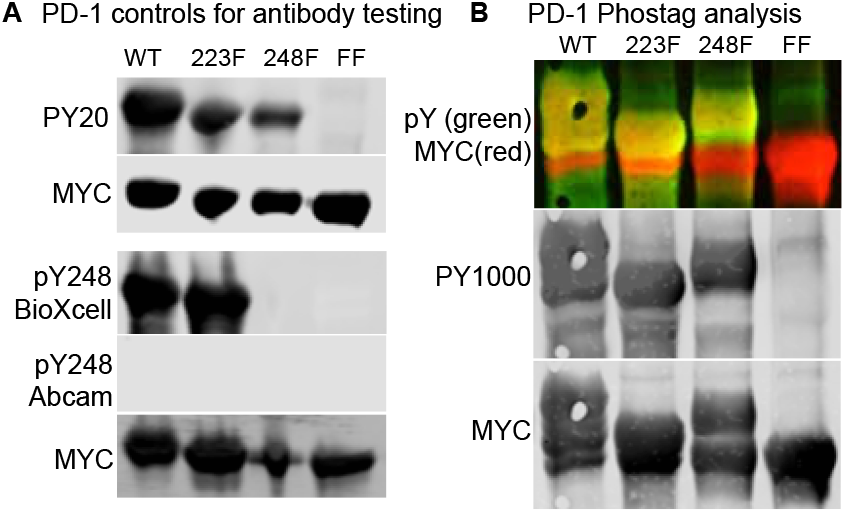
Extending the toolkit to produce PD-1 ITIM and ITSM phosphorylation sites. A) We produced WT PD-1 as a substrate in the toolkit, including the single and double F control mutations (223F and 248F). Using ABL kinase, and pan-specific pTyr we found phosphorylation of both sites (here PY20 is shown) and that only one of the PD-1 pY248 antibodies recognized 248, and it did so specifically. B) Here, a Phos-tag analysis shows the degree of phosphorylation that is achieved, demonstrating high degrees of phosphorylation on one and both sites.

Next, we wished to estimate the degree of phosphorylated forms of PD-1 we were capable of producing under this reaction and used Phos-Tag analysis. Interestingly, Phos-Tag separated the pY223 and pY248 to slightly different degrees, enabling the evaluation of individual site and double site phosphorylation in the wild type construct. The FF control highlights the complete lack of phosphorylation on this construct and the migration size of the unphosphorylated form, to which the unphosphorylated forms of the tyrosine-containing constructs can be compared. We observe that each individual site in this experiment was phosphorylated at about a 50% yield. The majority of the wild type protein is phosphorylated to some degree (approximately 75%) – with the appearance of higher molecular weights, consistent with some degree of double phosphorylation (though full resolution of this band is not possible).

## Discussion

This flexible toolkit, based on the idea of targeting kinases to substrates, enables the efficient production of phosphoproteins and phosphopeptides. Across the wide array of substrates and reaction conditions, the data strongly suggest that targeting augments innate specificity without overriding it. As this project progressed, we discovered an expanding range of protein products that could be produced, from simple phosphopeptides to complex multisite phosphorylated proteins. The ability to produce multisite phosphorylation on disordered proteins, such as receptor tails, could be highly useful for the study of complex molecular assembly that might occur at the proximal layer of intracellular phosphotyrosine signaling. In addition to disordered proteins, such as EGFRCtail and PD-1, we also showed more complex domains can be phosphorylated, such as the LYN SH2 domain. In addition to products that can be used to understand the effect of phosphorylation on proteins, the ability to produce folded phosphoproteins might prove to be highly beneficial for the purposes of generating antibodies for in situ applications – overcoming some of the limitations of antibodies raised to linear phosphopeptides that relegate many of them to western-based approaches.

Although this toolkit achieved the key aim of creating a tractable molecular biology approach for producing phosphoproteins, with higher fidelity in reproducing true tyrosine phosphorylation, we also ran into challenges. For example, the complexity of tyrosine phosphorylation on targeting sequences, and the possibility of SH3 domain phosphorylation results in a lack of clarity in how and when effective targeting strategies are at play. Future work could include the engineering of a targeting domain that is unaffected by possible tyrosine phosphorylation. We had also anticipated that there could be an affinity that is too strong – one that reduces the effective catalytic turnover of the kinase by having too high of a secondary interaction affinity, lowering the diffusion of the kinase and possibly introducing steric hinderance for other substrates. We did not see this possibility play out with the high affinity p40 sequence. However, given the phosphorylation of p40 reduces affinity with the SH3 domain, it is possible that our experiments never explored a truly high affinity secondary interaction. Finally, as we tested across a range of substrates, the role of additional domains, donated by native kinases, such as SH2 domains from SRC/ABL/BTK families, created additional processivity for multisite phosphorylation – an affect that can be beneficial when seeking to make multisite phosphorylated products, but which might also be undesirable, depending on the application. Despite lessons learned through experimentation, which can guide future toolkit designs enabling tighter and more predictive control over target production, the current toolkit is already highly efficient and usable as a tool for a range of molecular biology applications.

Some of the more surprising results of this work was the lack of need to drive higher kinase production than what was possible under the low copy number plasmid in the non-optimized BL21 DE3 strain. It was interesting that across strains, engineered for better control of the independent operon systems, that we ultimately found substrate production was the rate limiting step and that driving higher kinase activity had minimal gains on phosphoprotein yield. We were able to identify flexible methods within the DE3 system, such as serial induction, that resulted in higher overall kinase expression. However, the gains made by the kinase production had less obvious impacts on overall phosphoprotein production, though it remains a tool that can be used for increasing toolkit flexibility. Additionally, the flexibility of adding in vitro kinase reactions, where efficiency gains are made by the addition of the secondary interaction, opens a number of possible extensions (such as production of kinases in alternate recombinant systems if necessary for activity). Finally, we anticipate that this general approach – engineering a secondary interaction between a substrate and an enzyme and the controlled interaction of them in recombinant expression or by in vitro reactions – can be extended well beyond tyrosine phosphorylation and include enzymatic action on proteins of other post-translational systems or other types of chemical or biochemical modification.

## Materials and Methods

Reagents, part numbers, and vendors have been provided in Table 3, including antibodies, species, and dilutions used in western-based applications. All kinases, substrates, their DNA and protein coding sequences, along with their internal plasmid identifier, the regions they span in their human targets are provided in Table 2 and their Snapgene DNA files, including sequencing results, are included as supplementary information.

### E Coli strains and competency

BL21-Gold (DE3) (Agilent), TOP10 (Invitrogen), and DH5alpha for subcloning (ThermoFisher) strains were purchased as chemically competent. Strains MK01 were gifted from Dr. Manjunatha Kogenaru and Dr. Sander Tans by mail, as was the double auxotroph strain by Dr. Ron. T Hay (29). Strains KL390 (CGSC#6207), and KL385 (CGSC#6206) were purchased from the E. coli Genetic Resource Center (ECGRC), named Coli Genetic Stock Center (CGSC) at the time. Non-commercial strains were made chemically competent for transformation following the procedure by Chung and Miller (36).

### Cloning and mutagenesis

We used either restriction/ligation cloning or In-Fusion cloning for engineering of backbone vectors, insertion of tags and targeting, and all kinase and substrate insertions. We used Snapgene to design primers, and sequenced verified by Sanger sequencing using either GeneWiz or Eurofins. Sequencing was performed to get full coverage of insert. Some constructs were whole plasma sequenced to verify the entire plasmid. We used Agilent’s QuickChange mutagenesis kit to perform point mutations. The full GSTm (all tyrosines mutated to phenylalanines) was synthesized by GeneWiz and sent as a pUC57 backbone insert, which we then cloned and sequence verified into desired substrate backbones. We used carbenicillin at 100*µ*g per mL for selection of ampicillin plasmids and kanamycin at 50*µ*g per mL for selection of kanamycin plasmids.

### Plasmids

Full plasmid sources are provided in Table 2 and Snapgene DNA files containing full plasmid details of generated kinases are available at https://doi.org/10.6084/m9.figshare.28299335 and substrates are available at https://doi.org/10.6084/m9.figshare.28296383. pDONR223-LYN was a gift from William Hahn & David Root (Addgene plasmid # 23905; http://n2t.net/addgene:23905; RRID:Addgene_23905) (37); pDONR223-BTK was a gift from William Hahn & David Root (Addgene plasmid # 23918; http://n2t.net/addgene:23918; RRID:Addgene_23918) (37); pDONR223-EPHA4 was a gift from William Hahn & David Root (Addgene plasmid # 23919; http://n2t.net/addgene:23919; RRID:Addgene_23919) (37); pDONR223-EPHA4 was a gift from William Hahn & David Root (Addgene plasmid # 23919; http://n2t.net/addgene:23919; RRID:Addgene_23919) (37); pSG5-ABL was a gift from Nora Heisterkamp (Addgene plasmid # 31284; http://n2t.net/addgene:31284; RRID:Addgene_31284) (38); pcDNA3 c-SRC (WT) was a gift from Robert Lefkowitz (Addgene plasmid # 42202; http://n2t.net/addgene:42202; RRID:Addgene_42202) (39); pBi-Tet-EGFR-Nluc was a gift from Linda Pike (40); pCMV3-ALFA-PD1 was a gift from Diane Lidke.

### Protein induction and bacterial lysis

An isolated clone was used to inoculate 5mL LB media with appropriate antibiotics in a 15mL culture tube and incubated at 37°C overnight while shaking. For each 11mL of overall culture, we split the growth into two phases. In the first phase, we inoculated 5 mL of LB media supplemented with antibiotics (antibiotics were halved, compared to antibiotic concentration used for selection as noted in cloning) with 150 *µ*L overnight culture, incubating for 2.5 hours at 37°C. We then supplemented with 6mL additional LB media with antibiotics, growing cultures to an OD600 of 0.7 (log phase growth). We then induced with 0.5mM IPTG and/or 0.2% L-arabinose, growing at 37°C for four hours or overnight at 18°C. Bacteria were then pelleted and media was decanted and either directly lysed or frozen at −20°C and then lysed at a later date. We resuspended pellets in a 10:1 ratio of final growth volume to lysis buffer. Lysis buffer was 50mM Tris-HCl at pH7.0 and 150mM NaCl, supplemented with protease inhibitors, phosphatase inhibitors, and PMSF (1:1000). For small volume lysis of 1mL to 5mL, we used bead beating, adding 400*µ*L of beads to each 1mL of resuspended cell pellet and bead beat for 4 minutes. For larger lysis volumes, we sonicated resuspended pellets for 5 minutes with 40% pulse sequence. Following lysis, we frequently sampled and evaluated the raw lysate, prior to clarification by centrifugation at 12,000xg for 10 minutes at 4°C.

### Serial induction

Serial induction involves the same initial inoculation and induction phases with 0.5mM of IPTG to induce the substrate overnight at 18°C. Following this, the bacteria is pelleted, media is decanted, and then the pellet is resupsended in fresh LB, supplemented with antibiotics and 0.2% L-arabinose. Kinase induction by L-arabinose is performed for 4 hours at 37°C, then pelleting and lysing proceeds as described above.

### Nickel/6xHis Purification

We used Genscript Ni-NTA MagBeads, using a magnetic tube stand for isolating beads during decanting. Prior to protein binding, we equilibrated beads in lysis buffer (50mM Tris-HCl, 300mM NaCl (pH 8.0)) and rocked at 4°C for 10 minutes. Beads were isolated by magnet and decanted. We repeated equilibration two additional times, then added 1mL of clarified lysate to beads and incubated at 4°C for 1hr up to overnight, while rotating. Isolated beads were washed in a 10-fold excess (relative to bead volume) of wash buffer #1 (50mM Tris-HCl, 300mM NaCl, 40mM Imidazole, 0.5% Triton-X (pH 8.0)). We then washed beads in 1mL of wash buffer #2 (50mM Tris-HCl, 300mM NaCl, 10mM Imidazole, 0.5% Triton-X (pH 8.0)). Finally, we washed beads three times in lysis buffer (removing imidazole and detergent). Protein was eluted using 3 times the resin bed volume with elution buffer (50mM Tris-HCl, 300mM NaCl, 500mM Imidazole pH 8.0). We repeated elution two additional times, typically finding the highest yield in the first two elutions. We dialyzed protein following manufacturer directions for the Pierce Slide-A-Lyzer mini dialysis with a 3.5kDa cutoff into 50mM Tris-HCl, 150mM NaCl at pH. 7.8. Proteins were aliquoted and frozen at −80°C.

### SILAC standard production

We transformed the !Lys, !Arg auxotroph cells from Dr. Ron T. Hay (29) with sVd-EGFRCtail and optimized the SILAC protocol (taking two rounds of MS analysis) to achieve higher than 97% incorporation across the protein. We adapted our protein production protocol and nickel based purification protocol for increased purity of SILAC protein. We added 0.5% glucose to the initial overnight growth phase in LB to minimize the leaky production of protein in the absence of SILAC reagents. The overnight was then pelleted and washed twice with 1xM9 salts to remove all traces of LB (resuspending in M9 salts in excess and pelleting for one wash step), before being resuspending pellets in a 4:1 final volume to initial volume (e.g. a 10mL overnight would be resupsended in 40mL of induction media). Induction media was a M9 minimal media (1x M9 salts, 50*µ*g/mL carbenicillin, 1mM MgSO_4_, 0.5*µ*g/mL thiamine/B1 vitamin, 1*µ*g/mL biotin, trace metals 1*µ*L/mL, 0.5% glucose, 50*µ*g/mL lsyine and arginine). L-Lysine-HCl and L-Arginine-HCl were used during optimization processes and SILAC supplementation used L-Lysine-2HCl 13C6,15N2 and L-Arginine-HCl 13C6,15N4. Post-washing pellets were equilibrated in the supplemented M9 media for 1hour at 37°C, then IPTG was added (1mM) to induce protein and grown for 4 hours at 37°C before pelleting and commencing with lysis, purification, and dialysis as described previously.

### In vitro kinase reaction

In vitro kinase reactions were performed during Nickel-based purification. Following the first three wash steps, beads were resuspended in clarified E. Coli kinase lysate. These lysates were produced from single transformant TOP10 cultures induced for kinase and lysed in kinase reaction buffer (50mM Tris-HCl, 10mM MgCl2, 0.1mM EDTA, 2mM DTT, 0.01% Brij 23 at pH 7.5), supplemented with protease and phosphatase inhibitors. After resuspension of beads in kinase supernatant, the buffer was supplemented with a stock of 10mM ATP (for a final concentration of 0.5mM ATP). Kinase reaction proceeded for 1 hour at 30°C with shaking, at which point the kinase lysate was removed and three wash steps were performed using 50mM Tris-HCl, 300mM NaCl, 0.5% TX-100, and 40mM imidazole. Elutions proceeded as normal as described for purification.

### GST Purification

We used Pierce Glutathione Agarose and followed the manufacturer’s base protocol for GST-based purification. Beads were equilibrated in buffer (50mM Tris, 150mM NaCl) and protein capture was performed for one hour to overnight at 4°C. This same buffer was used for washing (three times), with a 10-fold excess volume, with rotation. We used either PreScission protease-based elution (as described below) or with reduced glutathione (20mM reduced glutathione, 50mM Tris-HCl pH 8.0, 150-300mM NaCl, 5% glycerol).

### PreScission Protease

We adapted a protocol provided the manufacturer (GenScript). In solution protease was performed by add 1*µ*L of protease to 75*µ*L of clarified lysate. Protease was incubated while nutating at 4°C overnight. For PreScission of protein while on the beads (either for targeting cleavage during nickel purification of substrates or for elution of GST-fused proteins), we resuspended beads in 50mM Tris-HCl, 150mM NaCl, and 3mM DTT, then added PreScission protease and incubated overnight at 4°C with nutation or inversion mixing. For 300*µ*L of buffer volume, we added 1.5*µ*L of enzyme. For nickel-based purification of substrates (post PreScission removal of targeting sequence), we proceeded with three wash steps and elution as described above. **Western-based analysis:** Samples were reduced in Laemmli loading buffer (Boston BioProducts) and boiled for 10 minutes at 95C, unless they contained imidazole, where we boiled at 70°C for 5 to 10 minutes to avoid breaking protein bonds. We performed standard SDS-PAGE analysis using purchased precast NuPAGE gels from Invitrogen, typically using 4-12% gradients for single 10% gels, running in a MOPS SDS running buffer (Novex Invitrogen). Transfer to nitrocellulose membranes occurred in a buffer (Novex Invitrogen) with 20% methanol (membranes were pre-wetted in transfer buffer). Pre-stained ladder from LI-COR (Chameleon Duo) was used along with IRDye-680 and −800 secondary antibodies for infrared scanning. We used Intercept Blocking Buffer (LI-COR), diluted 1:1 in Tris Buffered Saline (TBS) for blocking membranes (one hour at room temperature or overnight at 4°C) and preparing primary and secondary antibodies. Following primary incubation (1 hour at room temperature or overnight at 4°C) and secondary incubation (20-30 minutes at room temperature at 1:15,000, we washed membranes in a TBS-T (1% tween solution). Antibody stripping was done with 0.2N NaOH for 5-30 minutes, as needed for efficient stripping or with Restore Stripping Buffer (ThermoScientific). Membranes were scanned on a LI-COR Odyssey system.

### Phos-Tag

The Phos-Tag protocol was developed using Wako Phos-tag™ SDS-PAGE guidebook. Briefly, samples were mixed with 6X Laemmli’s SDS-PAGE sample buffer and heated at 95°C for 10 min. Samples were separated by hand-cast 10% Bis-Tris PAGE gels that were supplemented to 50 *µ*M Phos-Tag acrylamide and 100 *µ*M ZnCl2 with a 4% stacking gel. Phos-Tag reagent was purchased from Wako Chemicals, Japan or ApexBio Technology LLC, Houston, Texas. Phos-Tag gels were electrophoresed in running buffer (50 mM MOPS, 50 mM Tris Base, 0.1% SDS, 5 mM sodium bisulfite, pH 7.8) at 50 milliamps for 2.5 hrs at 4°C. The gels were transferred to nitrocellulose membranes in 1!NuPAGE Transfer Buffer with 20% methanol at 35V for 1.5 hrs. Standard immune-staining and imaging performed as described previously.

### SH3 domain far western

ABL SH3 domain was cloned into a the parent pGEX (GST-fusion) backbone, with the addition of the Avi-Tag sequence and purified following procedures described above for PreScission-based purification of GST-fused proteins. We biotinylated purified SH3 domain according to the manufacturer directions of the Avidity LLC BirA ligase kit (BirA500), estimating SH3 domain substrate concentration based on a molecular weight of 38.5kDa (determined by Coomassie). We reacted 10nM of SH3 domain with 2.5ug of BirA ligase for 40 minutes at 30°C. We tested biotinylation by western and a streptavidin-conjugated infrared dye (LI-COR), prior to using biotinylated SH3 domain in far western. For far western, we load controlled polyproline sequences in the 21-mer constructs based on BCA-based analysis and two rounds of preliminary Myc-based westerns and densitometry. Following SDS-PAGE separation and transfer and a preliminary blocking step, we incubated the membrane with SH3 domain (23*µ*g of biotinylated SH3 domain in 10mL of blocking buffer) for one hour at 4°C. Membranes were washed as per usual western-based approaches and 680IRdye-conjugated streptavidin was added for one hour at 4°C (at 1:1500), prior to final washes and LICOR-based imaging.

### *Mass spectrometry based analysis of EGFRCtail*. Proteomics Sample Preparation

The samples were digested as previously described using a modification of the filter-aided sample preparation method (41). The samples were mixed with 200 *µ*L of 100 mM Tris-HCL buffer, pH 8.5 containing 8M urea (UA buffer) and 25 mM dithiothreitol (DTT) and transferred on top chamber of a 10,000 MWCO cutoff filtration unit (Millipore, part# MRCPRT010). Proteins were reduced at RT for 45min followed by spinning in a microcentrifuge (Eppendorf 5424, Eppendorf Cat. No. 2231000767) at 14,000 rcf for 30 min. The flow through was discarded and the proteins were alkylated by addition of 100 *µ*L of 15 mM Iodoacetamide (Pierce, Ref. No. A39271) in UA buffer to the top chamber of the filtration unit and gyrating at 550 rpm in the dark at RT for 45 min using a Thermomixer (Eppendorf, Thermomixer R). The filter was spun at 14,000 rcf for 30 min and the flow through discarded. The urea buffer was exchanged into digestion buffer (DB), 50 mM ammonium bicarbonate buffer, pH 8. Two sequential additions of DB (200 *µ*L) with centrifugation after each addition to the top chamber were performed. The top filter units were transferred to a new collection tube and 0.25 mU of Lys-C (Wako Chemicals, cat.no. 129-02541) was added and samples were digested for two hours at 37°C followed by addition of 1 *µ*g of sequencing-grade Trypsin (Promega, Ref. No. V5113) and digestion overnight at 37°C. The filters were spun at 14,000 rcf for 30 min to collect the peptides in the flow through. The filter was washed with 50 *µ*L 100 mM ammonium bicarbonate buffer and the wash was collected with the peptides.

In preparation for desalting, peptides were acidified to 1% (vol/vol) TFA final concentration. The peptides were desalted using two micro-tips sequentially (PGC, BIOMEKNT3CAR) (Glygen) on a Beckman robot (Biomek NX), as previously described in Chen et al. (42). The peptides were eluted with 60% (vol/vol) acetonitrile in 0.1% TFA (vol/vol) and dried in a Speed-Vac (Thermo Scientific, Model No. Savant DNA 120 concentrator). The peptides were dissolved in 20 µl of 1% (vol/vol) acetonitrile in water. An aliquot (10%) was removed for quantification using the Pierce Quantitative Fluorometric Peptide Assay kit (Thermo Scientific, Cat. No. 23290). The remaining peptides were transferred to autosampler vials (Sun-Sri, Cat. No. 200046), dried and stored at −80°C for LC-MS analysis.

### LC-MS Data Acquisition

The peptides were analyzed using UPLC Orbitrap mass spectrometry (43) with the modifications described below. The samples in 1% (vol/vol) formic acid (FA) were loaded in 2.5 µl onto a 75 µm i.d. ! 50 cm Acclaim PepMap 100 C18 RSLC column (Thermo-Fisher Scientific) on an EASY nanoLC (Thermo Fisher Scientific) at a constant pressure of 700 bar at 100% A (0.1%FA). Prior to sample loading the column was equilibrated to 100%A for a total of 11 µl at 700 bar pressure. Peptide chromatography was initiated with mobile phase A (1% FA) containing 2%B (100%ACN, 1%FA) for 5 min, then increased to 20% B over 100 min, to 32% B over 20 min, to 95% B over 1 min and held at 95% B for 19 min, with a flow rate of 300 nl/min (Naegle_007) or 250 nl/min (Naegle_008). The data was acquired in data-dependent acquisition (DDA) mode. Full-scan mass spectra were acquired with the Orbitrap mass analyzer with a scan range of m/z = 325 to 1500 (Naegle_007, evaluation of SILAC incorporation) or 375-1500 (Naegle_008 the first batch of phosphoproteins with target of 5:1 spike-in) and a mass resolving power set to 70,000. Ten data-dependent high-energy collisional dissociations (HCD) were performed with a mass resolving power set to 17,500, a fixed lower value of m/z 100, an isolation width of 2 Da, and a normalized collision energy setting of 27. The maximum injection time was 60 ms for parent-ion analysis and product-ion analysis. The target ions that were selected for MS/MS were dynamically excluded for 20 sec. The automatic gain control (AGC) was set at a target value of 1e6 ions for full MS scans and 1e5 ions for MS2. Peptide ions with charge states of unassigned, plus one, greater than plus eight were excluded for HCD acquisition.

### Parallel Reaction Monitoring for EGFR Tryptic Peptides

Tryptic peptides representing tyrosine-phosphorylatable portions of the recombinant EGFR-tail protein were selected in silico based on the construct sequence. Peptides lacking tyrosine residues were selected for use as surrogate protein load normalization measured by data-dependent analysis of recombinant EGFR-tail tryptic digests and the 3 peptides with the highest MS1 signal intensity were chosen. During initial assessment of SILAC incorporation efficiency (Naegle_007), it was observed that the residual intensity of signal in the light isotope channel would be a confounding factor when attempting to quantify peptide percent phosphorylation based on the loss of signal when monitoring the unmodified peptide sequences. Thus, titration experiments were carried out to establish the optimal quantity of SILAC protein standard that would minimize light isotope signal from the standard while providing sufficient heavy isotope signal for quantitation (Naegle_013) – varying amounts (5*µ*g, 2*µ*g, 1*µ*g, and 0.5*µ*g) of EGFR C-tail protein were mixed with a constant amount (4*µ*l, 200ng) of heavy isotope labeled (SILAC) EGFR C-tail protein. For these titration experiments, data-dependent analyses were performed and measured ratios of light-to-heavy peptide were determined via manual selection of peak boundaries using the Skyline software (44) and integration of MS1 peak areas. These preliminary data-dependent analyses also served as a resource for selection of the optimal precursor charge states and fragment ions for downstream parallel reaction monitoring (PRM) analyses.

The PRM methods were designed to target 24 tryptic peptides in their isotopically light and SILAC heavy (U-13C, 15N lysine and arginine) forms, employing a scheduling strategy with 15-minute windows. For quantitative analyses, peaks of interest were determined by expert curation of the mass spectrometric data, manual selection of peak boundaries using the Skyline software and integration of the 3 most intense y-ions. For binary calls of peptide presence/absence, observation of the 3 most intense y-ions was required. Each phosphotyrosine containing EGFR fragment chromatogram is provided in Supplementary Information.

The mass spectrometry proteomics data have been deposited to the ProteomeXchange Consortium via the PRIDE (45) partner repository with the dataset identifier PXD060364.

## Supporting information

Supplementary Figures - MS2 calls

## Acknowledgment

Research reported in this publication was supported by the National Cancer Institute of the National Institutes of Health under Award Numbers R21CA212726 and R33CA259451. The content is solely the responsibility of the authors and does not necessarily represent the official views of the National Institutes of Health. The authors wish to thank the many collaborators who provided expertise and DNA constructs listed in the main text, including all depositions on Addgene or by mail, and also Diane Lidke and Elizabeth Bailey (PD-1) and Linda Pike (EGFR). The authors wish to thank Emily Gale, Kellie Kuenning, Roman Sloutsky, and Tom Ronan who contributed early ideas or effort in the very beginning stages of this project that ultimately shaped the project direction.

## Notes

https://doi.org/10.6084/m9.figshare.28299335

https://doi.org/10.6084/m9.figshare.28296383

### Competing Interest Statement

The authors have declared no competing interest.

### Summary of Updates

Added links to public datasets, PRIDE deposition, and updated supplements for MS calls. Fixed a figure issue.

