## Supplementary Figures - MS2 calls for "A signaling inspired synthetic toolkit for efficient production of tyrosine phosphorylated proteins"

### Product Ion Chromatograms for Phosphtyrosine Peptides

MS2 spectra and calls

Pages 2-14 Vials 30-47

Pages 15-27 Vials 51-65

### Vial number mapping to substrate/kinase
### Vials 30-47

| Vial Number | Targeting | Kinase |
|---|---|---|
| 30 | none | ABL |
| 31 | p40 | ABL |
| 32 | none | LYN |
| 33 | p40 | LYN |
| 35 | 3BP1 | LYN |
| 36 | 3BP2 | LYN |
| 37 | none | SRC |
| 41 | none | ABL |
| 42 | p40 | ABL |
| 43 | none | LYN |
| 44 | p40 | LYN |
| 45 | p41 | LYN |
| 46 | 3BP1 | LYN |
| 47 | 3BP2 | LYN |

EGFR pY998 MHLPSPTDSNF**Y**R (not observed in vial 32, 33, 37, 43, 44)

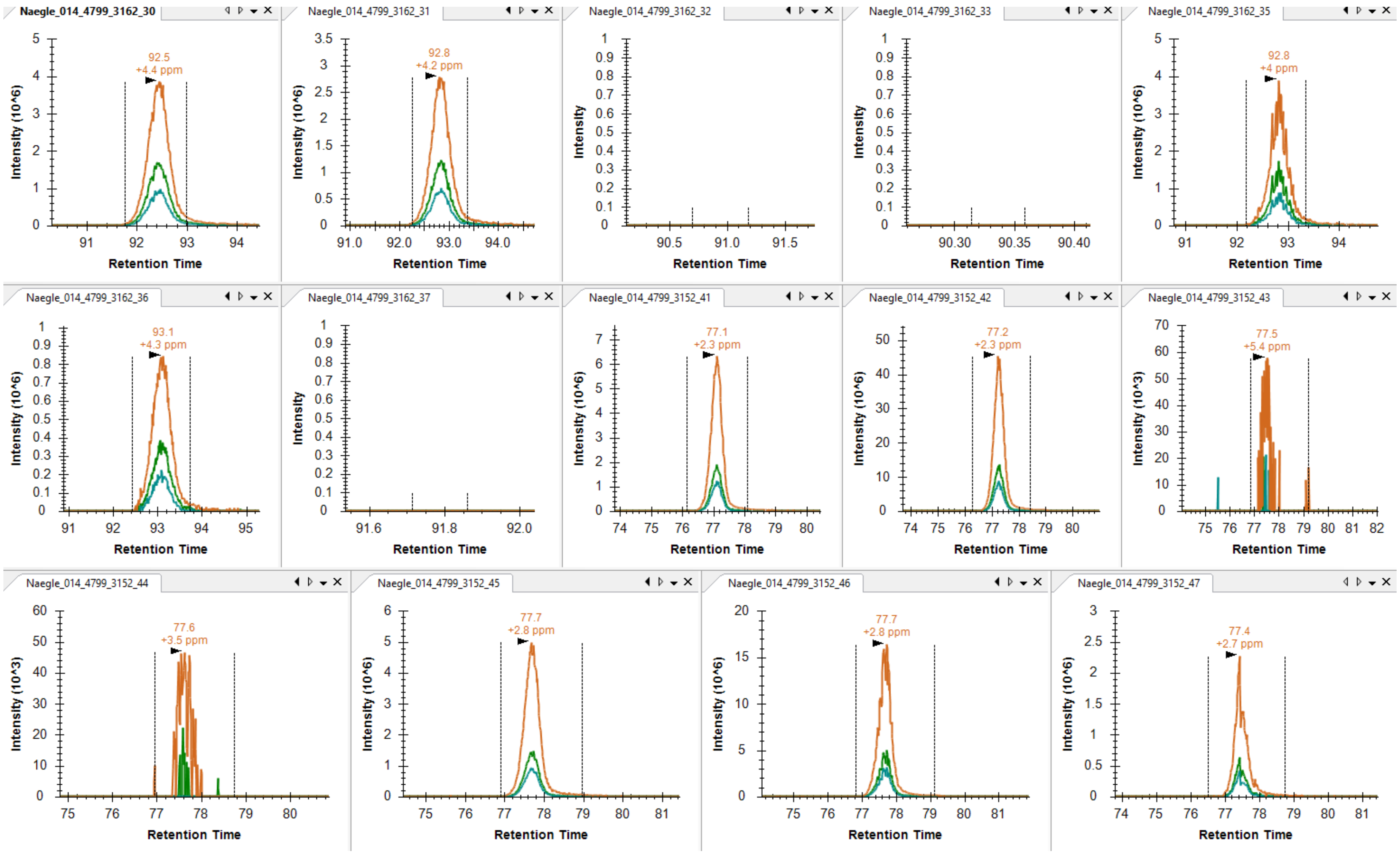

EGFR pY1016 ALMDEEDMDDVVDADE**Y**LIPQQGFFSSPSTSR (observed in vial 30, 32-36)

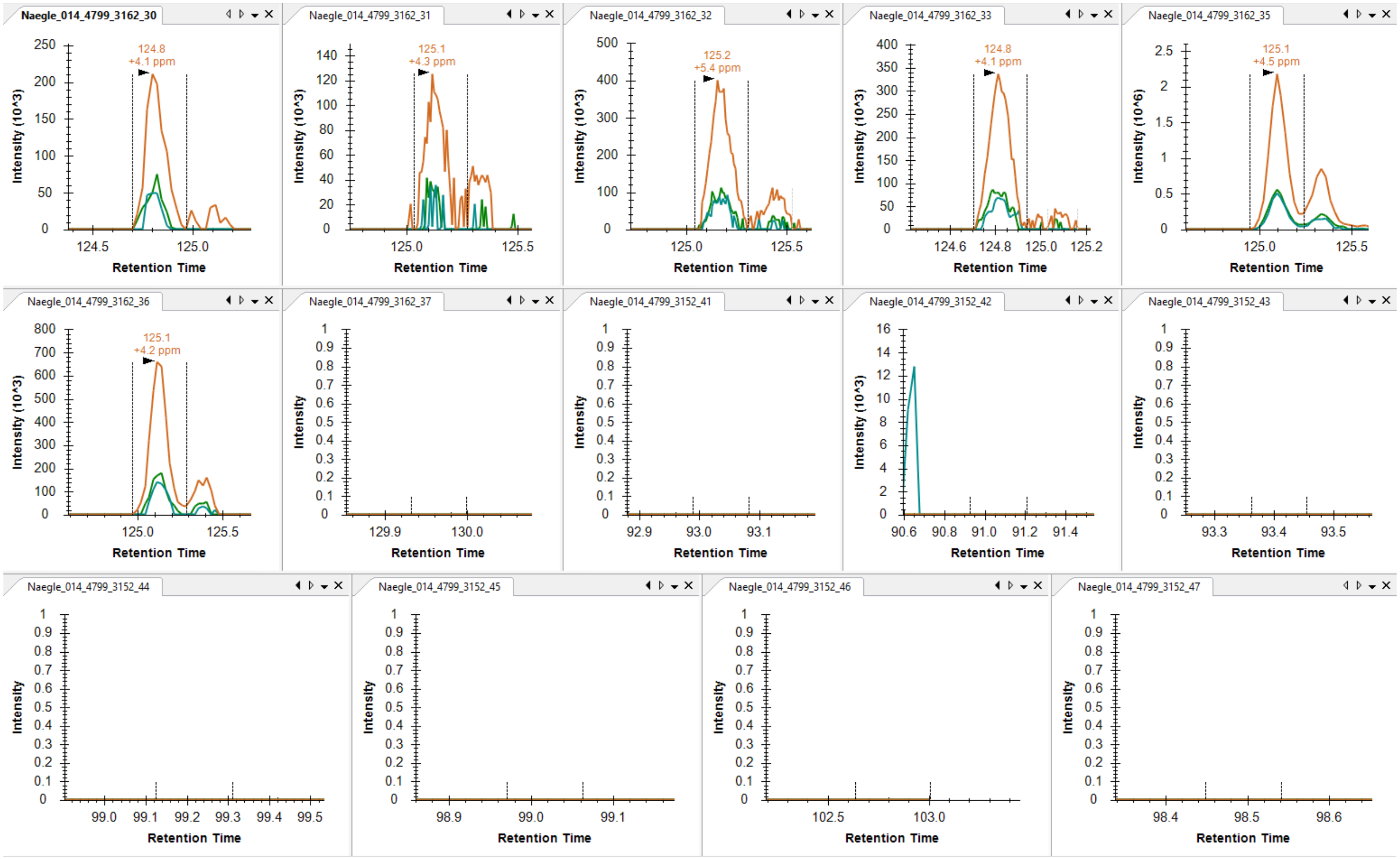

EGFR pY1092 YSSDPTGALTEDSIDDTFLPVPE**Y**INQSVPK (not observed in vial 33, 37, 43, 44, 47)

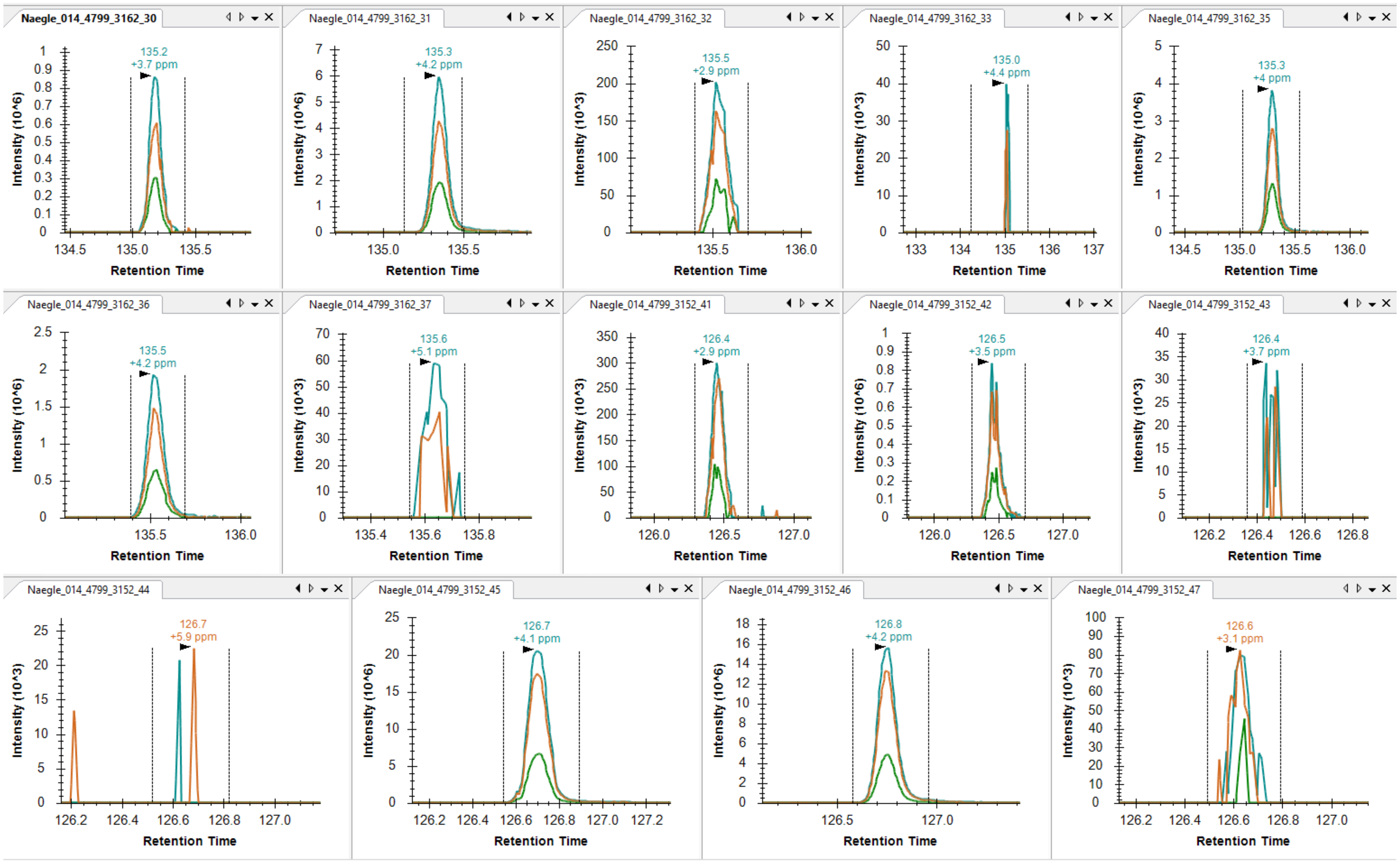

EGFR pY1069 **Y**SSDPTGALTEDSIDDTFLPVPEYINQSVPK (observed in vial 30, 31, 35, 36, 42, 45, 46)

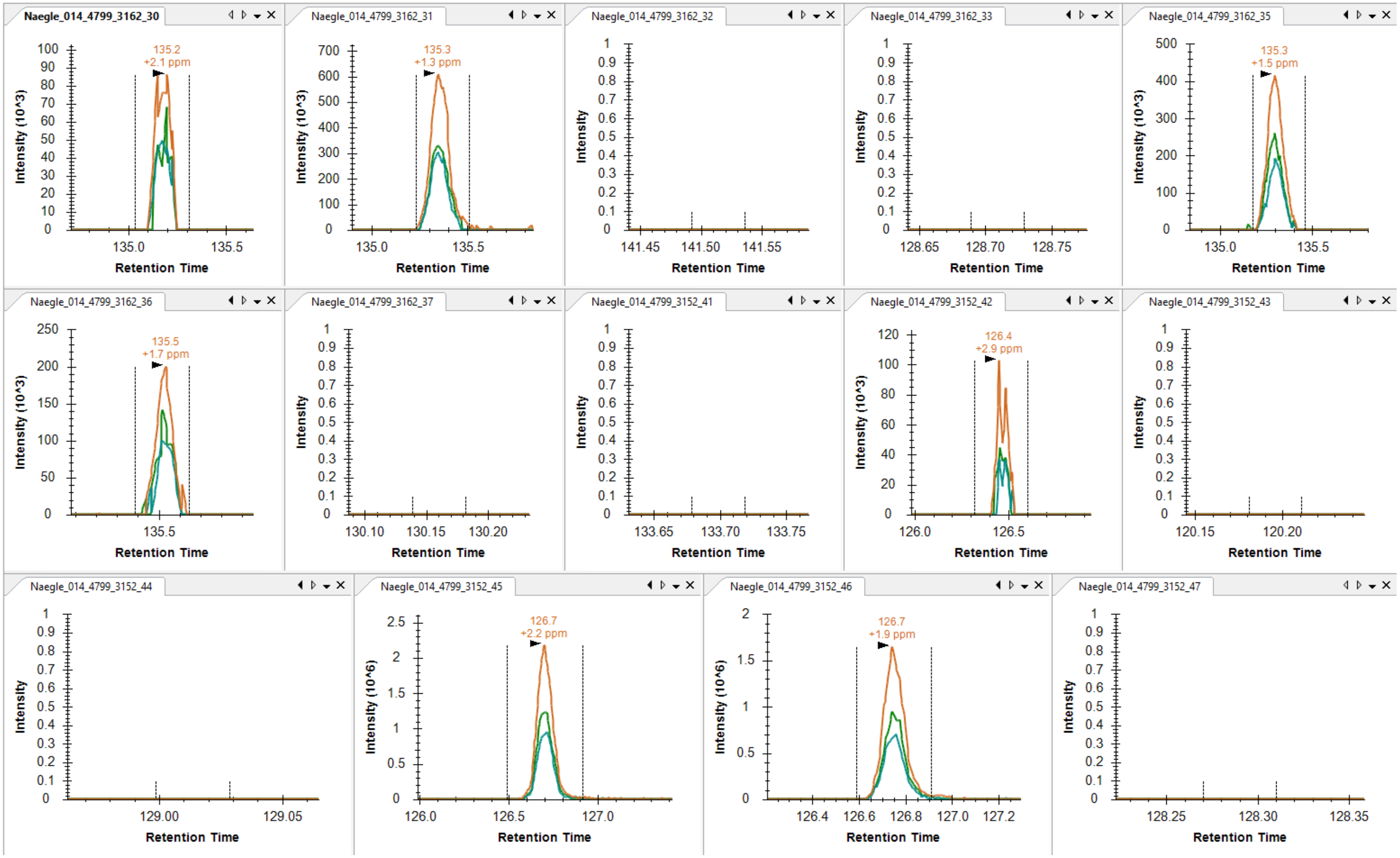

EGFR pY1069/pY1092

<u>**Y**</u>SSDPTGALTEDSIDDTFLPVPE<u>**Y**</u>INQSVPK (not observed)

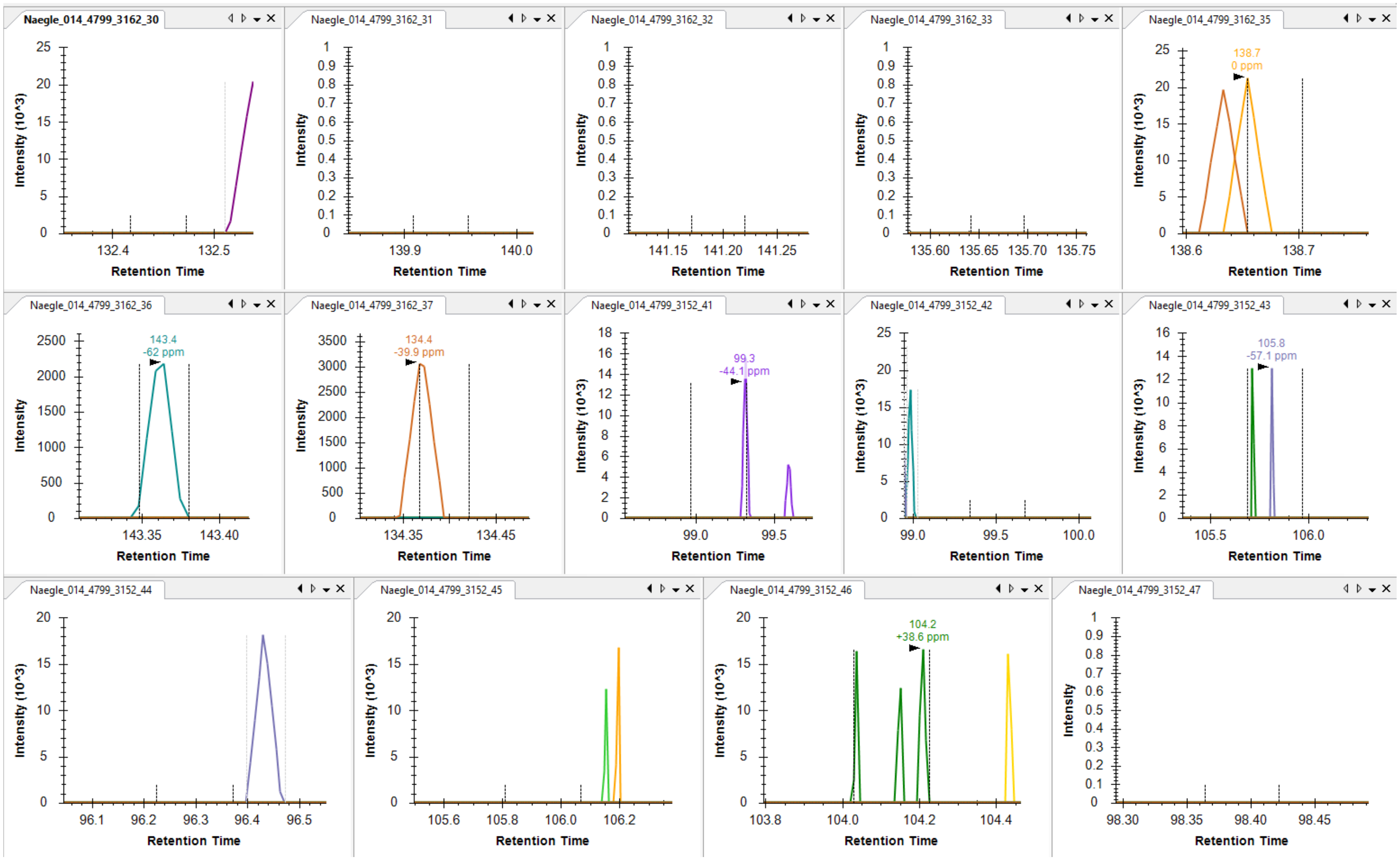

EGFR pY1110 RPAGSVQNPV**Y**HNQPLNPAPSR

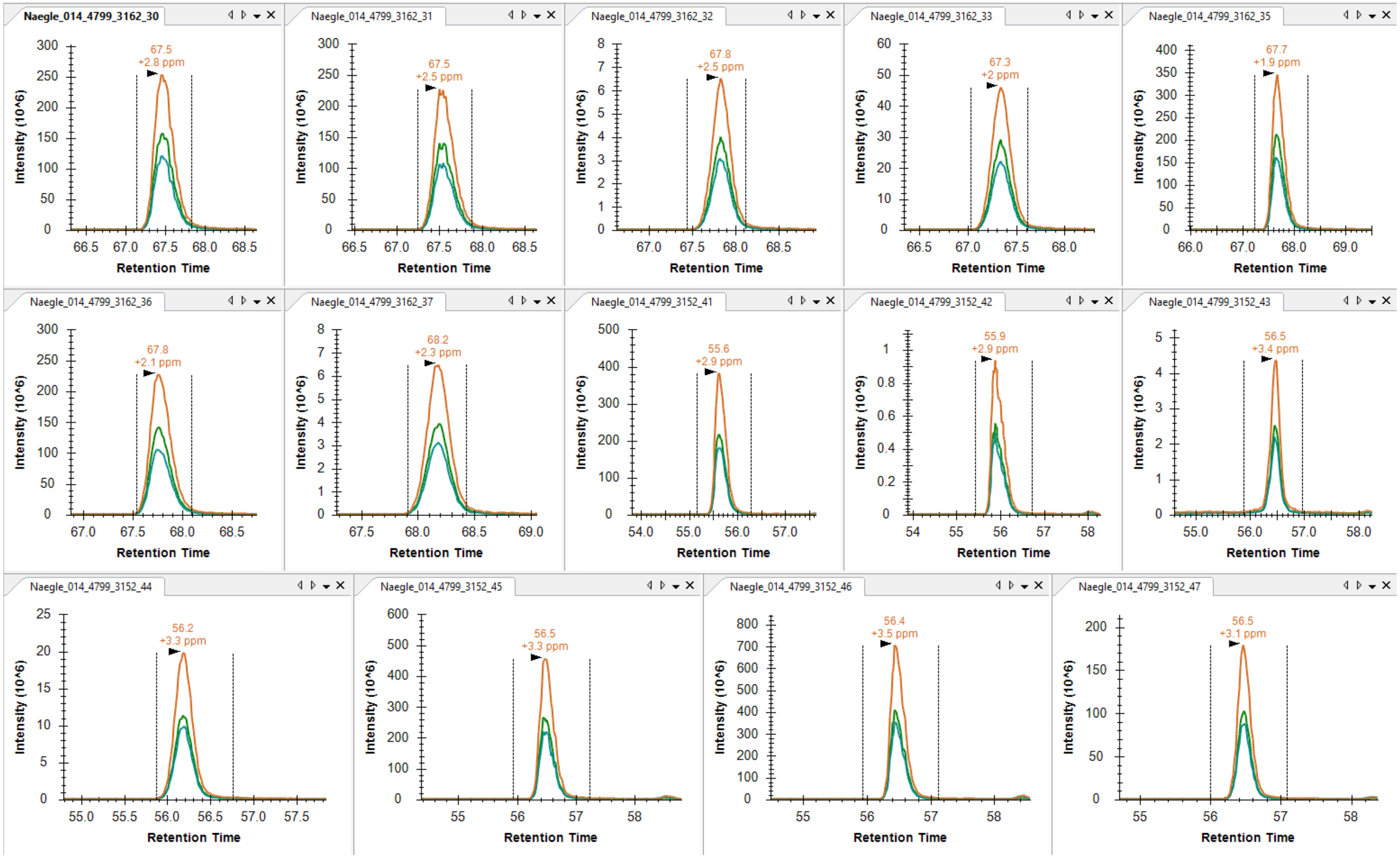

EGFR pY1110 PAGSVQNPVYHNQPLNPAPSR (not observed in vials 32, 37, 43)

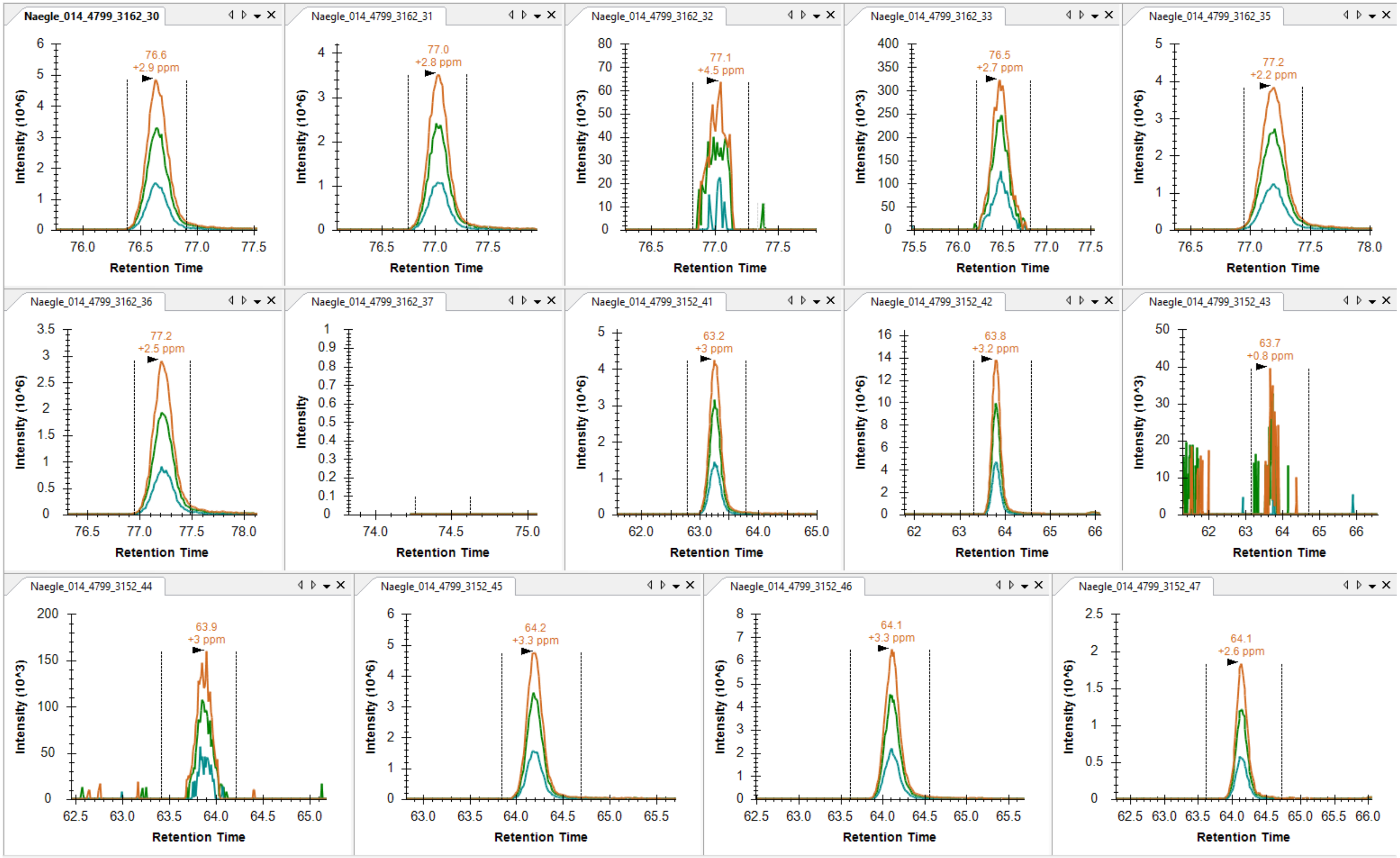

EGFR pY1125 DPH**Y**QDPHSTAVGNPEYLNTVQPT**C**VNSTFDSPAHWAQK (observed in vial 35, 42, 45, 46)

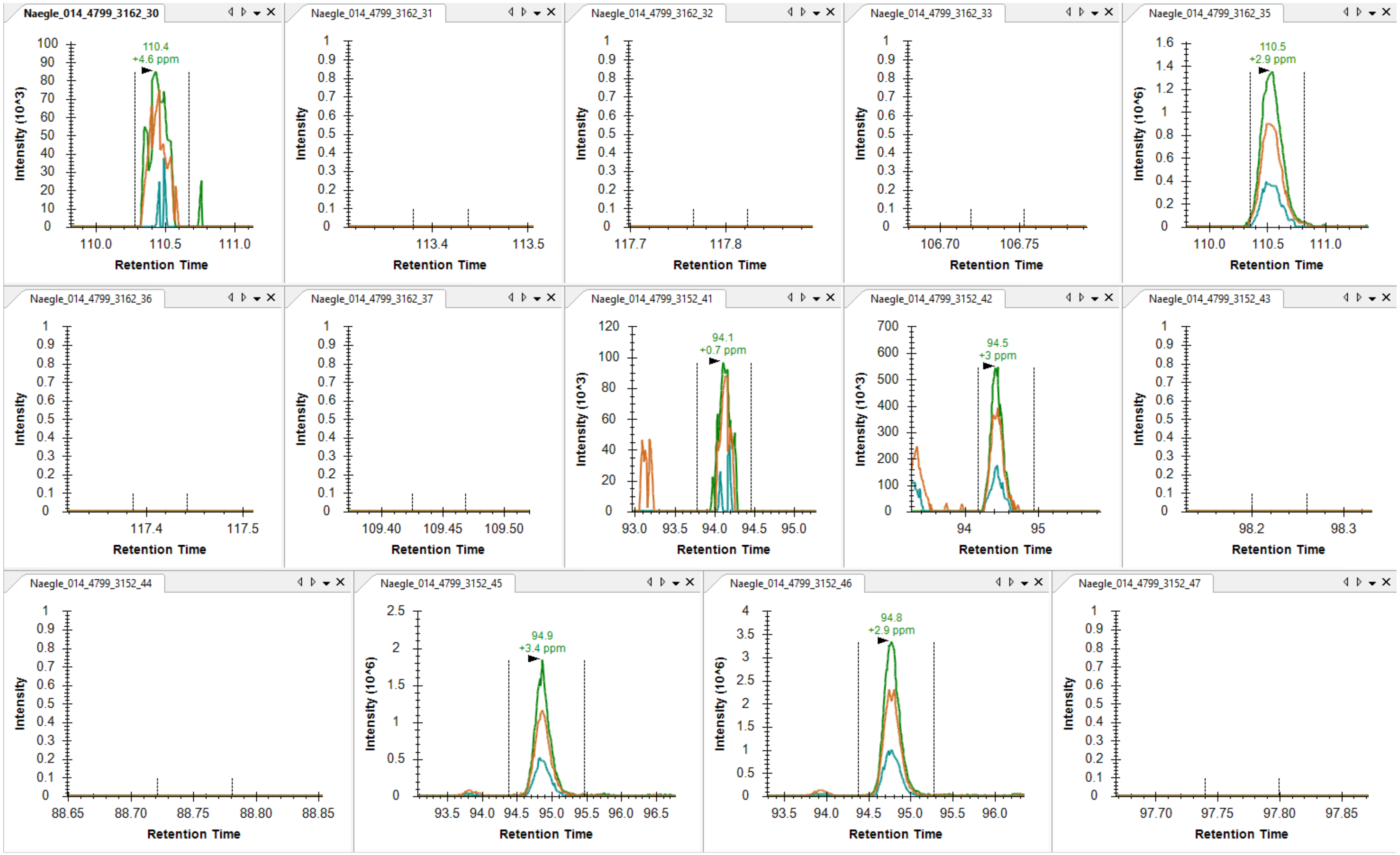

EGFR pY1138 DPHYQDPHSTAVGNPE**Y**LNTVQPT**C**VNSTFDSPAHWAQK (observed in vial 35, 42, 45, 46)

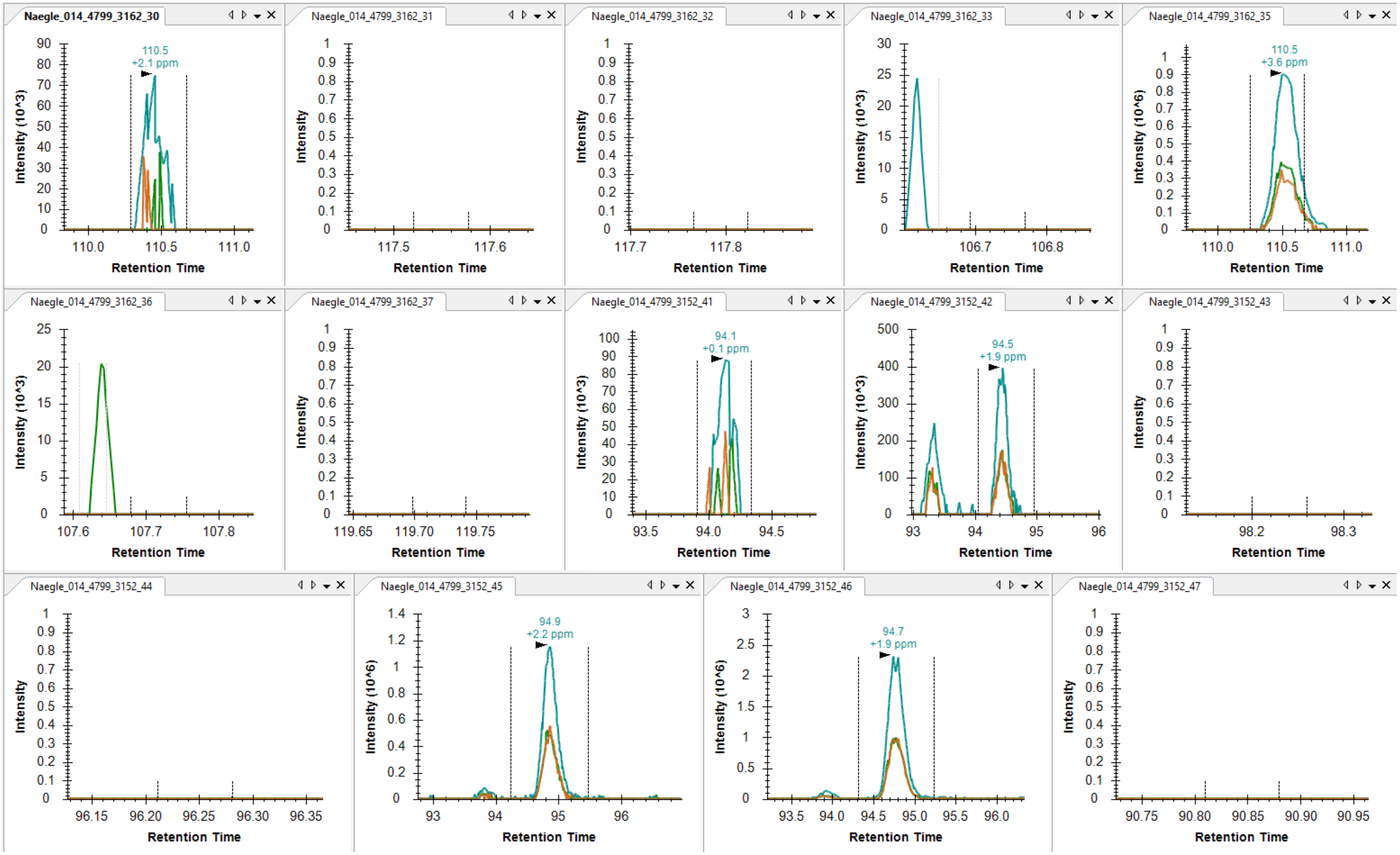

EGFR pY1125/pY1138

DPH**Y**QD[illegible]

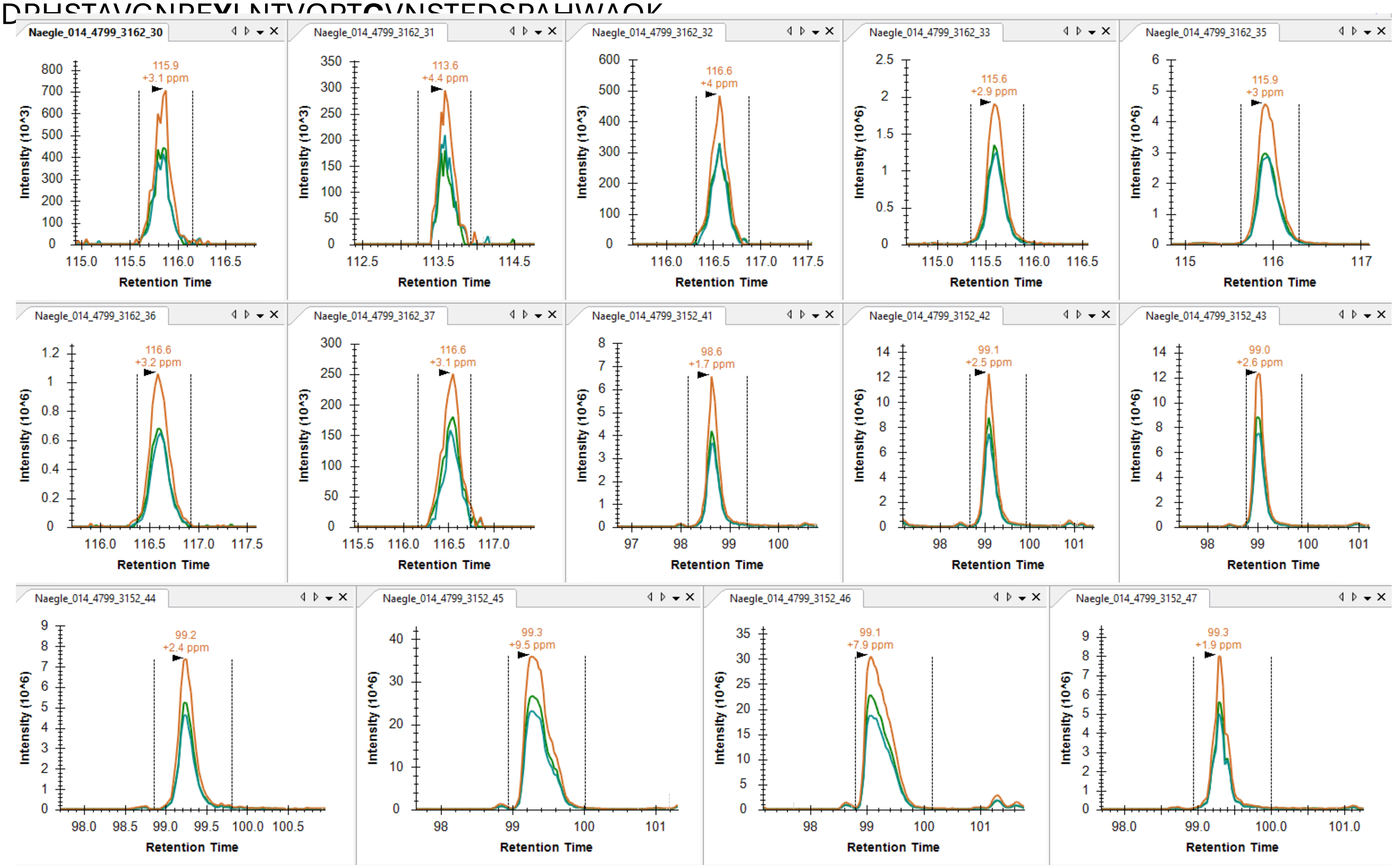

EGFR pY1172 GSHQISLDNPD<u>**Y**</u>QQDFFPK (not observed in vial 32, 33, 36, 37)

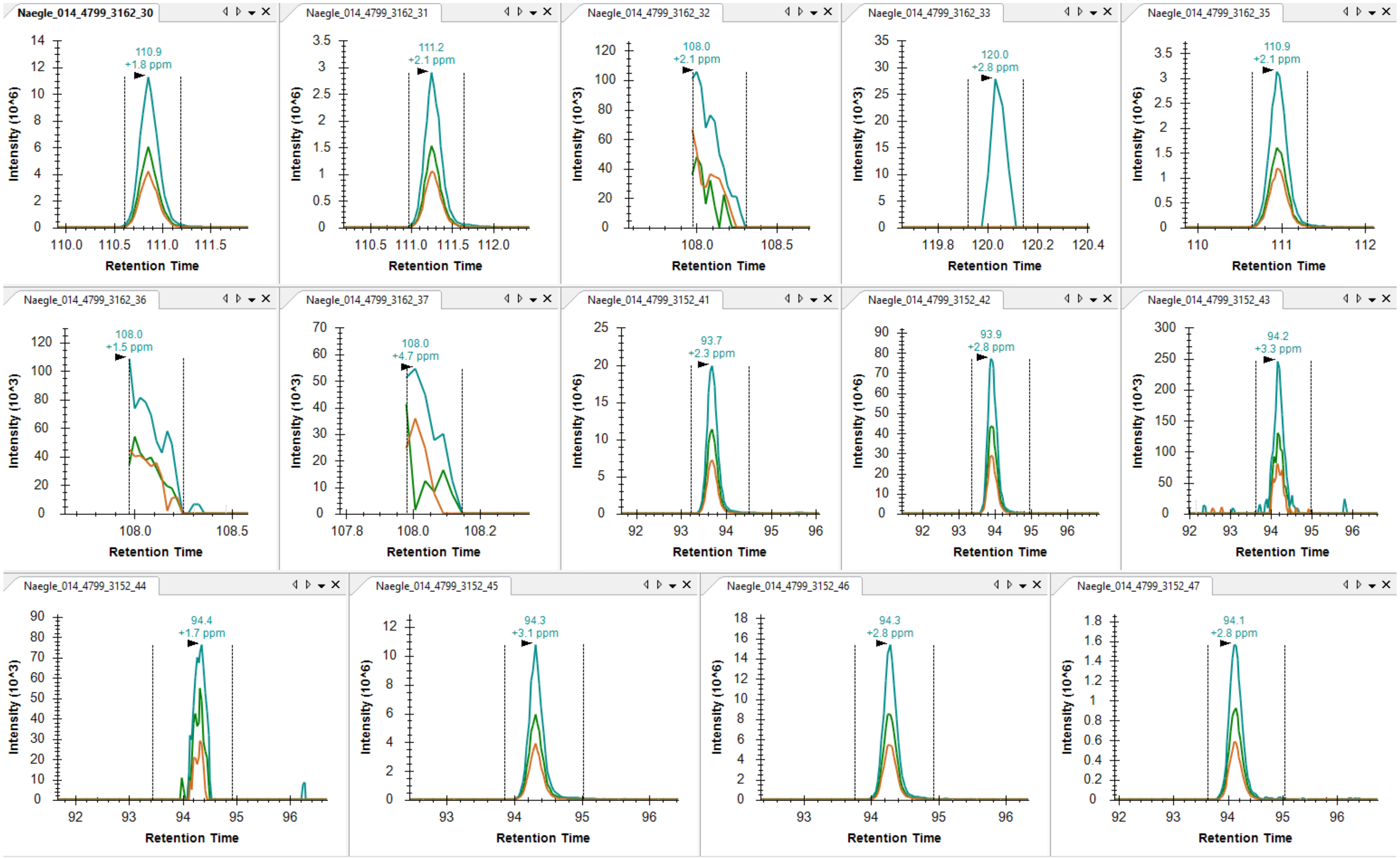

### EGFR pY1197 GSTAENAEYLR

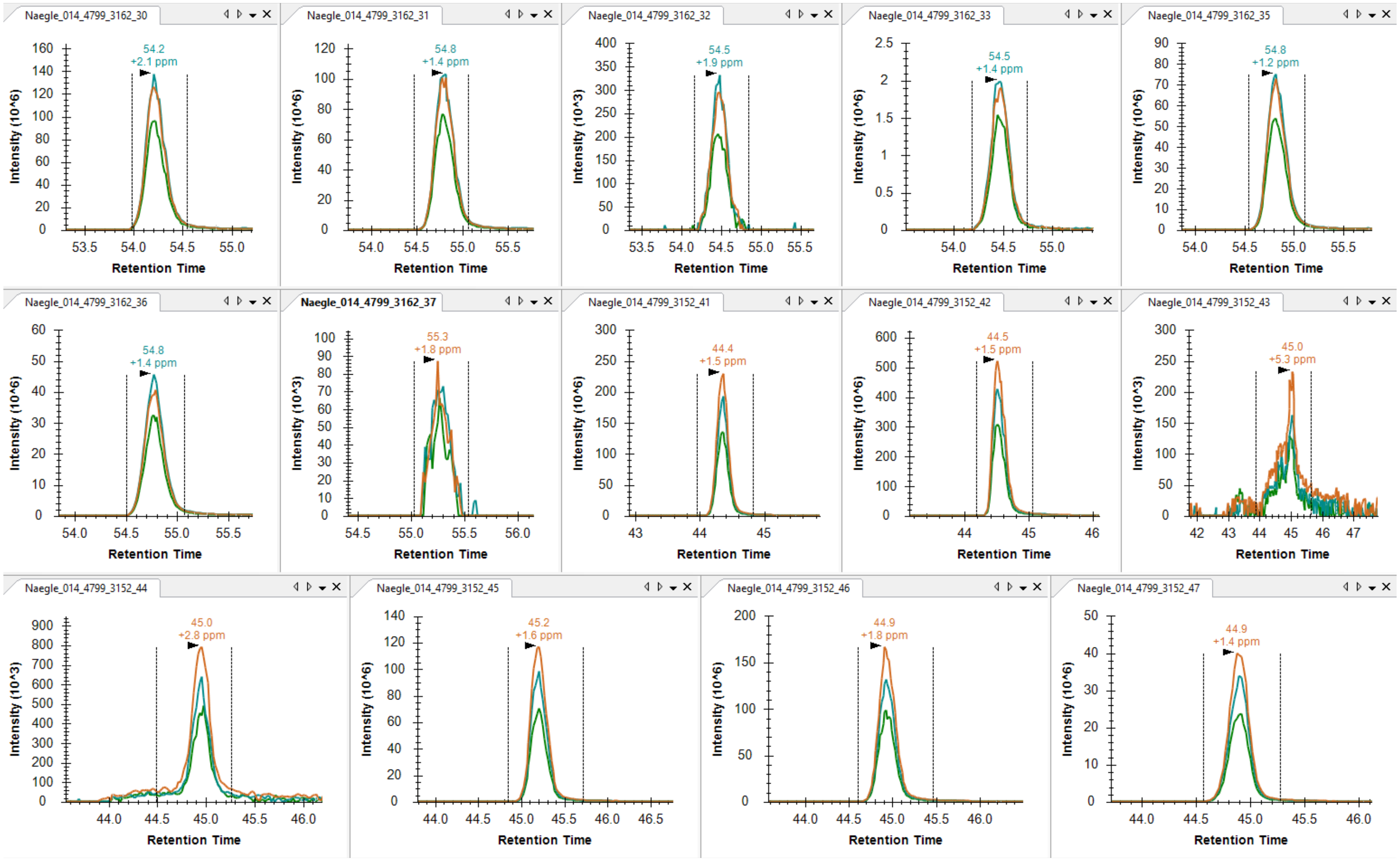

### Vial number mapping to substrate/kinase
### Vials 51-65

| Vial | Substrate Targeting | Kinase |
|---|---|---|
| 51 | p40 | BTK |
| 52 | none | EPHA4 |
| 53 | p40 | EPHA4 |
| 57 | none | SRC |
| 58 | p40 | SRC |
| 59 | none | BTK |
| 60 | none | BTK |
| 61 | p40 | BTK |
| 62 | p41 | LYN |
| 63 | p40 | SRC |
| 64 | none | EPHA4 |
| 65 | p40 | EPHA4 |

EGFR pY998 MHLPSPTDSNF**Y**R (observed in vial 58)

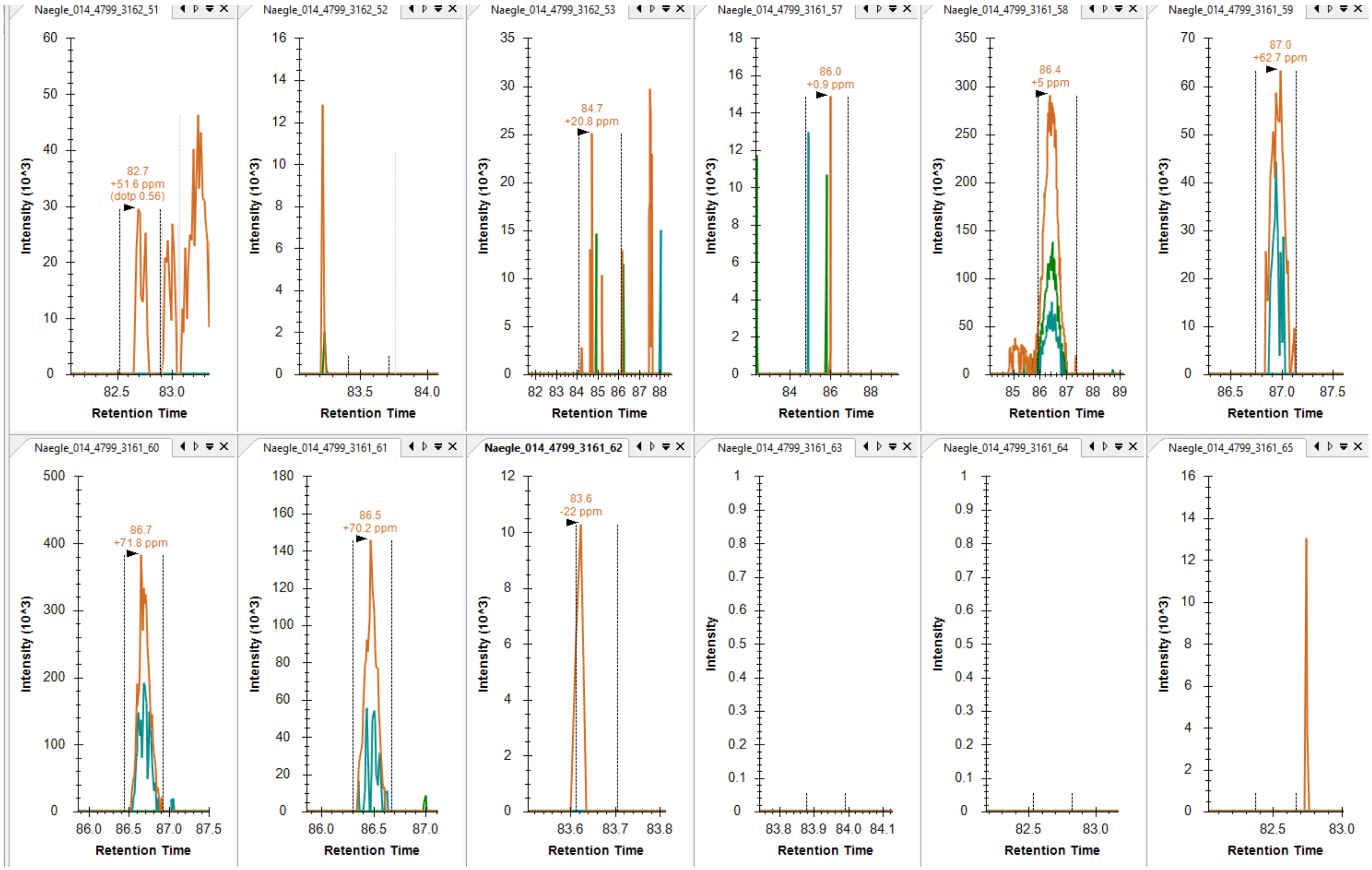

EGFR pY1016 ALMDEEDMDDVVDADE<u>Y</u>LIPQQGFFSSPSTSR (not observed in vial 57)

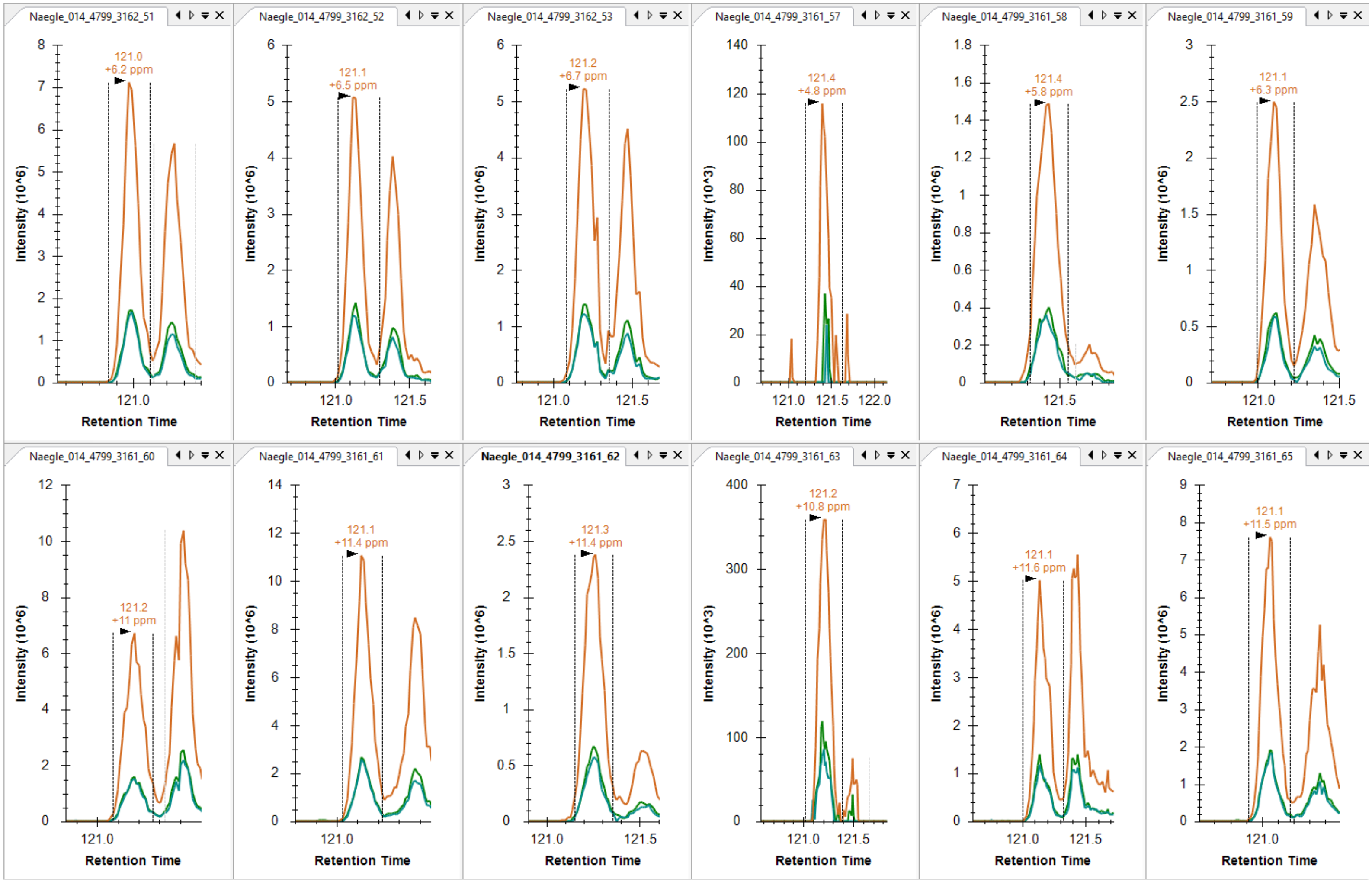

EGFR pY1092 YSSDPTGALTEDSIDDTFLPVPE**Y**INQSVPK (not observed)

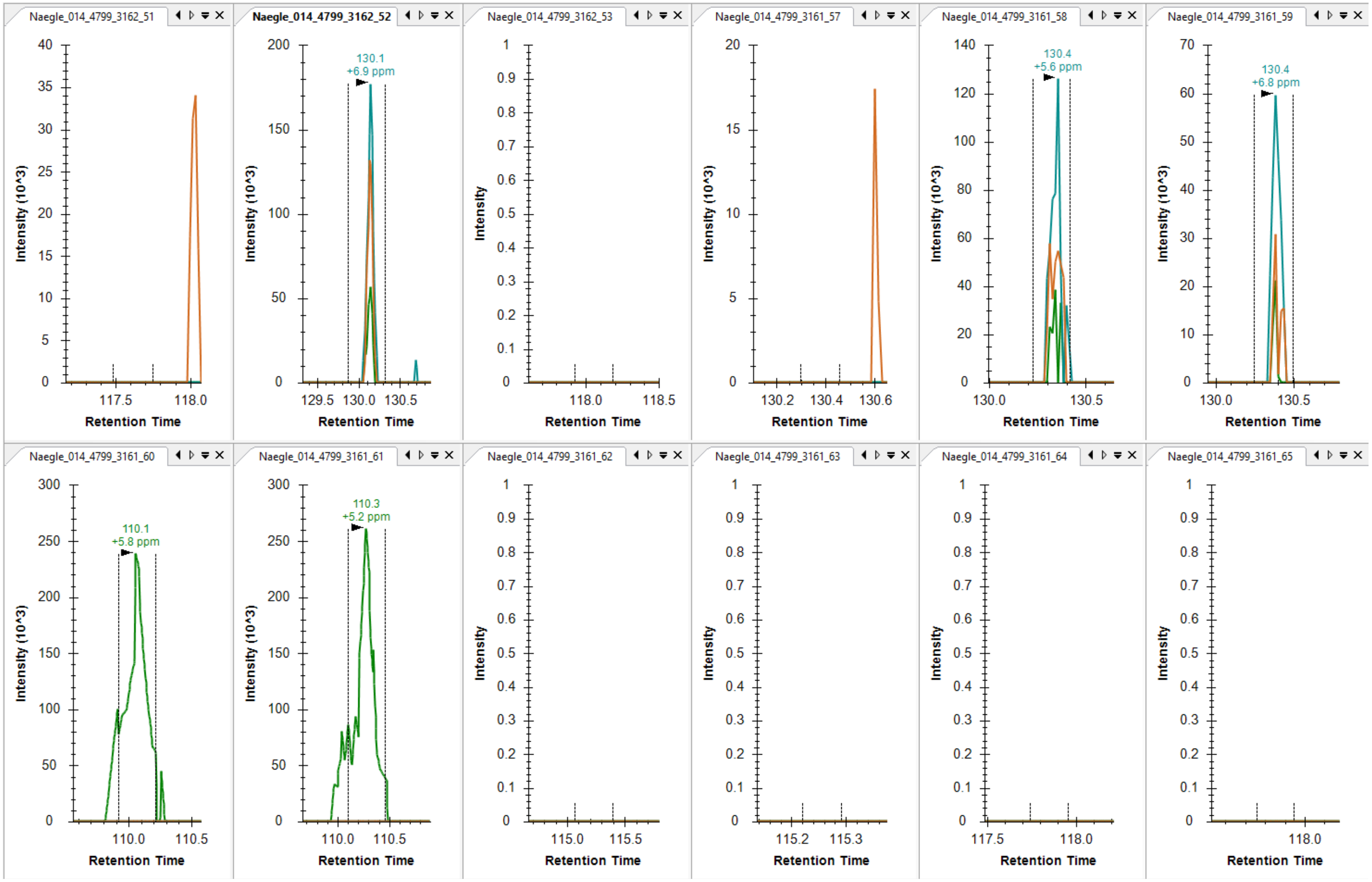

EGFR pY1069 **Y**SSDPTGALTEDSIDDTFLPVPEYINQSVPK (not observed)

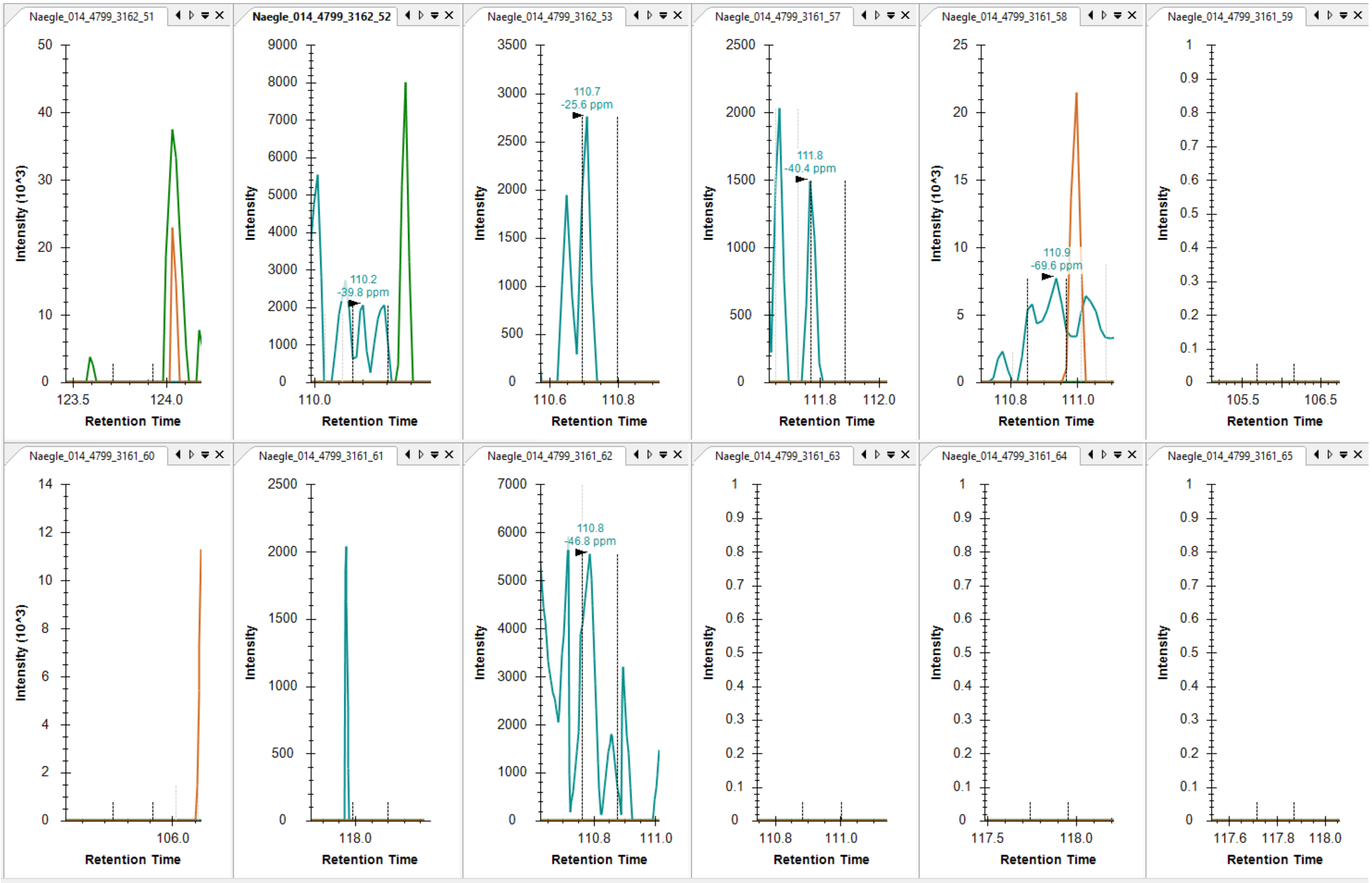

EGFR pY1069/pY1092 YSSDPTGALTEDSIDDTFLPVPEYINQSVPK (not observed)

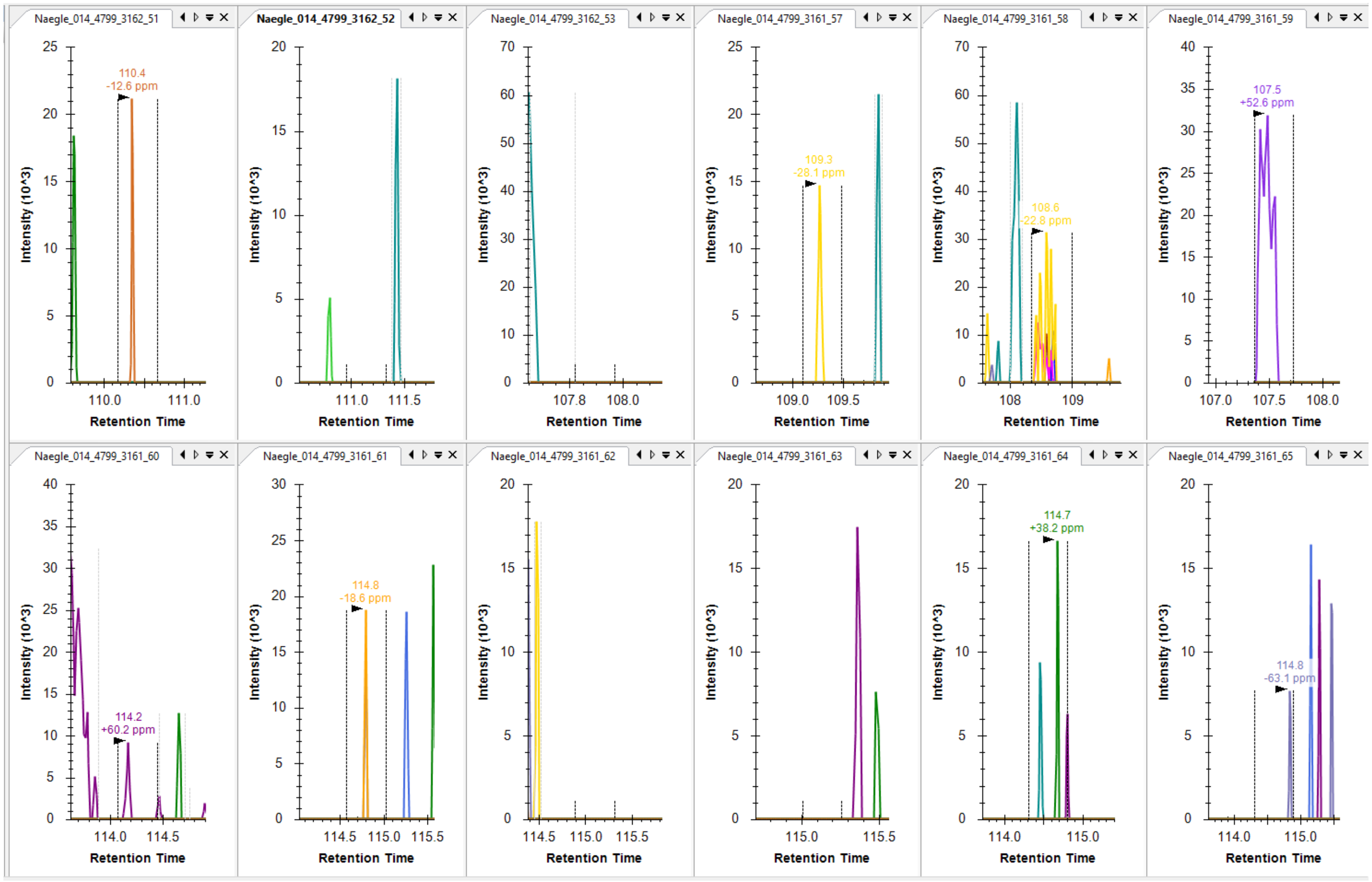

EGFR pY1110 RPAGSVQNPV**Y**HNQPLNPAPSR

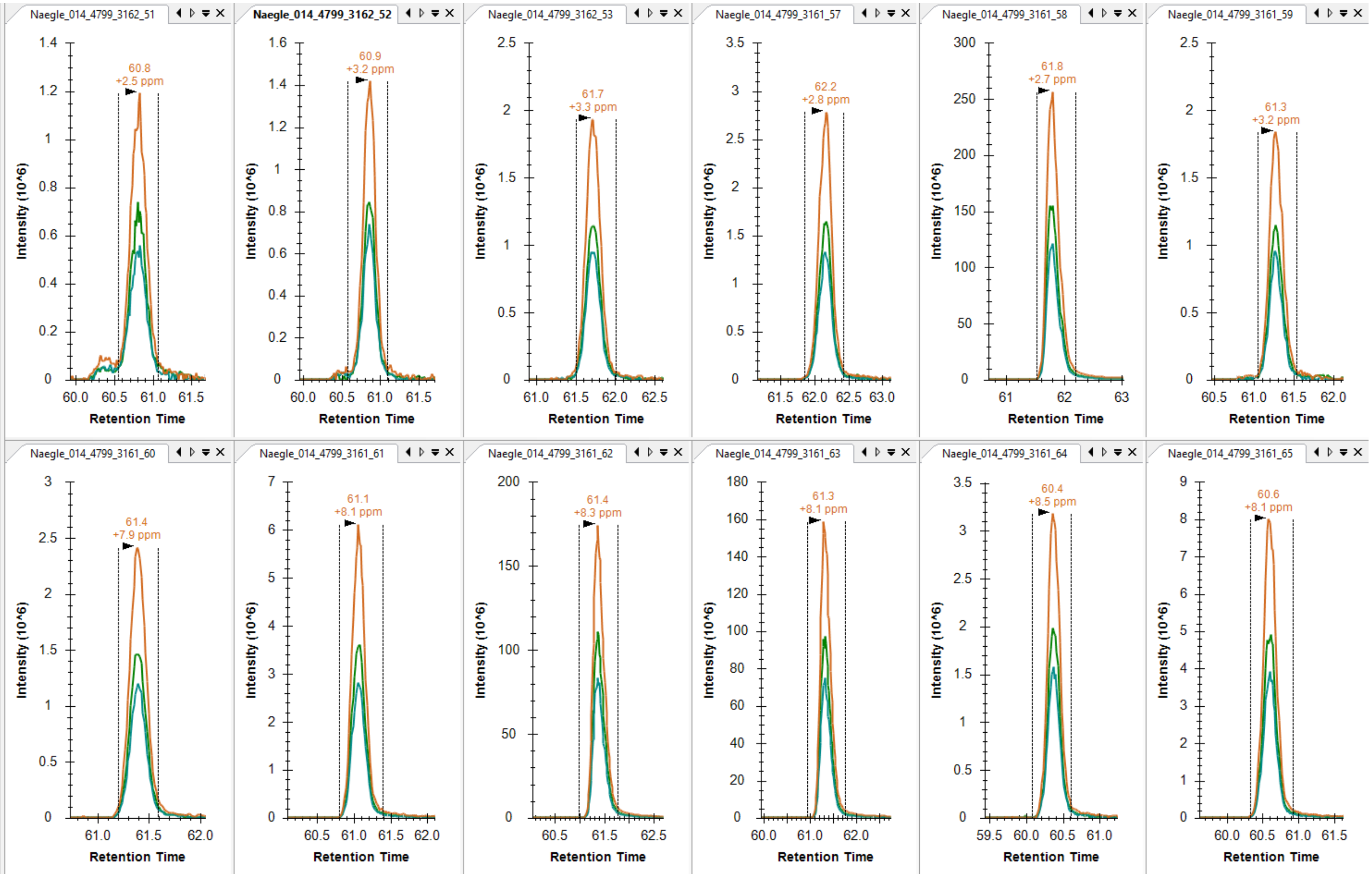

EGFR pY1110 PAGSVQNPV**Y**HNQPLNPAPSR (observed in vials 58, 62, 63)

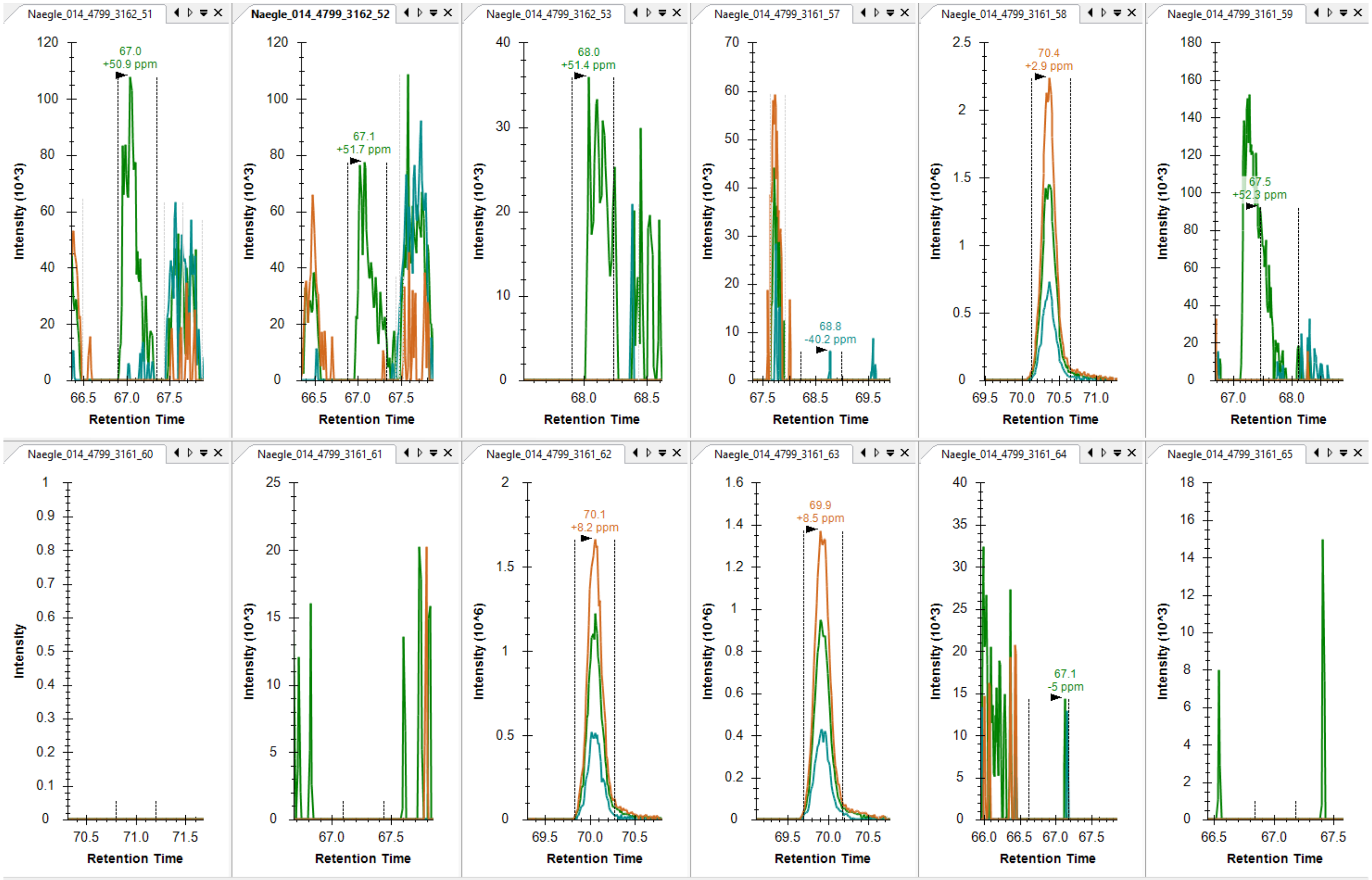

EGFR pY1125 DPH**Y**QDPHSTAVGNPEYLNTVQPT**C**VNSTFDSPAHWAQK (observed in vial 62, marginal signal in 65)

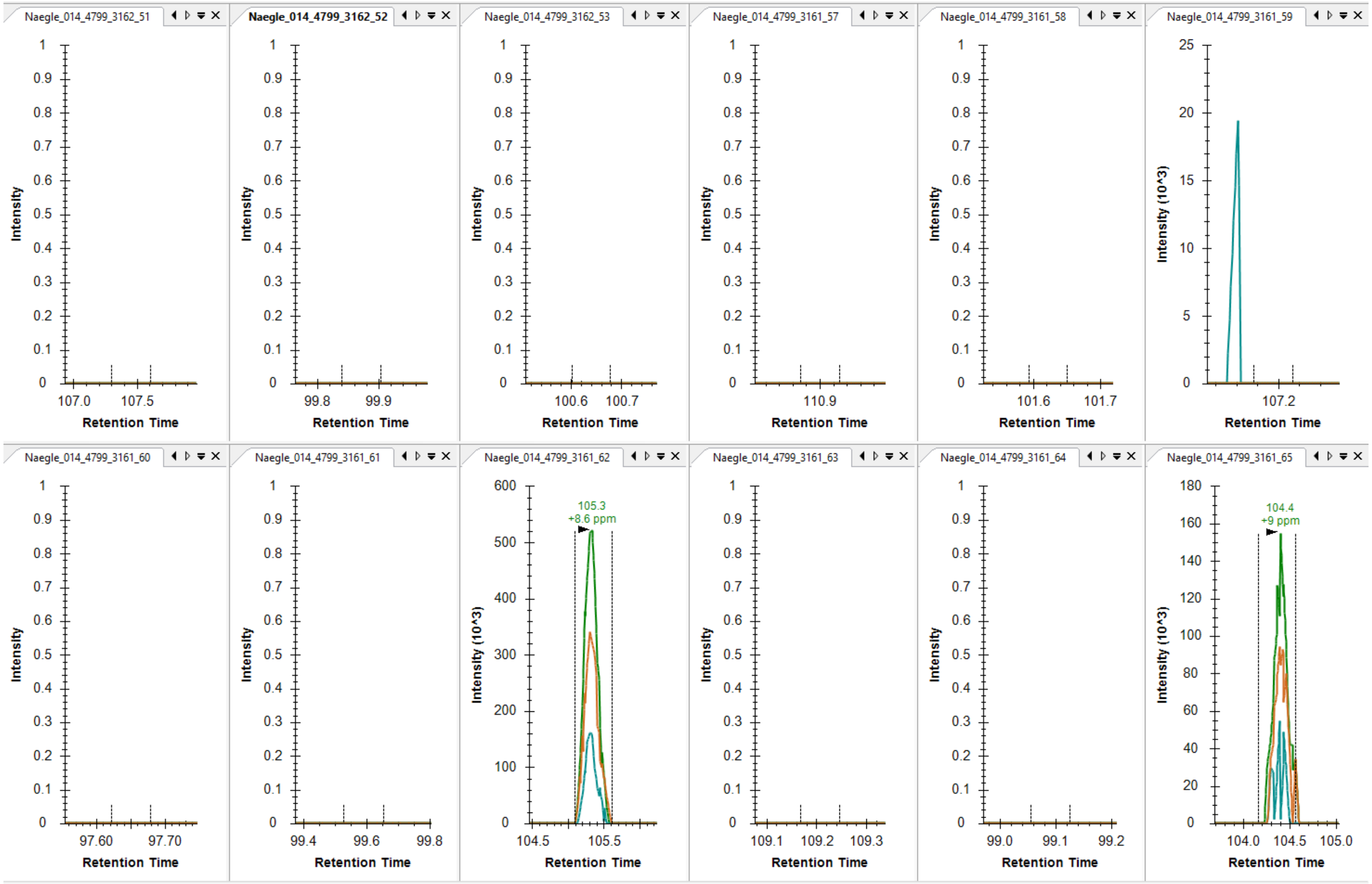

EGFR pY1138 DPHYQDPHSTAVGNPE**Y**LNTVQPT**C**VNSTFDSPAHWAQK (not observed)

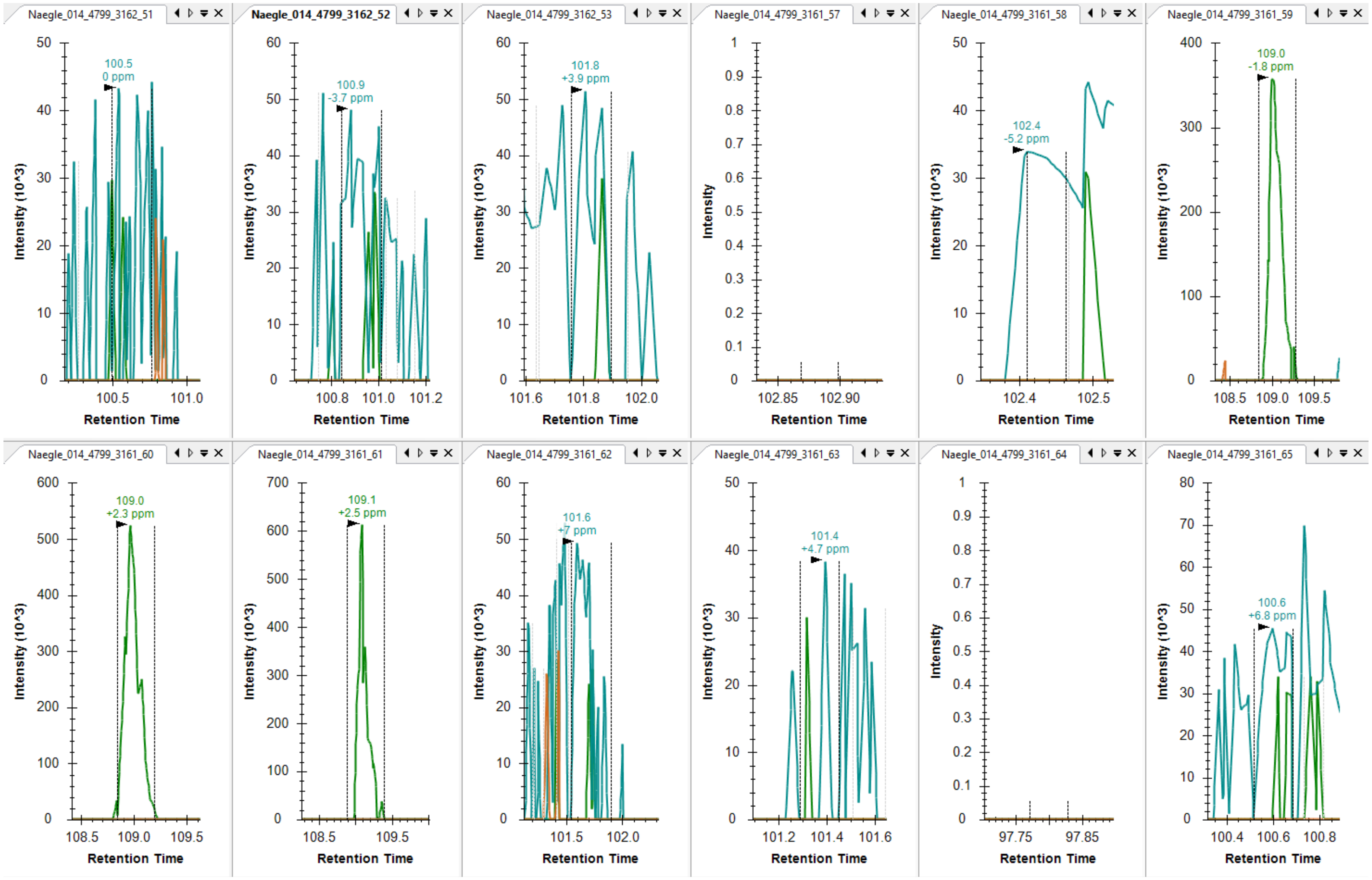

EGFR pY1125/pY1138 DPH**Y**QDPHSTAVGNPE**Y**LNTVQPT**C**VNSTFDSPAHWAQK

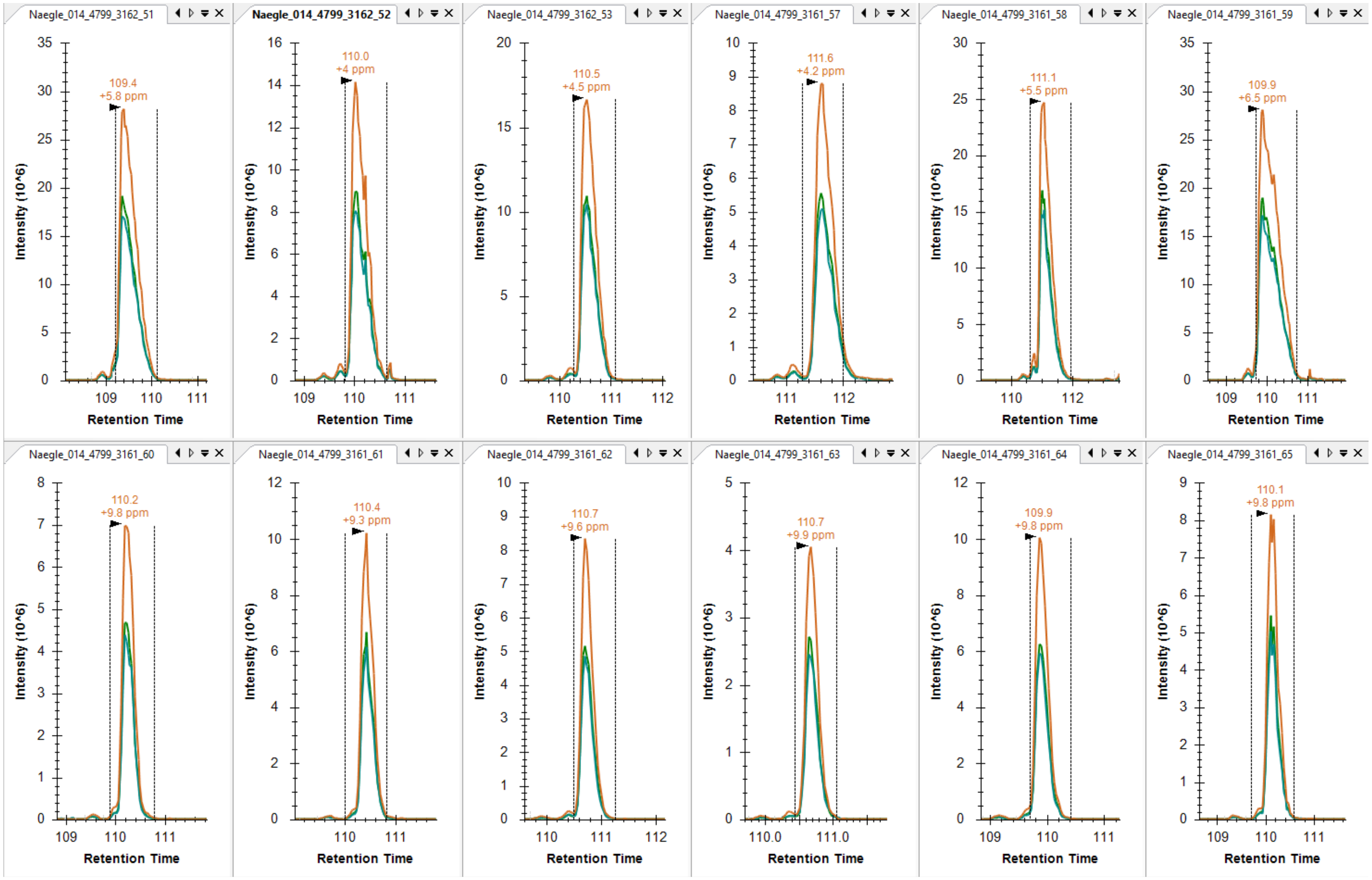

EGFR pY1172 GSHQISLDNPD**Y**QQDFFPK (not observed in vials 60, 61)

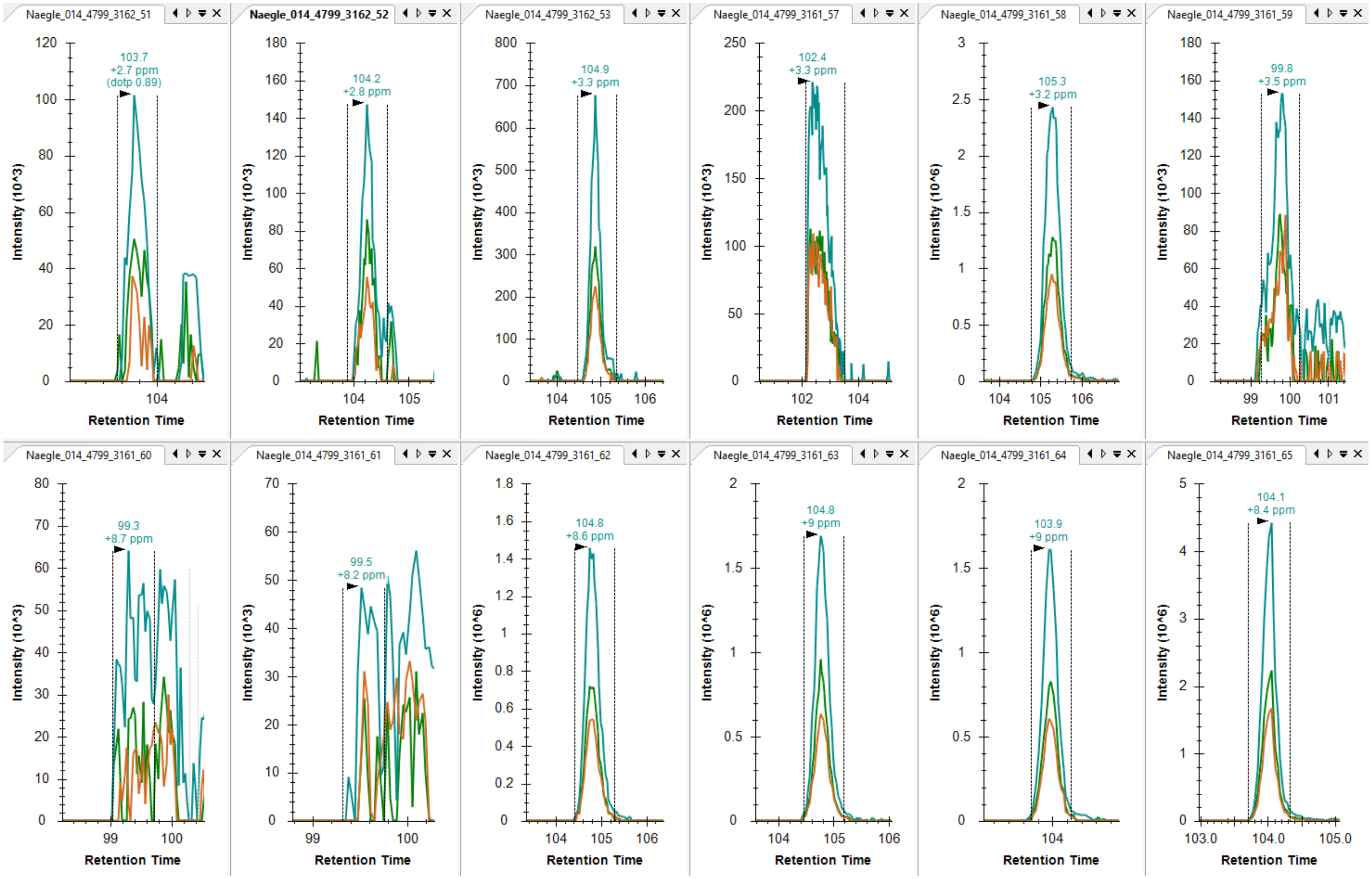

EGFR pY1197 GSTAENAE**Y**LR (observed in vials 58, 62-65)

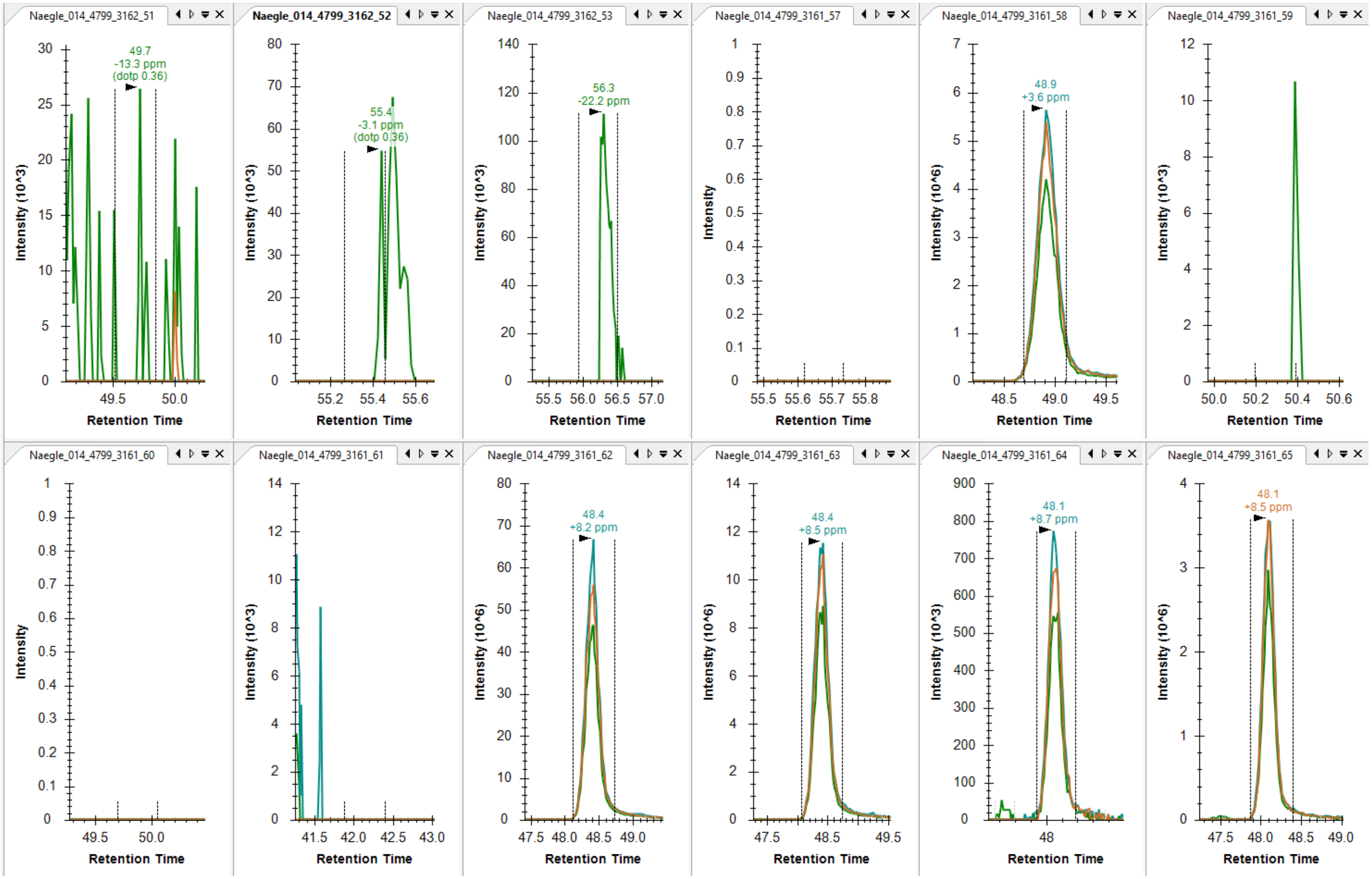
