## Supplementary Figures for "A signaling inspired synthetic toolkit for efficient production of tyrosine phosphorylated proteins": Supplement_SISAKiT_BioRxiv_Dec22.pdf

### This file includes:

**Supplementary Figure S1**: Example of activation loop mutagenesis and L-arabinose curves for kinases expression and activity.
**Supplementary Figure S2**: Phosphorylation of GST sequence.
**Supplementary Figure S3**: GST Y to F mutant (GSTm) does not bind GST resin.
**Supplementary Figure S4**: Testing Y67F in the smt3 domain on production of phosphoprotein.
**Supplementary Figure S5**: Testing E. coli strains engineered for dual induction of L-arabinose and galactose-based induction.
**Supplementary Figure S6**: Expression and purification of phosphosubstrate from three E. coli strains.
**Supplementary Figure S7**: Testing if central metabolism alterations can increase co-expression of kinase and substrate.
**Supplementary Figure S8**. Evaluation of in vitro, co-expression, and serial inductions on phosphoprotein yields.
**Supplementary Figure S10**. Testing the Light to Heavy ratios of non-phosphorylatable “barcode” peptides in the EGFRCtail across two sets.
**Supplementary Figure S11**: Lambda phosphatase treatment improves Coomassie-based protein estimations.
**Supplementary Figure S12**: 1069F control used to test antibody detection of PxxP tyrosines.
**Supplementary Figure S13**: Supernatant probes of the 21-mer co-expressions.
**Supplementary Figure S14**: Testing variability of kinase and substrate expression for two 21-mers.
**Supplementary Figure S15**: Low SNR with control highlights antibody issue.
**Supplementary Figure S16** Load control western shown here for SH3 far western experiment.

**Supplementary Table 1** Individual binary calls across pTyr EGFR Ctail peptides in the 25:1 experiment (014). Attached as Excel Book.
**Supplementary Table 2** Details of kinase and substrate proteins. Attached as Excel Book.
**Supplementary Table 3** Details of all antibodies and reagents used. Attached as Excel Book.

**Supplementary Table 1. Individual binary calls across pTyr EGFR Ctail peptides in the 25:1 experiment (014)**. Attached as Excel Book. This indicates if a quality peak was observed as detailed in methods for a site in the indicated reaction condition.

**Supplementary Table 2. Details of kinase and substrate proteins** Sheets indicate the type of information (kinase or substrate), sequencing and cloning details, and the regions spanned of human proteins covered. These describe the Snapgene DNA files that are included, which themselves have attached sequencing results.

**Supplementary Table 3. Details of all antibodies and reagents used.** Sheets include the reagent, part numbers, and manufacturer. Antibodies also include our dilutions.

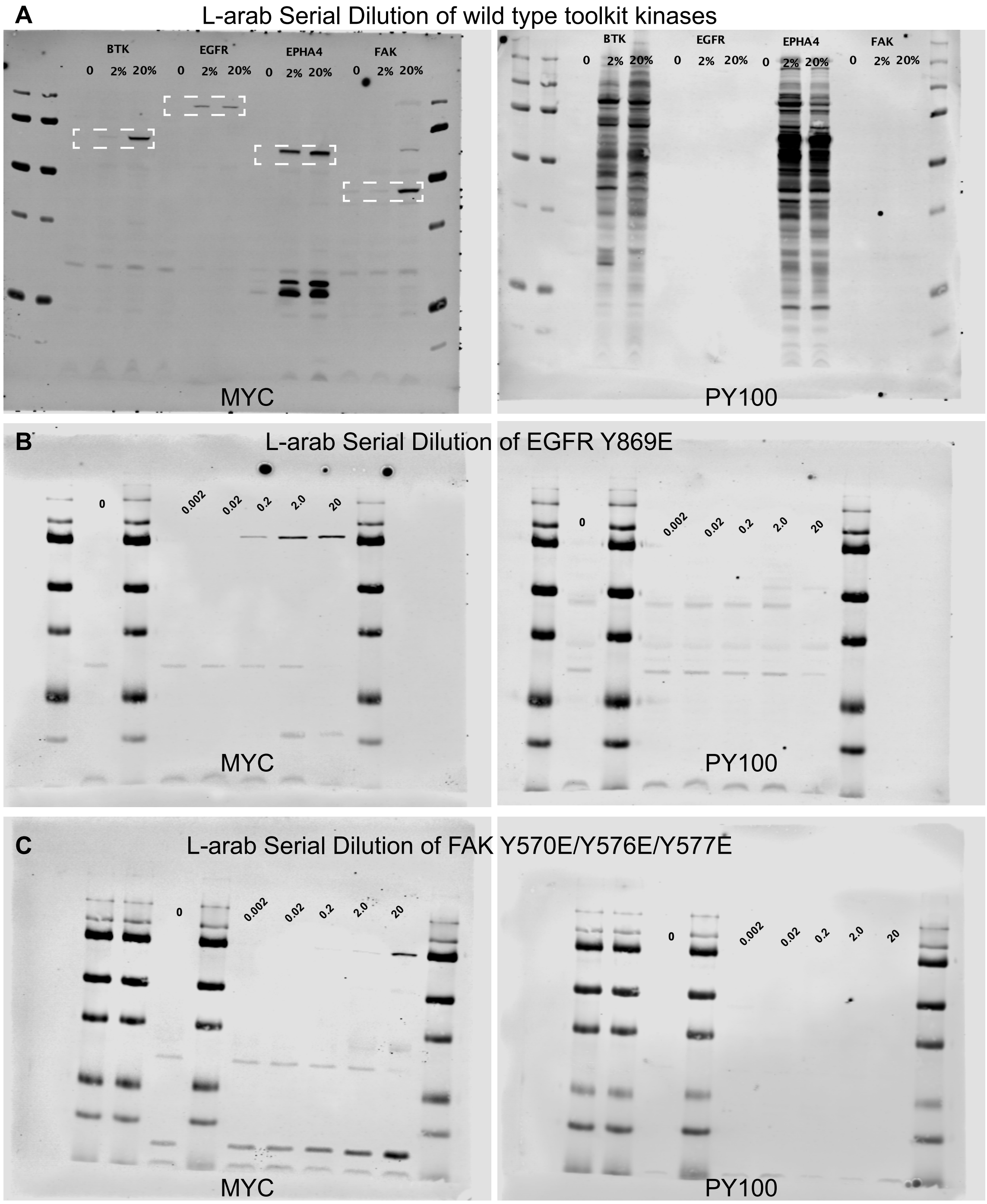


**Supplementary Figure S1. Kinase expression and activity testing. A)** Example of testing L-arabinose serial dilution and the effect of expression of kinase (MYC) and activity (PY100). Activity can be seen at very low MYC-based detection of kinase expression for BTK and EPHA4. Despite expression of EGFR and FAK, no activity of the kinase was seen. **B)** Mutagenizing the EGFR Y869 activation loop site to E is insufficient to drive activity. **C)** Similarly, the triple mutant of the FAK Y570/Y57/Y577 to E mutation is insufficient to drive activity.

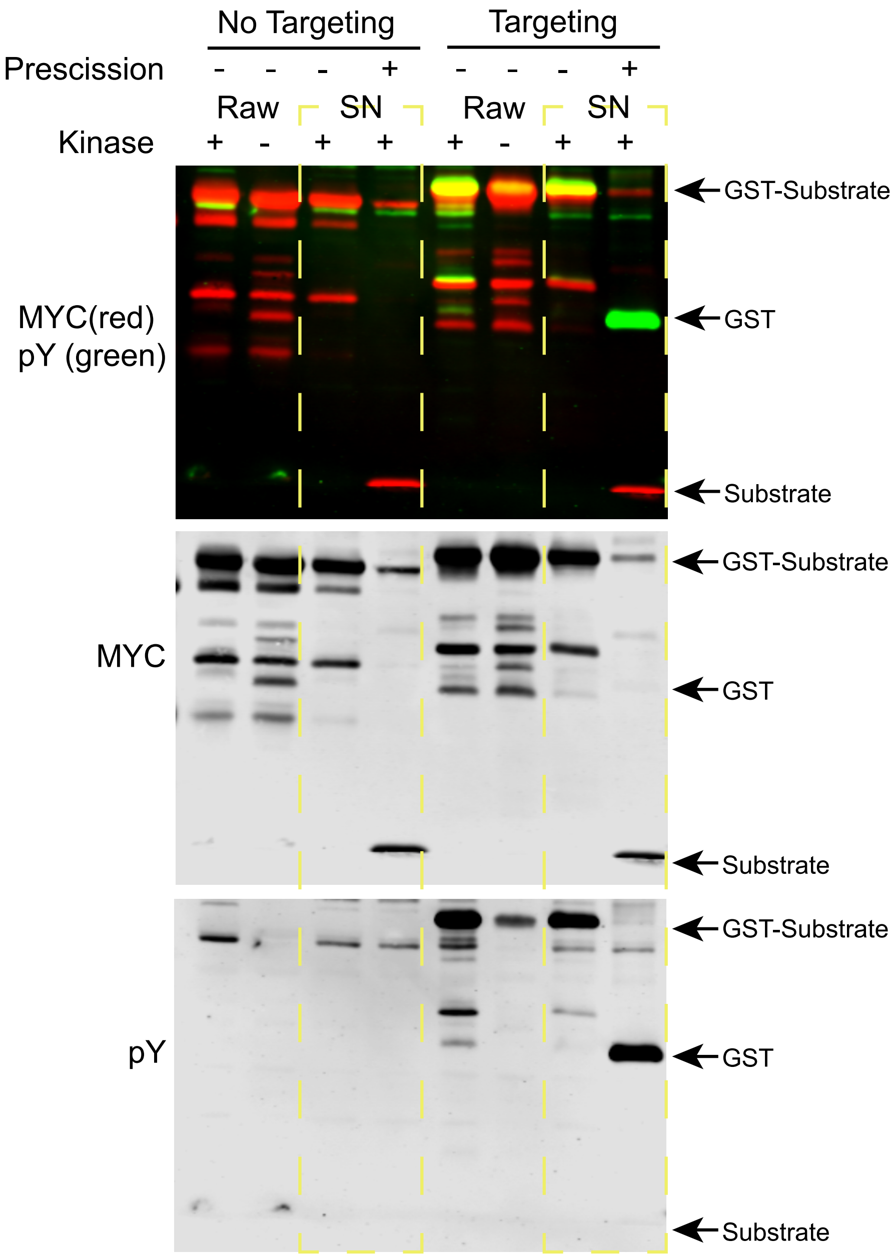


**Supplementary Figure S2. GST sequences is excessively phosphorylated under targeting conditions.** Here, co-expression of ABL kinase with a substrate on the original pGEX backbone (a GST fusion) shows no phosphorylation by anti-pY antibody in the absence of targeting (p40 sequence). This is boosted upon targeting, but using PreScission to separate the GST and p40-sequence in supernatant (SN) treatment shows that the majority of phosphorylation occurs on the GST sequence.

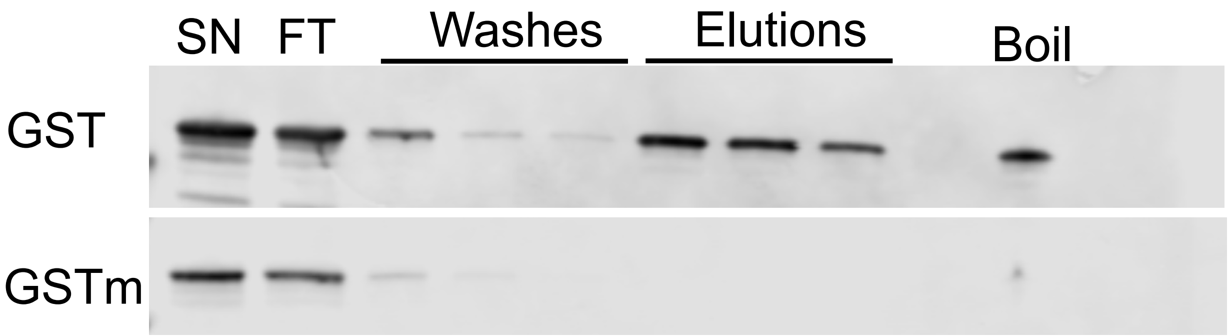


**Supplementary Figure S3. A mutant GST sequence (GSTm) fails to bind GST resin for purification.** Given the high degree of phosphorylation of GST sequence (Fig. S2), we synthesized a GSTm sequence, with all tyrosines mutated to phenylalanines. Here we show an attempt (of several that reproduced) where GST agarose does not capture GSTm sequence. The GRP1 PH domain is the fusion protein to the GST- or GSTm- sequences. No protein is found in either elutions or boil of the GSTm construct.

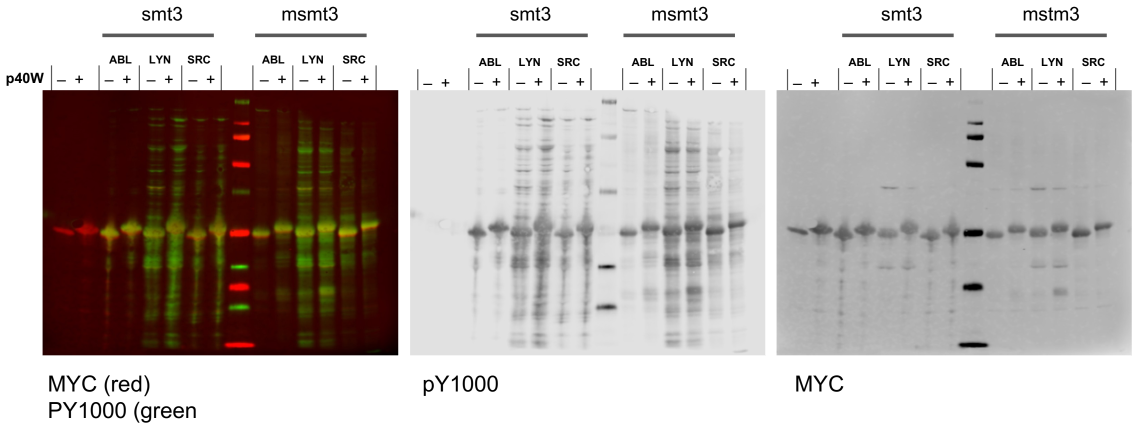


**Supplementary Figure S4. Testing Y67F in the smt3 domain on production of phosphoprotein.** We wished to eliminate non-targeting sequence tyrosines in the substrate fusion proteins, which includes a single tyrosine in the smt3 (yeast SUMO domain added for increased solubility). p40W indicates tyrosine in the p40 mutated to W. Here, we test production of EGFRCtail fused to smt3 or msmt3, with and without p40W targeting sequence and co-expressed with either kVh-ABL, kVh-LYN, or kVh-SRC (supernatants). Mutated smt3 sequence has no discernible effects on either production or phosphorylation of the substrate. All detected phosphorylation on the substrate protein in these constructs are of tyrosines in the EGFRCtail.

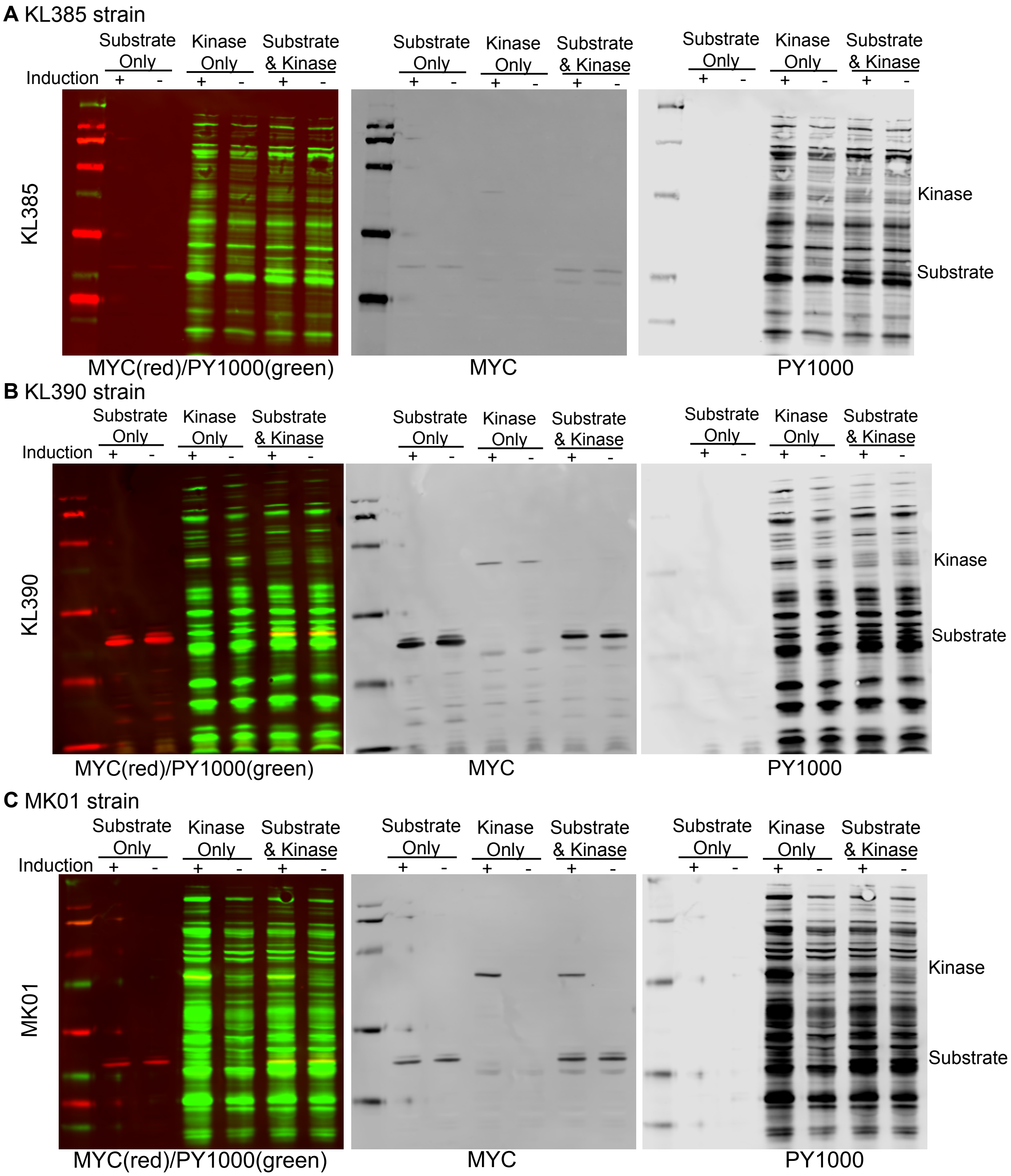


**Supplementary Figure S5. Testing E. coli strains engineered for dual induction of L-arabinose and galactose-based induction.** Strain production was tested using an SH2 domain substrate (LYN SH2 domain) with kVh-LYN kinase. We tested induction of substrate (+IPTG) or kinase (+L-arabinose) as single transformants and then as dual transformants. All substrate strains are in the presence of IPTG and Induction refers to the addition of L-arabinose (e.g. testing whether L-arab addition alone changes the ability to produce substrate in a single transformant or a dual transformant). A) KL385 strain shows poor induction of both substrate and kinase (as detected by MYC, though high kinase activity by PY1000 staining). B) KL390 strain shows improved substrate production, compared to KL385, but co-transformation reduces both kinase and substrate expression and leaky expression of the kinase. C) MK01 strain shows the highest kinase expression of engineered strains tested, but fairly poor production of the substrate.

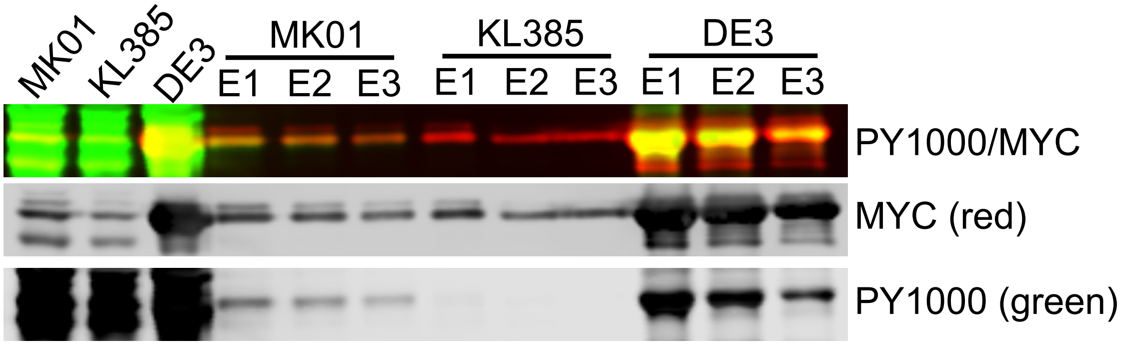


**Supplementary Figure S6. Expression and purification of phosphosubstrate from three E. coli strains.** Supernatants (input) on the left for MK01, KL385, and DE3 as a direct test of phosphoprotein production, where three elution volumes post-nickel purification from each strain are shown on the remaining lanes. We concluded that the DE3 strain, though not engineered explicitly for L-arabinose based expression, produces the highest phosphoprotein, since it produces the highest overall substrate protein with sufficient expression of the kinase.

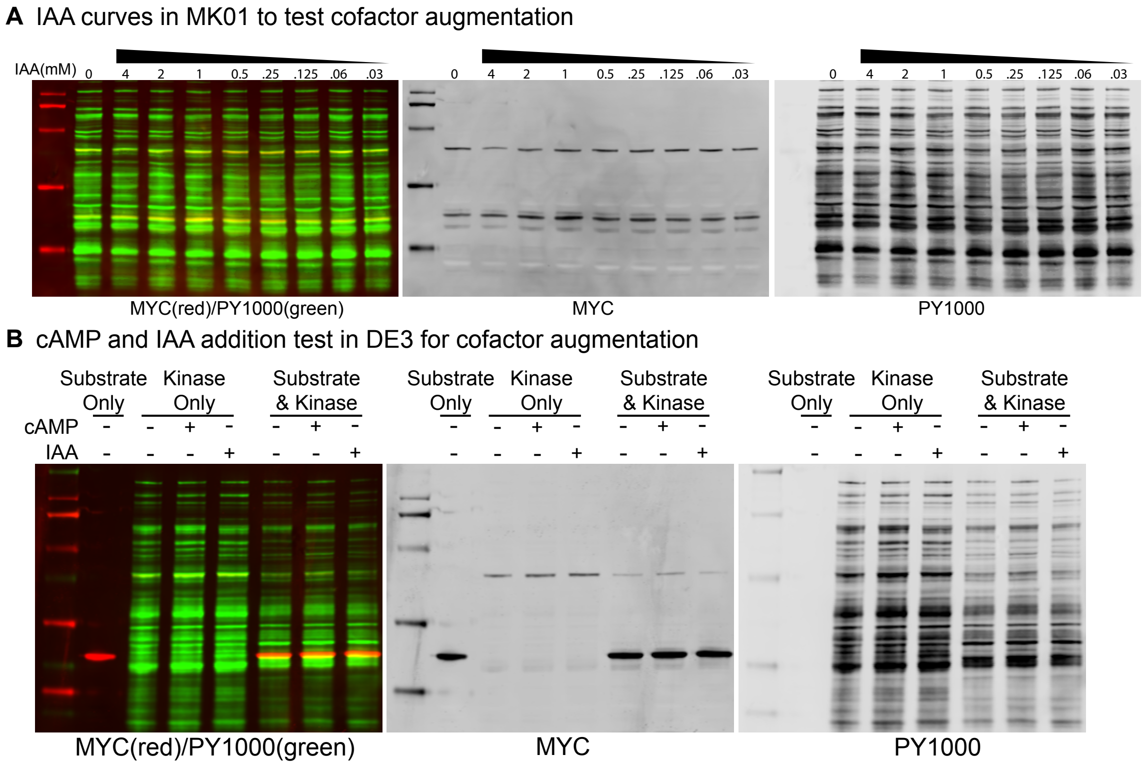


**Supplementary Figure S7. Testing if central metabolism alterations can increase co-expression of kinase and substrate.** A) We performed a serial dilution of indole-3-acetic acid (IAA) in the MK01 strain of a substrate (an SH2 domain) co-transformed with the kVh-LYN kinase and co-induced with 0.5mM IPTG and 0.2% L-arabinose, to test for improvement in expression of kinase or substrate with the addition of IAA. We saw marginal increases around 1mM of IAA. B) An example co-factor addition test, in DE3 cells, of cAMP or IAA to test for the increase in expression of kinase and substrate. Each of the transformations have been induced with IPTG (0.5mM) or L-arabinose (0.2%) in single or co-transformations of an SH2 domain substrate and the kVh-LYN kinase. cAMP or IAA were added at the time of induction at 1mM.

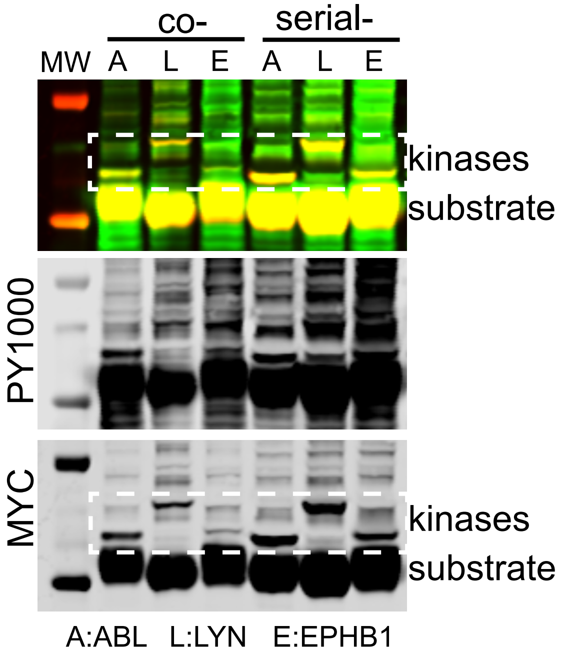


**Supplementary Figure S8. Evaluation of in vitro, co-expression, and serial inductions on phosphoprotein yields**. We directly compared the EGFRCtail (p40) phopshoprotein production by in vitro reactions, co-expression, or serial induction with three kinases (ABL, LYN, and EPHB1). Here, the supernatant expression of co-expression and serial inductions show kinase expression increases in serial induction as seen by MYC-based detection. Purified phosphoprotein is shown in main Figure 1E.

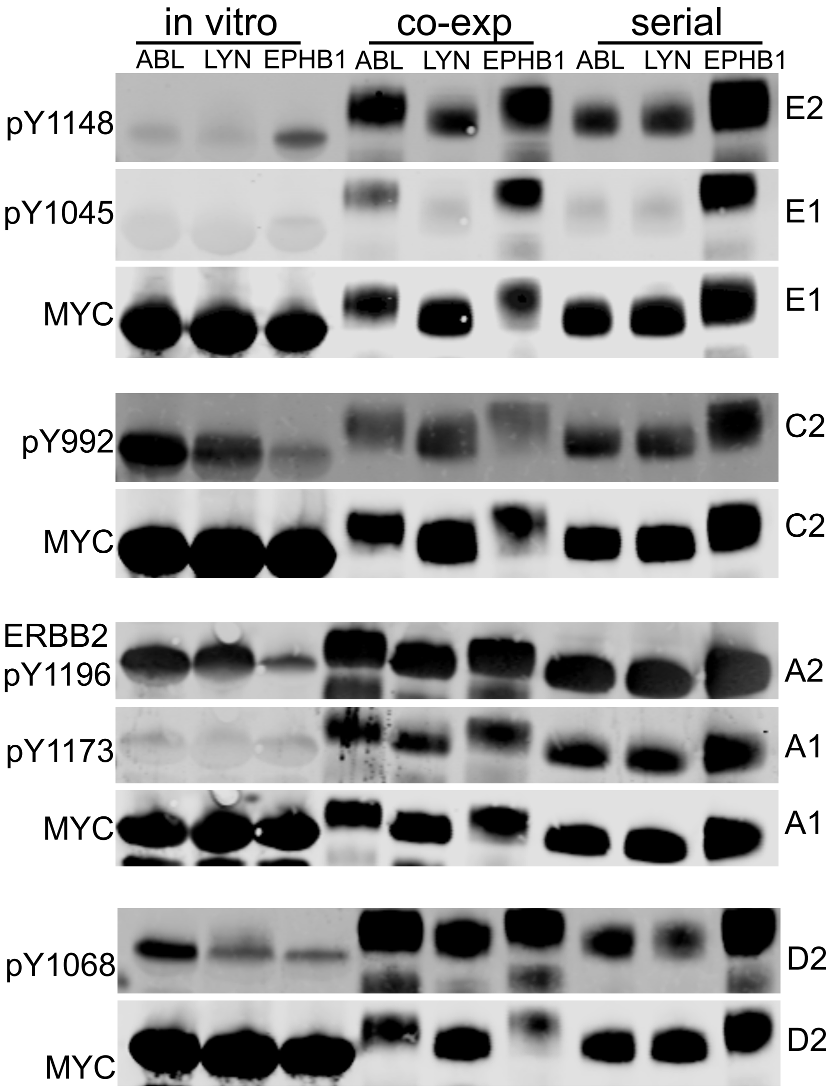


**Supplementary Figure S9. Evaluation of in vitro, co-expression, and serial inductions on specific phosphorylation differences using EGFR phosphospecific antibodies**. EGFR Ctail from in vitro, co-expression, and serial induction reactions with ABL, using the p40 fusion, were probed with EGFR and an ERBB2 phosphospecific antibodies to measure the relative differences in phosphorylation yields detected by that antibody. The ERBB2 pY1196 antibody was used as a possible detection of the homologous EGFR pY1138 site. The ABL kinase here was cloned from the Albanese 2018 library. The letter numbers indicate the individual blots (replicates A-E) and the probe number. For example, A2 indicates blot A was stripped and reprobed with the antibody given. We limited reprobes to one time.

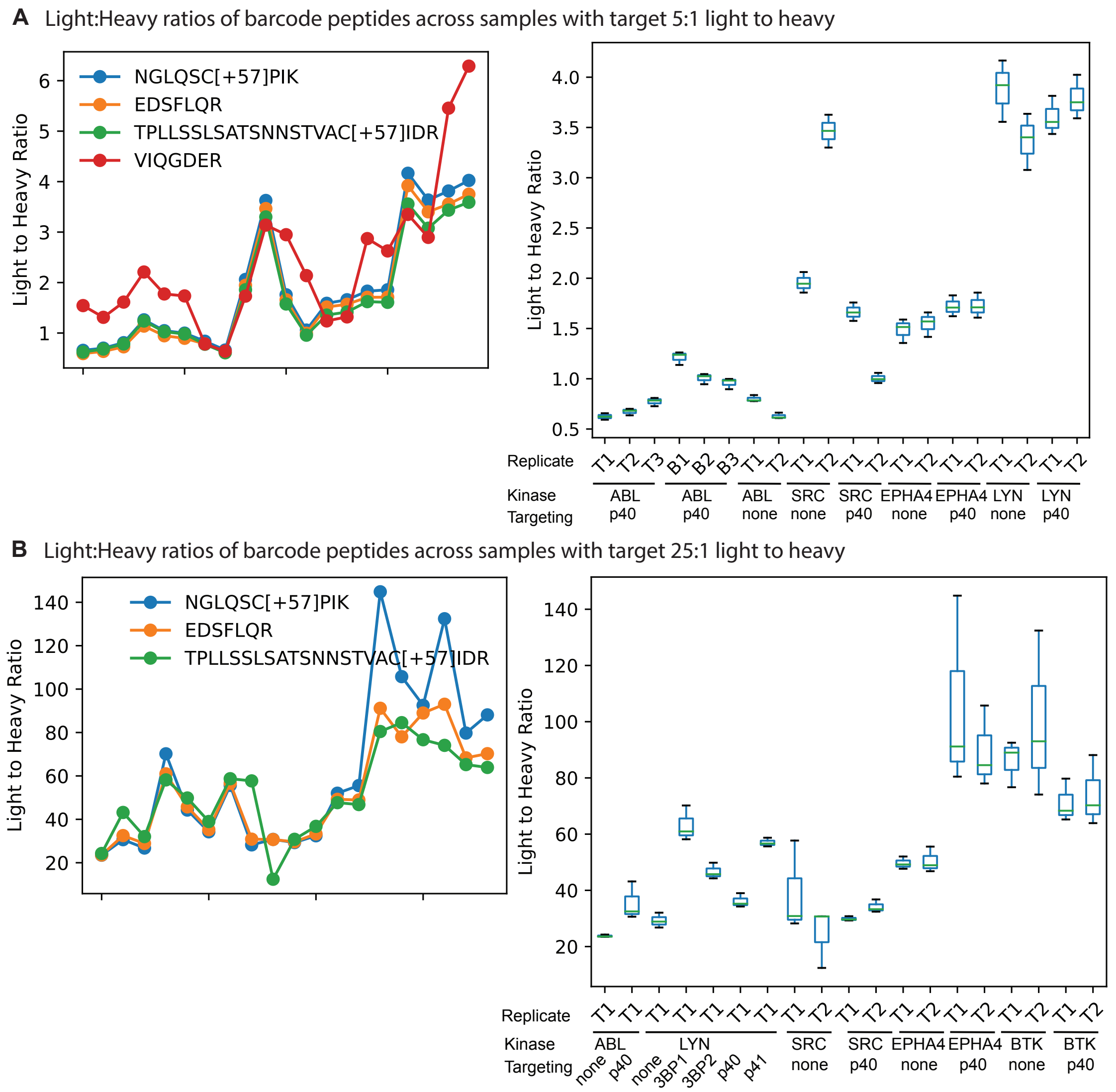


**Supplementary Figure S10. Testing the Light to Heavy ratios of non-phosphorylatable "barcode" peptides in the EGFRCtail across two sets.** A) Twenty samples were analyzed by LC-MS/MS where we included an admixture of 10ug of light phosphoprotein with 2ug of heavy non-phosphorylated standard (targeting a 5:1 ratio). Left: Four tryptic fragments were observed across all samples that cover EGFRCtail peptides that do not contain a phosphorylatable tyrosine and we refer to these as "barcode" peptides for possible normalization approaches of tryptic fragments with tyrosines. We find that the first three peptides produce very high replication of the relative ratio of light to heavy, having Pearson's correlation of more than 0.997. Right: Using the first three tryptic fragments, this shows the light to heavy ratios of each of the samples using each of the three barcodes. Technical replicates were run (T) and for one set (p40-targeted EGFR Ctail reactions with ABL), we performed a biological replicate (B). There is fairly good correlation between the estimated ratios of the light to heavy samples within replicates and all samples suggest systematically lower ratios than the target of 5:1 that was expected. B) Twenty-four samples, this time with 5ug of light phosphoprotein and 0.2ug of SILAC standard (targeting a 25:1 ratio) were analyzed and the same three bar codes used in the first set are shown here for their overall concordance and range produced for each of the samples.

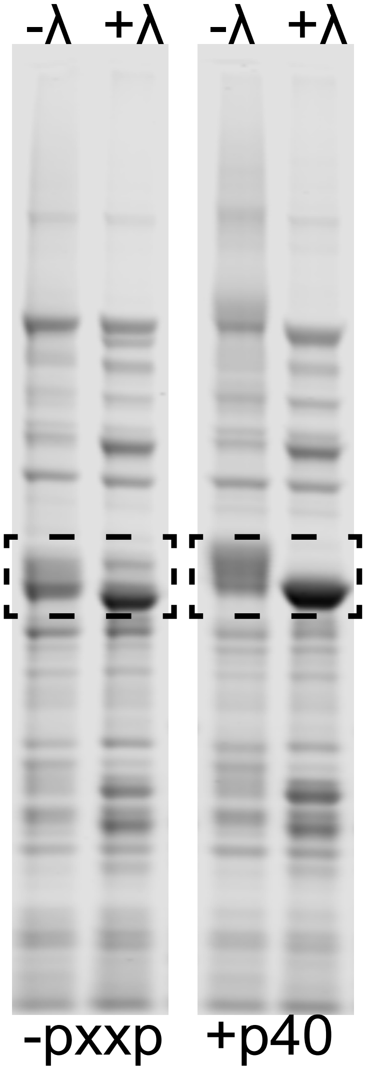


**Supplementary Figure S11. Lambda phosphatase treatment improves Coomassie-based protein estimations.** Example of protein estimation issues caused by high degrees of phosphorylation. Here, the EGFRCtail was co-expressed with kVh-ABL kinase, in targeting (p40) or non-targeting (pxxp) reactions and purified using nickel-based purification. This Coomassie shows the collapse of high phosphospecies into a centralized band when treated with lambda-phosphatase. We found that substrates containing more than two phosphotyrosine sites cause estimation issues in protein concentration by Coomassie due to spread of the band, but also likely dye binding changes that are affected by the phosphorylation.

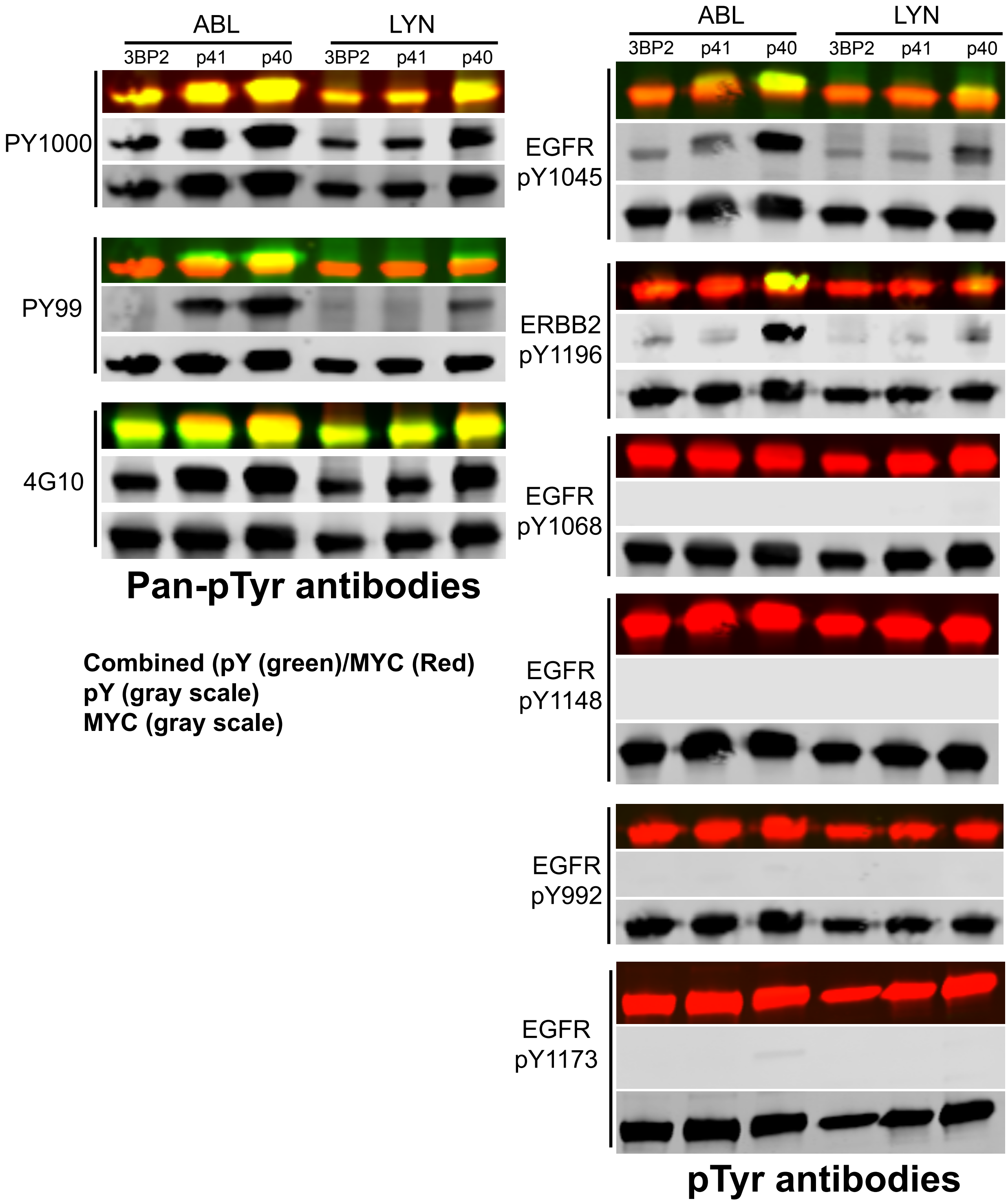


**Supplementary Figure S12. 1069F control used to test antibody detection of PxxP tyrosines.** Using the 1069 21-mer sequence, we mutated the 1069 site to a phenylalanine, leaving only the tyrosine contained in the polyproline sequence. We then co-expressed this with ABL or LYN kinase and detected phsophorylation on all tyorsine containing PxxP sequences (3BP1, p41, and p40) by pan-specific pTyr antibodies. Using these as controls, we then tested the EGFR family phosphospecific antibodies to be tested on 21-mer sequences. We found that three of the five antibodies did not detect PxxP pTyr (PY1068, pY1138, and pY992), suggesting these antibodies can be used without requiring targeting sequence removal. However, EGFR pY1045 and ERBB2 pY1196 did detect at least one targeting sequence (including p40, the high affinity site). Some westerns were stripped and reprobed up to one time, but all probes included combined MYC and pTyr antibodies in dual channels, so we have included the accompanying MYC control signal for each blot.

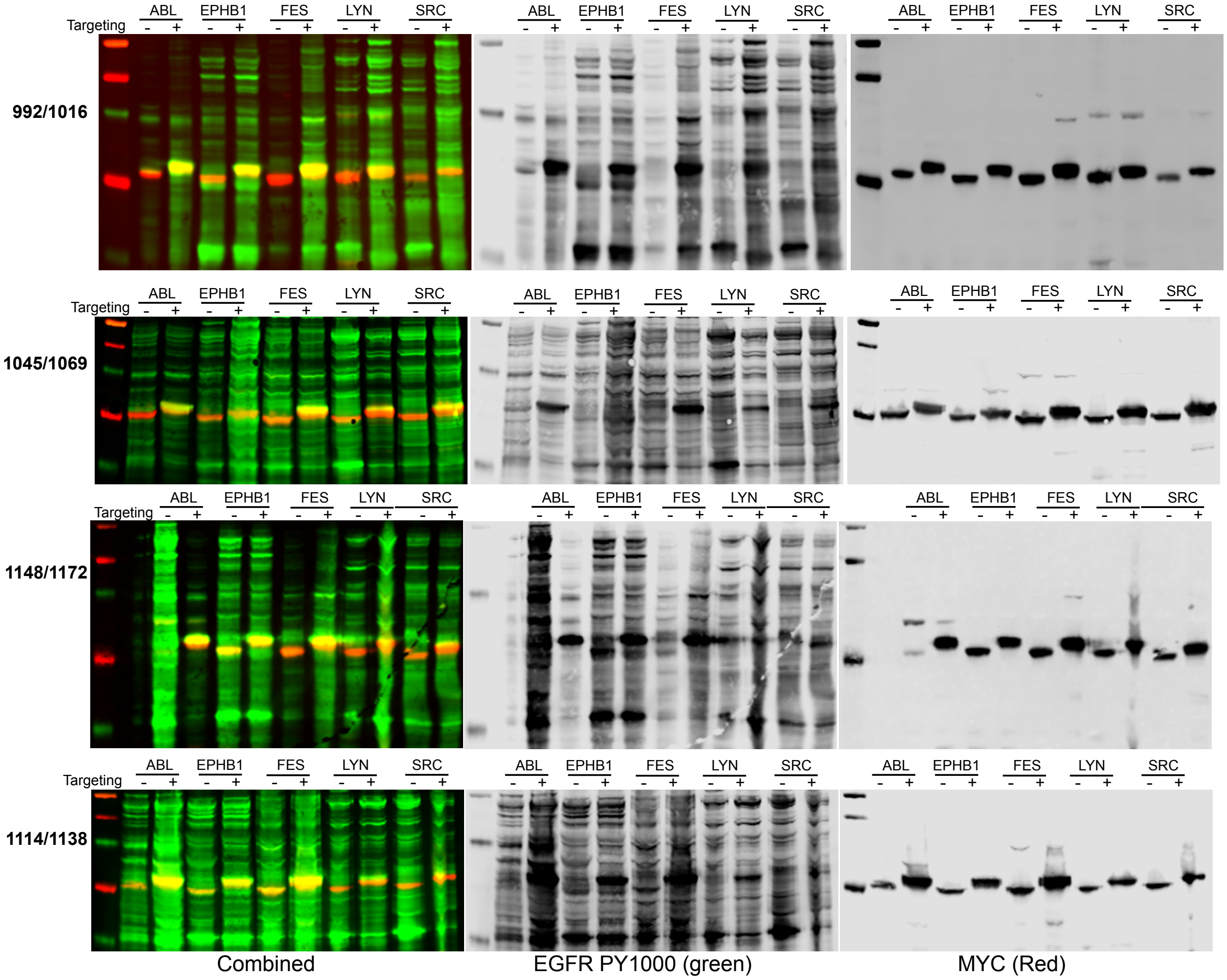


**Supplementary Figure S13. Supernatant probes of the 21-mer co-expressions.** Supernatants of the 21-mer coexpression experiment, testing five kinases against specific EGFR tyrosine 21-mer peptides are shown here, probed with MYC(red) and PY1000(green) to test the general activity and expression of kinases and substrate expression.

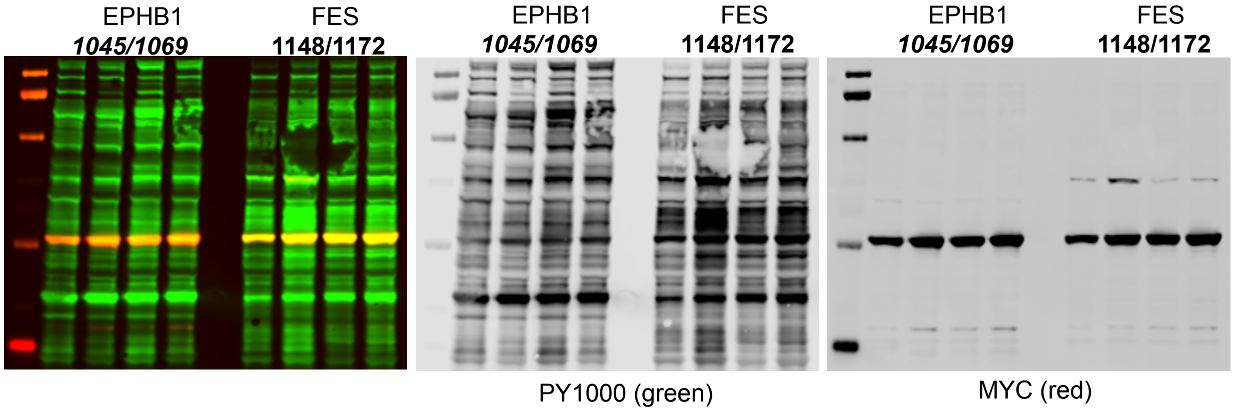


**Supplementary Figure S14. Testing variability of kinase and substrate expression for two 21-mers.** Fig. S13 we saw low kinase expression in the no PxxP 21-mer experiments of Y1069 with EPHB1 and Y1148 with FES. Here, we repeated expression on the glycerol stock of the co-transformant (left lanes) and selected three additional clones of the co-transformants and replicated co-induction. We observed each of these produced high expression of kinases and substrate, suggesting the variability observed was not due ot the inherent clone, but to the induction process.

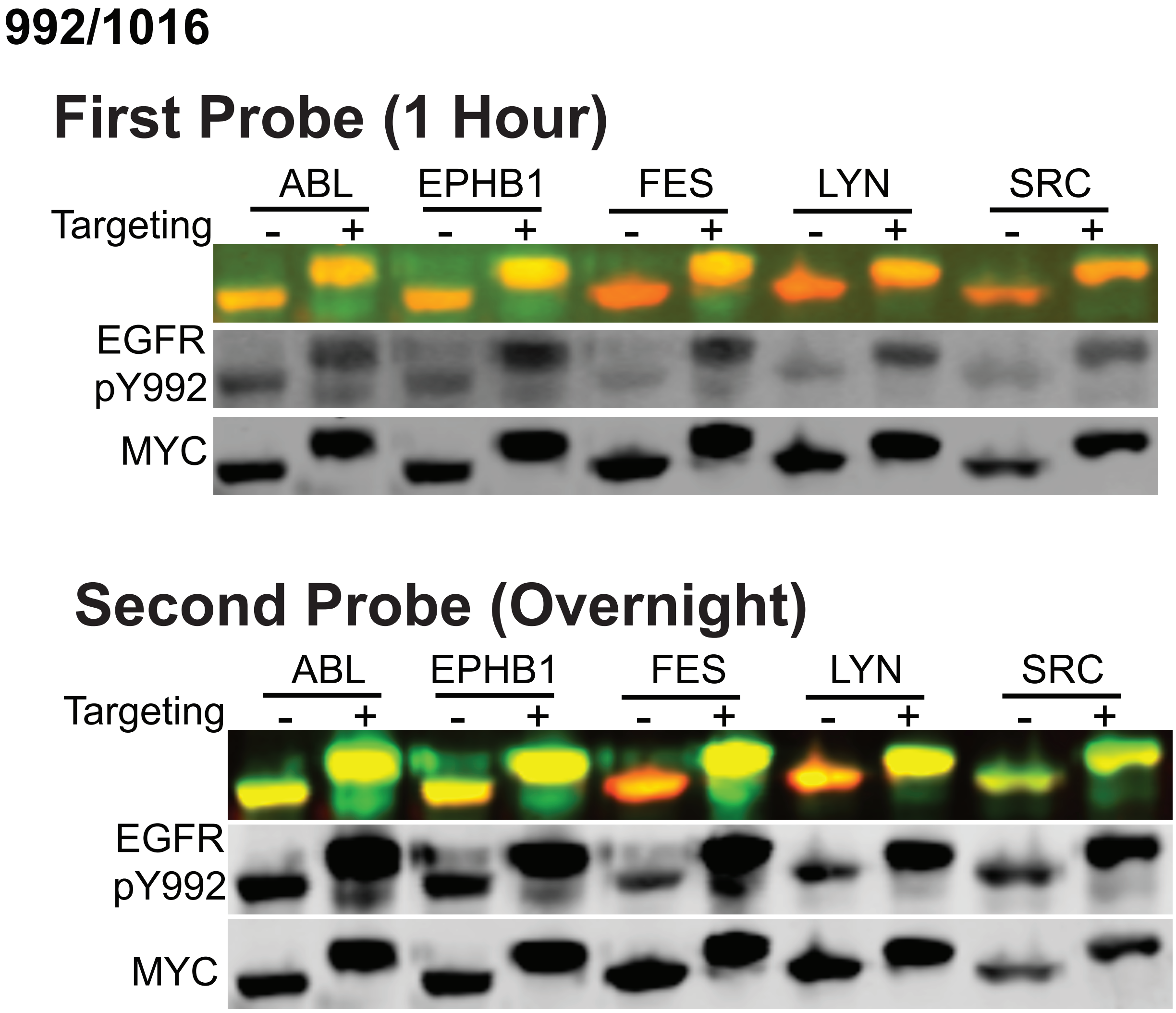


**Supplementary Figure S15. Low SNR with control highlights antibody issue.** In an initial EGFR pY992 probe on 21-mers, we saw low signal to noise. We extended the incubation of the primary antibody to overnight and reduced dilution from 1:3000 to 1:1000 and significantly improved signal. Both probes were on the same membrane (second probe after a stripping).

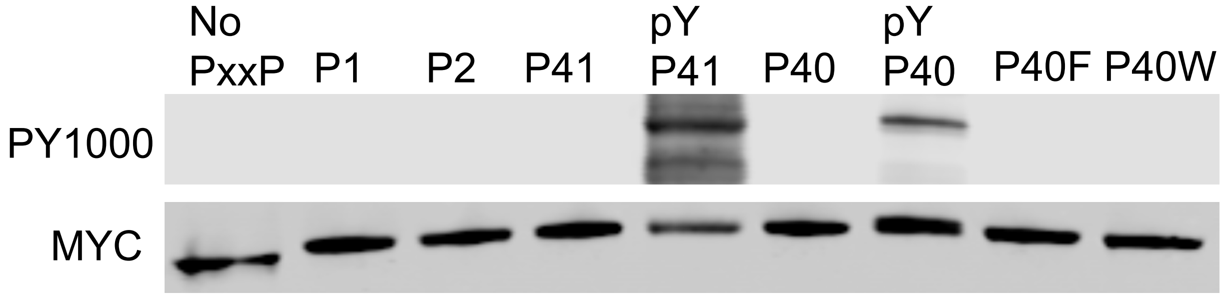


**Supplementary Figure S16. Load control western shown here for SH3 far western experiment.** Lysates containing 21-mer 1069F sequences were loaded to be approximately equal by initial western analysis. This western was run on the same samples and same concentrations as the SH3 far western and was probed with MYC (600 channel) and then reprobed by PY1000 antibody (800 channel) to confirm phosphorylation of the p41 and p40 samples.
